# *NicheDiv*: A DAPC framework to quantify niche divergence across highly multivariate environmental space

**DOI:** 10.64898/2026.06.19.733388

**Authors:** Daniel Schönberger, Zachary G. MacDonald, B. Christian Schmidt, Julian R. Dupuis

## Abstract

Quantifying niche divergence is crucial to understanding the ecological and evolutionary processes underlying range limits, coexistence, speciation, biogeography, and macroevolution. Yet available approaches rely on low-dimensional climate summaries, are vulnerable to multiple biases, or struggle with high-dimensional collinear data. We introduce the *R* package *NicheDiv*, which adapts discriminant analysis of principal components (DAPC) to quantify pairwise niche divergence across any number of abiotic and biotic environmental variables associated with occurrence records. Our method first addresses correlations among environmental variables through principal component analysis. It then identifies a single discriminant axis that maximizes separation between predefined groups (species/lineages/populations), summarizing multivariate niche structure into one dimension. Significance is assessed by a permutation test that reshuffles group identities to mimic a shared niche. To characterize ecological differentiation, *NicheDiv* calculates Schoener’s D as an overlap index and extends the niche divergence plane to multivariate space, providing metrics such as niche dissimilarity and exclusivity. Extracted variable contributions from the discriminant axis identify environmental variables that contribute most to divergence. Using simulations and empirical data together with a large set of environmental layers, we demonstrate that *NicheDiv* is computationally scalable, detects subtle divergence in high-dimensional space despite multicollinearity, distinguishes different forms of niche divergence (weighted, nested, soft, hard), and identifies the variables that potentially drive divergence. Compared with alternative divergence tests (PCA-env, hypervolumes, MVNH, PERMANOVA, PCA-space, and logistic regression), *NicheDiv* generally retains more variation, scales more consistently with increasing divergence, and returns more interpretable effect sizes. *NicheDiv* automatically extracts such environmental data from preconfigured and user-supplied GIS layers and implements a preprocessing pipeline that reduces known biases: delimiting accessible background space, spatially thinning occurrences, balancing sample sizes, filtering low-information variables, and screening predictors for between-group environmental analogy. We test our framework with empirical analyses of *Hemileuca* buck moths and demonstrate that their niches are structured by a range of seasonal abiotic and biotic variables rather than annual climatic averages. Overall, *NicheDiv* offers a robust framework for characterizing niche divergence across multiple environmental axes in support of species delimitation, local adaptation, community ecology, biogeography, and macroevolution.

## INTRODUCTION

Assessments of niche divergence are essential for understanding patterns of biodiversity and the processes that generate them. For example, testing for niche divergence helps clarify species’ range limits, biogeography, and community structure (Glor & Warren, 2011; Soberón, 2007). It reveals patterns of coexistence among species that appear ecologically similar through resource partitioning and habitat specialization (Leibold & McPeek, 2006; Schoener, 1968). By linking ecological differentiation to reproductive isolation and lineage diversification, niche divergence tests mechanistically link ecological processes, such as habitat filtering and environmental specialization (Blair et al., 2013; McCormack et al., 2009), to the broader patterns of speciation and macroevolution (Castro-Insua et al., 2018; Kozak & Wiens, 2010). It also provides evidence independent of morphology and genetics for evaluating species limits and identifying cryptic taxa (MacDonald et al., 2025; Raxworthy et al., 2007). Finally, such approaches enable quantitative insight into niche shifts during biological invasions and range expansions (Guisan et al., 2014; Liu et al., 2020). These applications require efficient methods that accurately characterize niche divergence.

The ecological niche is often defined as an *n*-dimensional hypervolume encompassing biotic and abiotic dimensions (Hutchinson, 1957). However, niche divergence studies have predominantly focused on climatic, annually averaged variables, such as *WorldClim*’s BIO1–BIO19 variables (Hijmans et al., 2005). Yet capturing the full niche of a species requires axes extending beyond climate to include other abiotic and biotic variables (Elith & Leathwick, 2009; Kearney & Porter, 2009; Soberón, 2007). Annual summaries may obscure key ecological processes while seasonal variables capture phenological and seasonal dynamics that drive resource availability, physiological stress, and interspecific interactions (Prajzlerová et al., 2025; Zimmermann et al., 2009). A shift from annual climatic variables toward seasonal and multi-axis treatment of abiotic and biotic variables is now feasible, given the growing availability of high-resolution GIS datasets on climate, vegetation, soils, terrain, and urbanization (Batjes et al., 2024; Hufkens et al., 2018; Potapov et al., 2021).

As GIS layers become more available, datasets become more multivariate and collinear, complicating model parameterization and interpretation, thus posing challenges for existing niche divergence methods. For instance, PERMANOVA, a distance-based multivariate method, emphasizes centroid separation but is sensitive to differences in within-group dispersion (Anderson, 2001). PCA-based comparisons, such as *PCA-env* (Broennimann et al., 2012), are unsupervised, maximize total rather than between-group variance, and focus only on the first two axes of variation. Hypervolume methods approximate multivariate niches but face key drawbacks: kernel density estimates (Blonder et al., 2014) capture complex, multimodal niches but lose interpretability and robustness as dimensionality increases, while multivariate normal hypervolumes (Lu et al., 2021) are computationally efficient but rely on covariance estimates that become unstable under strong collinearity. Correlative environmental niche models paired with identity or background-similarity tests (Warren et al., 2010) are flexible but can make it difficult to distinguish true niche differences from differences in the environmental conditions available within each species’ accessible area, and test statistics depend on model tuning and thresholding (Broennimann et al., 2012; Brown & Carnaval, 2019). Although the niche divergence plane framework (Ascanio et al., 2024) allows differentiation among divergence types, its application is restricted to single variables. These constraints motivate a new method that handles multivariate collinear datasets while returning interpretable metrics of divergence and variable contributions.

Niche divergence testing is further complicated by several practical and conceptual pitfalls. First, spatial autocorrelation and uneven sampling effort can inflate separation (Dormann et al., 2013). Second, comparisons in geographic space via correlative niche models can conflate true niche similarity with uneven environmental availability (Soberón, 2007; Warren et al., 2010). This can bias overlap estimates toward prevalent habitats and confound niche overlap with differences in environmental availability between groups (Brown & Carnaval, 2019). Third, differences in accessible areas can create artefactual divergence even when species share similar niches, so null models must disentangle abiotic and biotic environmental availability from the processes shaping species’ distributions (Brown & Carnaval, 2019). Furthermore, groups may appear differentiated because they occupy non-overlapping environments rather than because their realized niches diverge, so comparisons should be restricted to shared, analogous environmental conditions (Brown & Carnaval, 2019). Lastly, non-equilibrium distributions are a major bias, as species’ ranges are often assumed to reflect equilibrium with environmental conditions, yet represent transient snapshots shaped by historical lags of species’ range responses to past environmental change, competitive exclusion, and short-term seasonal fluctuations (Brown & Carnaval, 2019). Consequently, present occupancy can under-represent the set of tolerable conditions, and inferring niche limits or divergence directly from ranges is risky (Araújo & Pearson, 2005; Peterson et al., 2011). To mitigate these biases, tests should reduce spatial autocorrelation, match accessible areas in extent and environmental availability, compare taxa in shared environmental space, restrict to analogous environmental conditions, and summarize divergence on a common axis using seasonal predictors.

We present a novel application of discriminant analysis of principal components (DAPC; Jombart et al., 2010) for pairwise tests of niche divergence. As input, the method uses any number of abiotic and biotic ecological variables (hereafter “environmental variables”) derived from occurrence coordinates of two predefined groups. First, variables are transformed into uncorrelated axes (principal components or PCs) via principal component analysis (PCA). Using the axes (PCs) that capture most environmental variance, discriminant analysis then maximizes between-group separation while minimizing within-group variation. This yields a single discriminant axis that summarizes the multivariate niche structure between both groups in one dimension, allowing direct statistical and visual interpretation. Using empirical and simulated occurrences combined with an extensive environmental dataset, we assess whether DAPC can detect divergence in high-dimensional space, distinguish niche divergence types, and detect the variables contributing most to separation. We compare our approach against six alternative niche divergence tests. Finally, we provide a flexible function for extracting multiple environmental layers and a standardized preprocessing pipeline for reducing known biases. Our *NicheDiv* framework (Figure 1) is implemented in *R* (R Core Team, 2025) and includes a step-by-step tutorial (Github-link-to-be-inserted).

**Figure 1.**
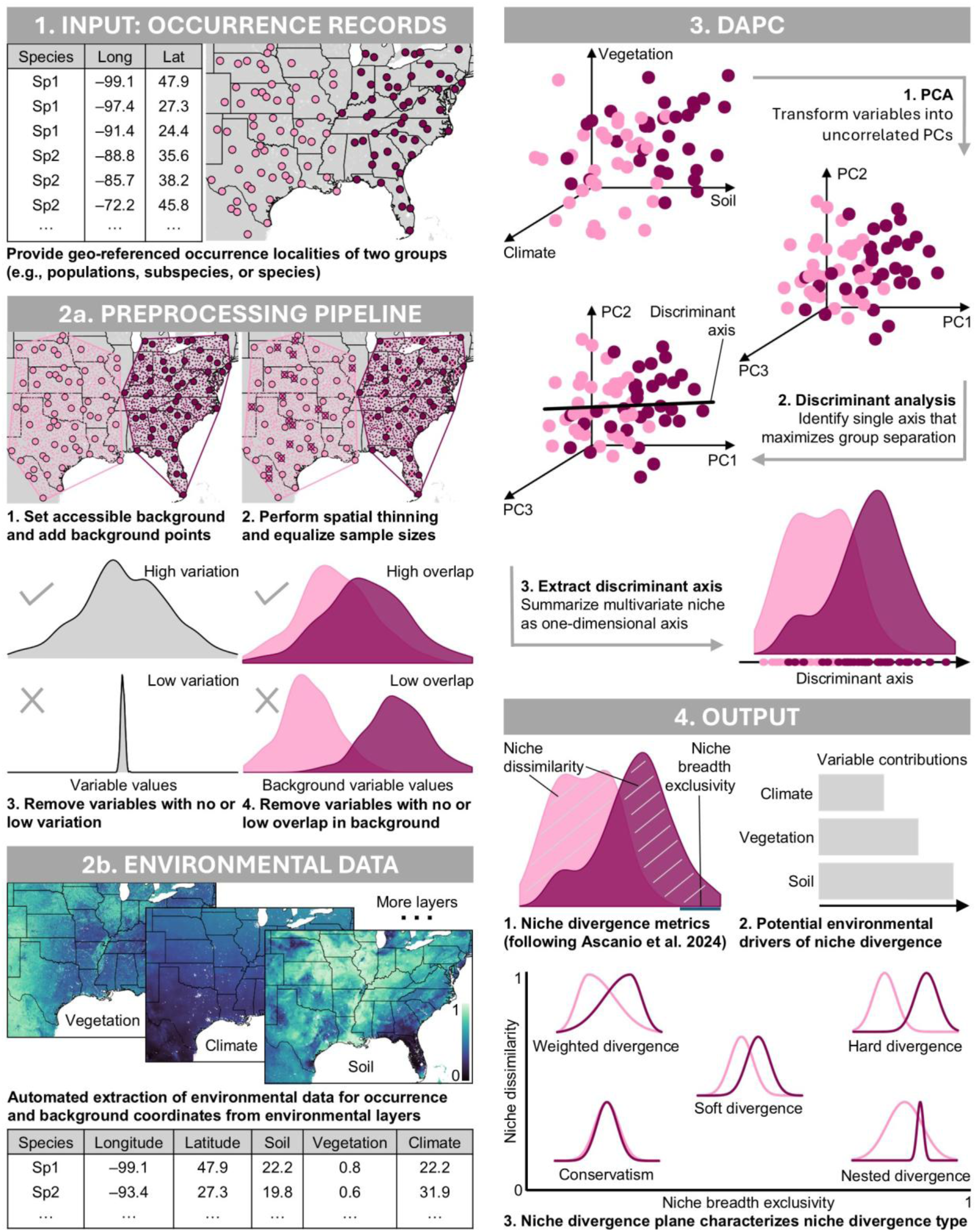
Overview of DAPC niche divergence framework in *NicheDiv* R package. The summary is based on two theoretical species pairs and three environmental layers. The framework can incorporate many environmental GIS layers, including user-specified ones.

## METHODS

### DAPC

Jombart et al. (2010) developed DAPC to identify genetic clusters from large sets of genetic markers. We adapt DAPC to environmental data associated with geo-referenced occurrence records to test for niche divergence between two *a priori* groups (e.g., species, lineages, populations) (Figure 1). Like genetic markers, environmental variables are often numerous and correlated, making DAPC well-suited for this application.

The core of DAPC is discriminant analysis, a supervised classification method for detecting subtle differences between groups in multivariate space (Lachenbruch & Goldstein, 1979). Rather than maximizing overall variance like PCA, discriminant analysis maximizes between-group variation while minimizing within-group variability. This is done by identifying synthetic variables (“discriminant functions”), which are linear combinations of original variables and define discriminant axes that maximize group separation. Samples are assigned probabilistically to each group based on their position along these discriminant axes. However, discriminant analysis alone is unsuitable for most genetic and environmental datasets due to model instability under multicollinearity and high dimensionality (Lachenbruch & Goldstein, 1979). DAPC resolves this statistical issue by performing a PCA prior to discriminant analysis. This transforms the original variables into uncorrelated orthogonal axes (PCs) that summarize most variation, mitigating model instability due to multicollinearity and high dimensionality. Discriminant analysis is then performed on the retained PCs.

A challenge in DAPC is determining how many PCs to retain for the discriminant analysis: retaining too few results in information loss and poor group separation, while too many lead to overfitting and inflated predictive accuracy. To identify the optimal number of PCs, our method performs two-stage cross-validation. In each replicate, occurrences within each group are randomly partitioned into 90% training and 10% validation subsets, and the corresponding PC scores are used for model fitting and evaluation. A discriminant analysis is fitted to the training set, and predictive accuracy is evaluated on the validation set. Cross-validation is performed in two stages to reduce runtime. Stage 1 searches up to the maximum possible PCs in steps of 10, and stage 2 refines the search within a 30-PC window around the best stage 1 candidate. The optimal number of PCs is the value minimizing the cross-validated root mean squared error of group assignment, representing the balance point between underfitting (loss of discriminatory signal) and overfitting (noise captured as signal), where additional PCs no longer improve predictive accuracy. After cross-validation, the main discriminant analysis is fitted with the determined PC number. Two metrics are calculated for model fit: mean assignment accuracy (reflects the distinctiveness of groupings) and the adjusted Rand index (quantifies how well DAPC-predicted groupings match true taxon labels while accounting for chance).

To evaluate the significance of observed niche divergence, our method implements a permutation procedure that constructs a Null expectation of group separation, emulating a shared niche (Supplementary Methods M3). Group labels are randomly reassigned across individuals while preserving group sample sizes, and the discriminant analysis is re-run. Repeating this process yields a Null distribution of assignment accuracy, allowing evaluation of whether niche separation between taxa exceeds chance expectations. If divergence is non-significant, an optional test equivalent to the Background statistic of Brown & Carnaval (2019) can be performed. This test evaluates whether an observed lack of divergence reflects niche equivalence or limited power to detect divergence due to similar environmental conditions occupied by both taxa within their accessible areas. This is done by replacing one taxon’s occurrences with random samples from the other’s background (and vice versa) to generate a Null distribution conditioned on each taxon’s available environment. Assuming a non-significant permutation test, a significant background test indicates niche similarity, whereas a non-significant result suggests limited power to detect niche differences given the environment available to each taxon.

The DAPC method is implemented in *run.DAPC.crossval.permutation*, which handles data processing, cross-validation, DAPC fitting, and permutation testing. Its input is a matrix or data frame containing environmental variables (rows = occurrences, columns = variables) and a two-level column of group assignments (Figure 1). Because DAPC cannot accommodate missing values, occurrences exceeding a user-defined missing-data threshold are removed. Remaining missing values can either be imputed using the median of each variable or handled by removing incomplete cases.

### Niche divergence metrics and visualization

Generating a univariate discriminant axis allows multivariate niche space to be summarized in a single dimension, facilitating the interpretation of niche divergence and identifying its different types, such as nested or partially exclusive niches. Our package provides tools to evaluate and visualize these outputs (Supplementary Methods M4).

Permutation results are shown as a null distribution of mean assignment accuracy, with the empirical value plotted for comparison. Niche overlap is visualized as density curves that characterize multivariate niche divergence along the discriminant axis. Niche overlap is then measured with Schoener’s D (Schoener, 1968), which captures the proportion of shared density and ranges from zero (no overlap) to one (complete overlap). To distinguish different types of divergence, we further calculate four metrics following the two-dimensional niche divergence plane of Ascanio et al. (2024): 1) niche dissimilarity (the degree of non-overlapping density), 2) niche breadth exclusivity (the proportion of the discriminant axis range not shared between groups), 3) niche divergence magnitude (a composite index increasing with both dissimilarity and exclusivity), and 4) niche divergence angle (the relative contribution of density separation versus range exclusivity). This extends the niche divergence plane, originally developed for single environmental variables (Ascanio et al., 2024), to multivariate space. It quantifies both the amount and type of divergence, distinguishing density shifts within shared environmental space (weighted divergence), partial overlap in density and extent (soft divergence), nested environmental ranges (nested divergence), and little to no overlap in either density or extent (hard divergence) (Figure 1). For example, niche breadth exclusivity values near zero and divergence angle values near 90 degrees indicate divergence driven mainly by density differences within shared space, whereas niche breadth exclusivity values near one and divergence angle values near zero degrees indicate divergence driven more by exclusive range differences. Following Brown & Carnaval (2019), unequal environmental availability can be corrected by up-weighting rare and down-weighting common environments along the discriminant axis.

To identify variables contributing most to discriminant-axis separation, we summarize variable contributions. The discriminant axis is a linear combination of the retained PCs, which themselves are combinations of the original variables. By back-transforming this axis to the original variables, each receives a loading coefficient that reflects its weight. Variable contributions are then calculated as the squared loadings of each predictor normalized to sum to one, representing the relative proportion of discrimination explained by each variable. The direction of effect is determined by the sign of the loading in relation to the relative positions of the group centroids along the discriminant axis. Because PCA removes collinearity only for model fitting, contributions reflect shared covariance structure rather than independent effects and should be interpreted cautiously when predictors are strongly correlated.

We also visualize univariate density distributions of the top predictors. These are displayed as kernel density estimates for each group, linking the discriminant axis back to familiar ecological variables and revealing whether niche separation is driven primarily by shifts in means, dispersion differences, or divergence in distribution tails. Finally, we display the geographic basis of divergence analyses by plotting occurrence and background points on a basemap.

### Extraction of environmental layers and background

Using occurrence coordinates as the only input, our package generates species-specific background points and extracts a comprehensive standardized set of environmental variables (Supplementary Methods M1). These include abiotic and biotic environmental axes across fifteen datasets at monthly, seasonal, or annual temporal scales, such as climate, phenology, vegetation, soils, hydrology, terrain, land use, disturbance, and urbanization. We used the most up-to-date layers and multi-year averages where possible for each dataset, with most datasets representing conditions from the 2000s–2020s as static layers, single-year products, or recent multi-year composites, and a smaller subset representing longer-term climatologies spanning 1960–2000. The function standardizes data acquisition and raster processing across multiple environmental datasets while allowing flexible incorporation of user-supplied GIS layers. By extracting environmental values for occurrences and backgrounds within accessible areas, it generates the input for our test.

### DAPC preprocessing pipeline

We implement a standardized pipeline (Figure 1) to prepare occurrence and background data prior to DAPC and reduce common biases in niche divergence testing (Supplementary Methods M2): Accessible background areas are estimated by buffering convex hulls around species’ occurrences (alternatives: bounding boxes, concave hulls, point buffers). The buffer distance is flexible and should be chosen to approximate the dispersal capability or range expansion limits of the focal organisms (Barve et al., 2011). Occurrences are spatially thinned to reduce spatial autocorrelation and sampling bias (Dormann et al., 2007), and then downsampled to equal group sizes to avoid biases from unequal sample sizes in discriminant analysis (Lachenbruch & Goldstein, 1979). Environmental variables with negligible variability are excluded, as they provide minimal discriminatory power and can introduce numerical instability (Dormann et al., 2013). To reduce bias from non-analogous environments (Brown & Carnaval, 2019), variables lacking sufficient environmental analogy between species’ accessible backgrounds can be filtered out using univariate kernel density and bivariate histogram overlap. This removes variables showing little environmental overlap between species’ background spaces to ensure comparisons are restricted to environmental conditions available to both. Framing divergence tests within accessible background environments, while restricting comparisons to analogous conditions and incorporating seasonal predictors, minimizes bias from non-equilibrium distributions, where realized occurrences reflect only a subset of the fundamental niche (Brown & Carnaval, 2019).

We recommend conducting our DAPC niche divergence test first within analogous-only environments. Strong and significant divergence, or the absence of any remaining analogous variables after processing, suggests that the group’s niches have diverged within shared accessible environmental space (Brown & Carnaval, 2019). When niche divergence is low and non-significant, we recommend repeating the DAPC analysis on the full environmental dataset (analogous and non-analogous). If this test shows strong and significant separation, it indicates that both taxa occupy different environments. However, due to methodological constraints, such separation cannot by itself be used to infer the presence of adaptive niche divergence. Running DAPC on the full dataset is recommended even when the analogous-only analysis indicates strong separation, because restricting the analysis to analogous environments can exclude environmental axes along which niche divergence is present, thereby reducing discriminatory power and inflating type II error. When separation is low and non-significant in either analysis, the background test can be performed to evaluate whether this observed insignificance reflects limited statistical power.

### Simulations

To evaluate the robustness of our DAPC framework, we simulated multivariate niches for two simulation sets with 100 synthetic species each (Supplementary Methods M5). Environmental background data (see Empirical test) were projected into a 16-axis PCA space, and species-specific suitability surfaces were generated using Gaussian response functions with fixed standard deviations in the *virtualspecies R* package (Leroy et al., 2016). Multivariate divergence was introduced by progressively shifting species’ means away from the null species in two ways: across PC1–PC2 space capturing major axes of variation (set 1), and across PC7–PC8 mimicking subtler gradients of differentiation across minimal-variation axes (set 2). Suitability surfaces were rescaled to occurrence probabilities using a logistic function, from which presence-only records were sampled. This generated realistic synthetic distributions (Supplementary Figures S1–S2) and a controlled stepwise divergence gradient for evaluating the performance of niche tests.

### Empirical test – *Hemileuca maia* species group

We tested our framework using the *Hemileuca maia* species group (Saturniidae). These day-flying buck moths occur throughout the US and southern Canada and show considerable morphological and ecological variation (Lemaire, 2002; Tuskes et al., 1996). The complex is taxonomically challenging, with spurious species and population boundaries that have resulted in variation in the number of recognized species (Dupuis et al., 2018; Rubinoff et al., 2017; St Laurent, 2023). The group is also of significant evolutionary and conservation interest due to complex patterns of ecological adaptation in habitat and host use, coupled with incongruent genetic divergence (Dupuis et al., 2018; Rubinoff et al., 2017). Because larvae feed on specific plant groups, and pupation and adult emergence are tied to local soil conditions and seasonally restricted windows of temperature, humidity, and photoperiod (Lemaire, 2002; Tuskes et al., 1996), this group is an ideal test for our method that integrates multiple environmental axes.

We focus our analysis of niche divergence on western versus Great Plains populations, which have traditionally been considered to comprise a single species, *Hemileuca nevadensis* Stretch, 1872. This name has long been applied to all *maia*-group populations ranging from western North America into the Great Plains (Ferguson, 1971; Lemaire, 2002). Recently, Great Plains populations have also been associated with the name “*H. latifascia*” and proposed to represent a separate species-level taxon from western *H. nevadensis* (Schmidt, 2022, preprint). However, their taxonomic status remains unresolved because distributional gaps, morphological clines, host-use variation, and shared mitochondrial haplotypes with eastern populations complicate clear assignments (Dupuis et al., 2018; Schmidt, 2022; Tuskes et al., 1996). We discuss the details and taxonomic debate surrounding these taxa in Supplementary Methods M7. Here, we use “*H. latifascia*” and *H. nevadensis* as operational labels following Schmidt (2022), with the aim of using their broad distributions as a case study for testing *NicheDiv* while refraining from taxonomic conclusions.

We compiled up-to-date occurrence records for this species complex based on online and expert-curated datasets (Supplementary Methods M8–M9). We approximated each taxon’s accessible area using a convex hull around occurrences buffered by 5 km, from which we sampled 10,000 background points. We assembled an extensive spatial dataset of 328 monthly to annual variables comprising multiple environmental axes (Supplementary Figure S3), such as soil, climate, vegetation, land use, urbanization, photoperiod, burned area, and a predation pressure proxy. These variables were used by our preprocessing pipeline. We also consider two additional taxon pairings within the *H. maia* complex for testing niche divergence analyses across different spatial and environmental extents (Supplementary Methods M7).

### Comparisons with other niche divergence methods

Using our simulated and empirical datasets, we compared our DAPC approach against six other niche divergence tests (Supplementary Methods M6): 1) environmental PCA with equivalency and similarity tests in *ecospat* (henceforth, “*PCA-env*”) (Broennimann et al., 2012), 2) stratified PERMANOVA (permutational multivariate analysis of variance) (Anderson, 2001), 3) multivariate niche overlap based on hypervolumes (Blonder et al., 2014) and 4) the MVNH (multivariate-normal hypervolumes) framework (Lu et al., 2021), 5) the niche divergence plane (Ascanio et al., 2024) in PC1 and Schoener’s D in PC1–PC2 space of analogous variables (henceforth, “PCA-space”), and 6) logistic regression of group identity on environmental predictors. All niche tests were applied to each simulated and empirical dataset using our preprocessing pipeline.

## RESULTS

### Simulations – DAPC accurately recovers multivariate niche divergence gradients

We simulated 100 species that represented progressively greater multivariate divergence from a fixed null species. Across both simulation sets, our DAPC approach captured this increasing divergence with a gradual decrease in overlap values (Schoener’s D) and increasing niche divergence magnitude (Figure 2; Supplementary Figures S4–S6). Maximum divergence was quickly reached in simulation set 1, with species means shifted along major axes of variation (Figure 2). In simulation set 2, where divergence was introduced along low-variance axes, DAPC recovered a similar pattern, but divergence increased more slowly and reached a weaker maximum separation (Supplementary Figure S5).

**Figure 2:**
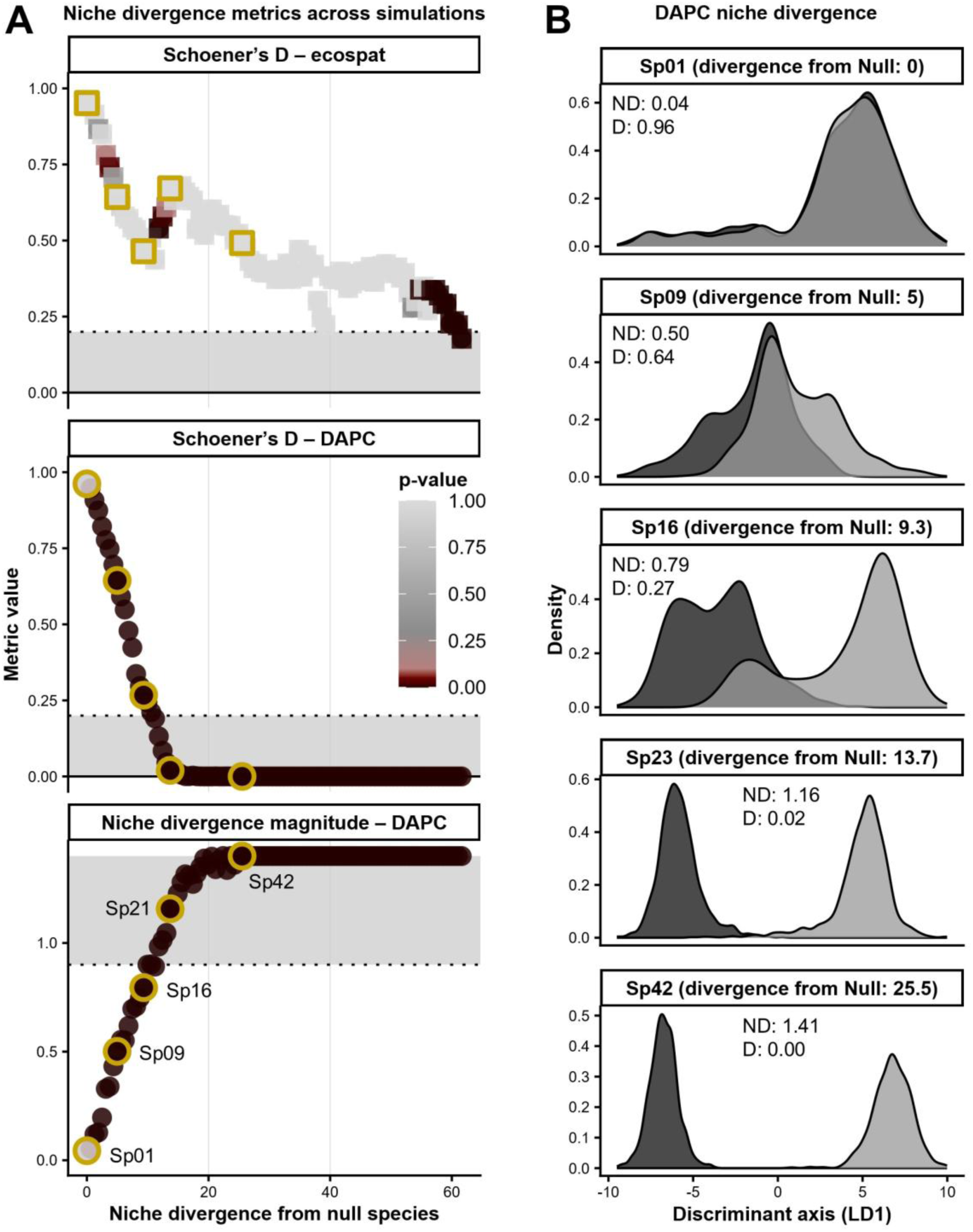
Multivariate niche divergence across 100 synthetic species simulated with increasing levels of divergence in PC1–PC2 suitability space. **A)** Divergence from the Null species (Sp00) vs. Schoener’s D and ND (niche divergence magnitude) from *ecospat PCA-env* and our DAPC method, with points representing simulations, focal species outlined in gold, point color encoding p-values, and grey areas representing strong niche divergence (D ≤ 0.2; ND ≥ 0.9). **B)** LD1 density curves for the focal species with increasing divergence from the Null species. Sp01 represents a simulated niche fully shared with Null species.

Permutation tests followed these gradients but indicated significance already at low to moderate divergence levels. Especially in simulation set 1, p-values became significant early in the divergence gradient, well before overlap values indicated moderate divergence (Figure 2A). This early significance reflects the high statistical sensitivity of discriminant analysis when divergence occurs along major axes of variation. Simulation set 2 showed a more gradual decline in p-values, consistent with the increasing simulated divergence along the low-variance axes (Supplementary Figure S5). Null comparisons representing a fully shared niche without divergence showed appropriate type-I error behavior in both simulation sets: p-values remained non-significant and divergence metrics indicated no separation (Figure 2; Supplementary Figure S7).

Other methods for assessing niche divergence generally captured the broad divergence gradient but showed less consistent patterns and lower estimated divergence than DAPC, particularly in simulation set 1 (Figure 2; Supplementary Figures S4–S6). Schoener’s D values from *PCA-env* were less consistent across the divergence gradient compared to our DAPC-based values. Although *PCA-env* captured the increase in divergence at a similar rate in simulation set 2, the decline in D was more compressed and erratic in set 1. Likewise, *PCA-env* equivalency-test p-values in simulation set 1 were inconsistent, with significant values at low divergence levels but non-significance at higher levels. PERMANOVA recovered the increasing separation along the gradient based on increasing test statistics and declining p-values. However, significance in simulation set 1 was often inconsistent with increasing divergence, and multivariate dispersion tests indicated that the PERMANOVA assumptions were frequently violated. Hypervolume and MVNH tests similarly showed higher divergence with increasing distance from the null, but their overlap estimates showed erratic behavior and increased again at high divergence levels in simulation set 1. PCA-space tests reproduced the same overall trend as DAPC but detected divergence less consistently, more slowly, and with lower maximum separation. Logistic regression reliably detected divergence based on its p-values, which followed a pattern similar to that from our permutation test.

### Empirical example – DAPC revealed strong niche divergence between “*H. latifascia*” and *H. nevadensis*

After data processing, background cropping, and thinning, we retained 167 occurrences and 10,000 background points for each of “*H. latifascia*” and *H. nevadensis* (Figure 3). Of the initial 328 environmental variables, 27 were removed due to negligible variance, 6 due to excessive missing data, and 261 due to non-analogous species’ backgrounds. The final dataset included 34 analogous variables.

**Figure 3:**
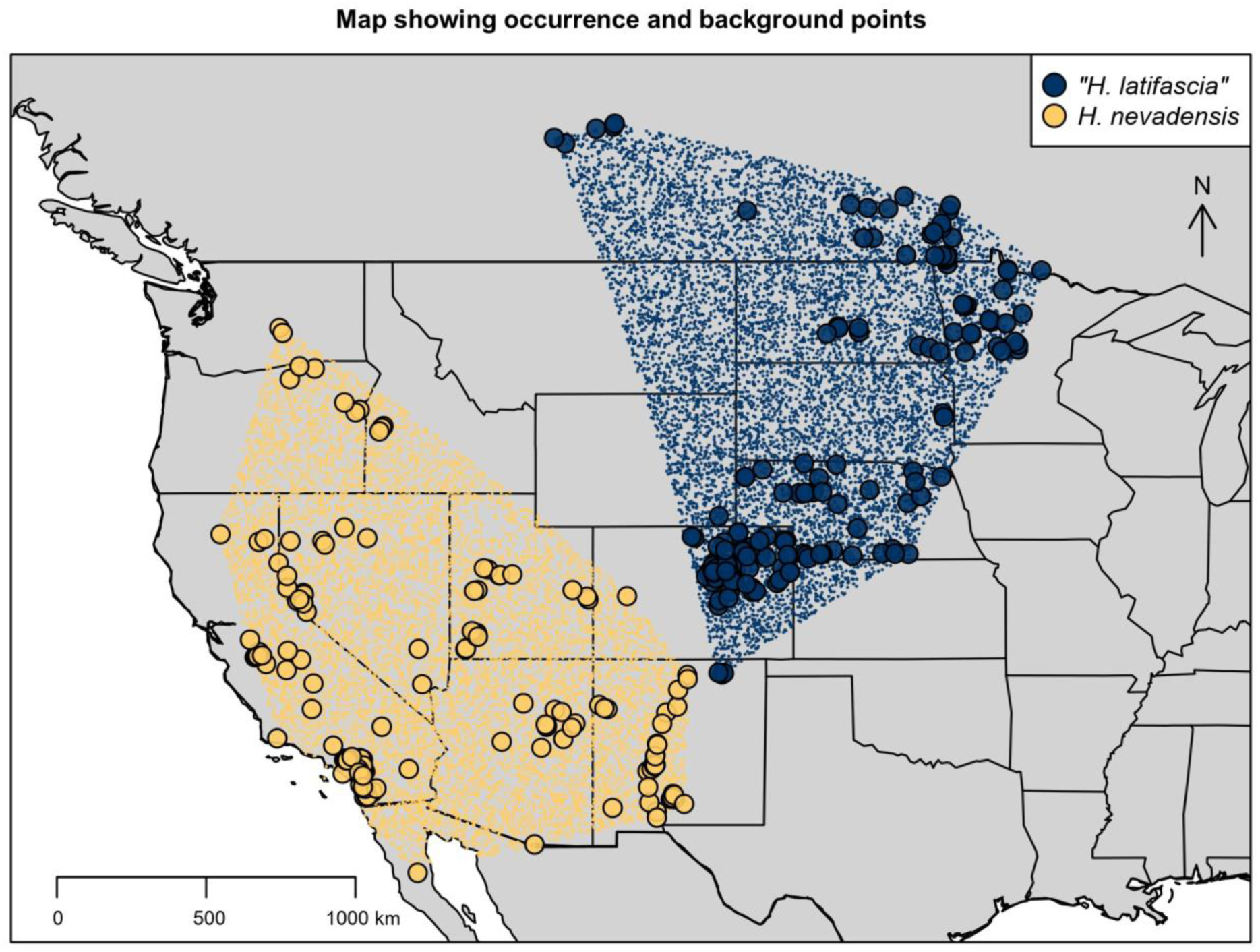
Occurrence (large points) and background records (small points) for “*Hemileuca latifascia*” and *H. nevadensis* across North America (empirical dataset 1). Shown are thinned and down-sampled records used as input for the DAPC niche divergence test.

The DAPC strongly separated the multivariate environmental niches of both taxa, with minimal overlap (Figure 4A; Supplementary Tables S2–S4), and the observed assignment accuracy was significantly higher than expected under a null distribution corresponding to a shared niche (Figure 4B). Our niche divergence metrics provided further evidence of strong ecological separation. Environmental overlap was essentially absent (Schoener’s D = 0.01), niche dissimilarity was nearly complete (NDS = 0.99), and breadth exclusivity was high (NE = 0.65). Divergence magnitude (ND = 1.18) indicated strong separation in both niche position and shape, while the divergence angle (θ = 56.7°) revealed a combination of range exclusivity and density differences. DAPC fit to the full environmental dataset without analogous restrictions supported strong divergence (Supplementary Table S3).

**Figure 4:**
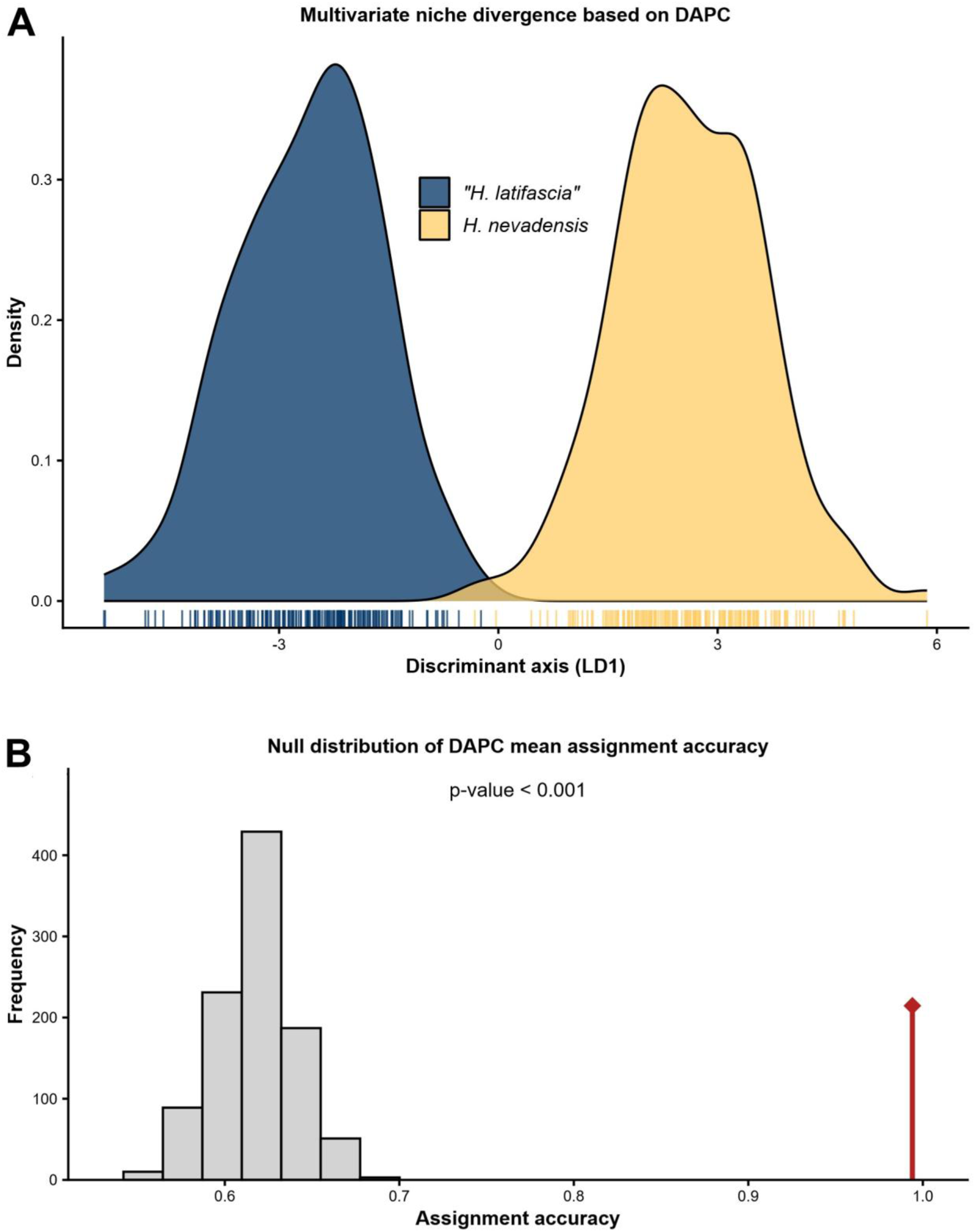
Discriminant analysis of principal components (DAPC) approach revealed strong multivariate niche divergence between “*Hemileuca latifascia*” and *H. nevadensis*. A) Divergence along the discriminant axis. **B)** Null distribution of classification accuracy from permutations, with the observed value shown as a red line.

Other niche divergence methods produced mixed support (Supplementary Tables S2–S4; Supplementary Figures S8–S10). Logistic regression rejected the null model of a shared niche. MVNH indicated strong dissimilarity that was primarily size-rather than centroid-driven. Hypervolume analyses revealed moderate overlap, with large exclusive fractions for each taxon. *PCA-env* produced intermediate overlap but supported niche divergence based on a significant niche equivalency test. PCA-space returned high overlap with low divergence values, and PERMANOVA indicated no significant centroid differences.

Variable contributions indicated that relative humidity overwhelmingly contributed to the separation between the two taxa, accounting for 98.1% of the discriminant axis (Figure 5A). “*Hemileuca latifascia*” was associated with lower relative humidity values in April, July, and October, whereas *H. nevadensis* was associated with higher values in March, August, September, and November. The remaining variables contributed marginally to the discriminant axis. Univariate distributions of the six strongest predictors broadly confirmed these patterns but also revealed more subtle contrasts (Figure 5B). Whereas the discriminant axis captured the combined multivariate “hard” separation, these univariate distributions showed different divergence scenarios, notably nested and weighted divergence.

**Figure 5:**
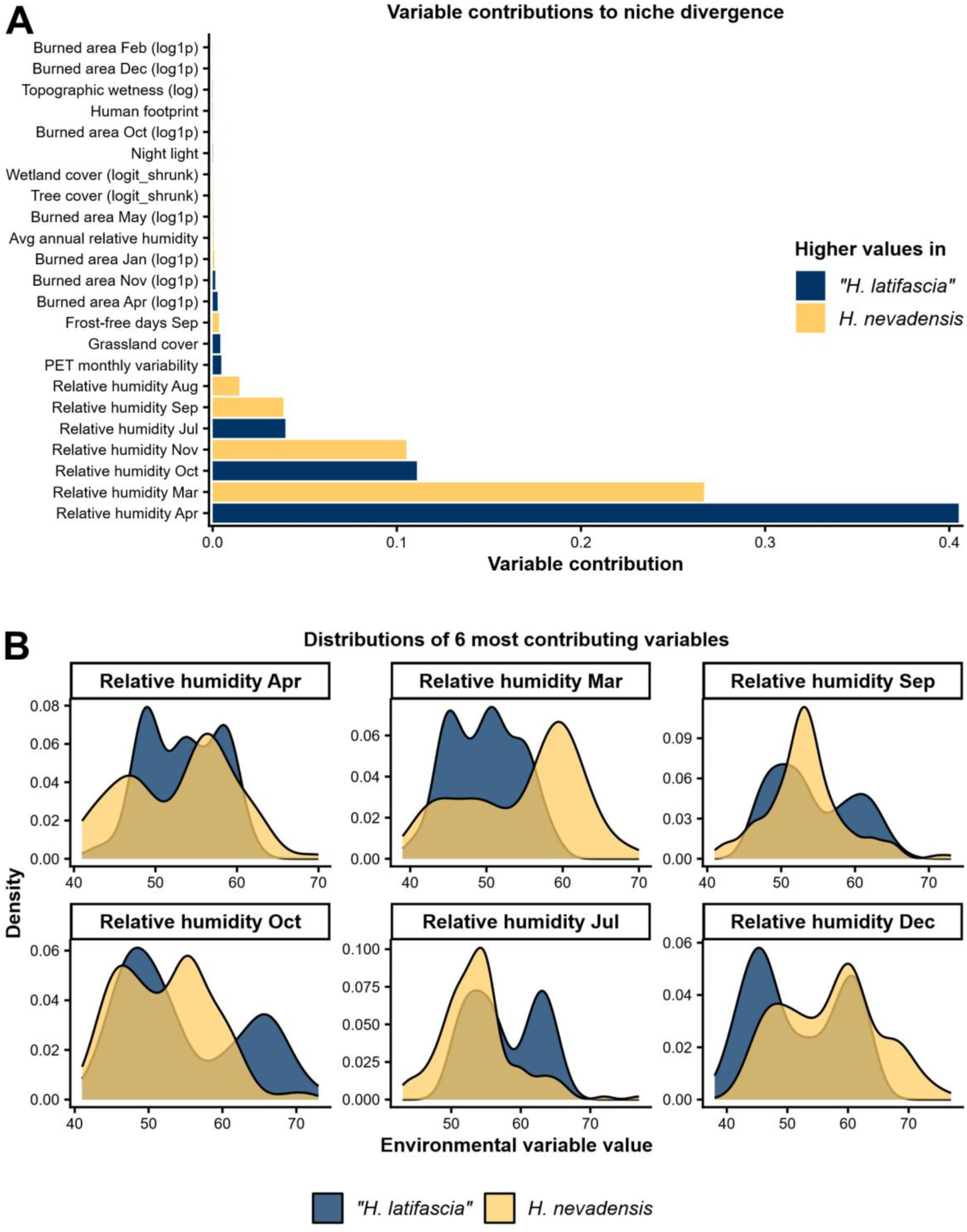
Variable contributions to multivariate niche divergence based on discriminant analysis of principal components (DAPC). **A)** Variable loadings to discriminant axis, with colors indicating the species with higher values (transformations in parentheses; not shown: 16 variables with contribution ≤ 0.01%). **B)** Distribution density plots of the six top-contributing variables (x-axis: percent relative humidity).

### Supplementary empirical examples reveal varying levels of ecological divergence

We used two additional taxon pairings in the *H. maia* complex to test the performance of niche divergence analyses across alternative empirical cases (Supplementary Methods M7). Niche divergence patterns varied in strength and underlying drivers (Supplementary Tables S3–S6). In the first pairing, divergence was moderate but significant and driven primarily by soil, land use, topography, and fire-related variables (Supplementary Figure S13). In the second example, vegetation productivity, burned area, and cropland cover contributed to a stronger divergence, with separation arising almost entirely from density differences (Supplementary Figure S19). In both cases, other niche divergence methods generally underestimated divergence relative to DAPC.

## DISCUSSION

We developed a framework for quantifying niche divergence using highly multivariate correlated environmental data, applicable to questions ranging from species delimitation, local adaptation, and community assembly to biogeography and macroevolution. By adapting DAPC from its primary use in population genetics (Miller et al., 2020) to environmental data together with a bias-reducing preprocessing pipeline and an automated environmental extraction function, we show that *NicheDiv* provides a robust niche divergence test. Across simulations and empirical examples, DAPC recovered the dominant axes of separation and different niche divergence types, while our divergence metrics scaled predictably with imposed divergence and maintained minimal type I error.

### A DAPC view of Hutchinsonian hypervolumes

The ecological niche is classically conceptualized as an *n*-dimensional hypervolume (Hutchinson, 1957). Recent methodological developments have operationalized this idea using kernel-density (Blonder et al., 2014) and parametric multivariate-normal hypervolumes (Lu et al., 2021). Such approaches quantify niche size, overlap, and shape in many dimensions, but interpretation becomes challenging as dimensionality and collinearity increase. Our DAPC test offers an alternative by using supervised group separation to identify the primary direction along which two realized hypervolumes differ. The resulting discriminant axis represents the combination of environmental variables that maximizes between-group separation, providing a simplified view of Hutchinsonian hypervolumes by projecting each species’ high-dimensional niche onto its most informative axis of divergence.

The advantage of this is evident in our empirical tests. Univariate density curves for the strongest individual predictors showed relatively subtle differences between taxa, whereas the multivariate discriminant axis resolved more reliable and consistent separation. This demonstrates that DAPC captures the multivariate structure underlying niche divergence–signals that are not apparent when predictors are examined individually.

### Advantages of DAPC relative to other niche divergence tests

Our comparisons with other methods for assessing niche divergence highlight several advantages of our method.

First, DAPC was the only method that consistently returned near-zero divergence when species shared the same niche and showed a progressive increase in divergence along the imposed gradient across both simulation sets (Supplementary Figures S4–S7). Divergence was generally detected earlier, and separation was stronger than in other methods, especially in simulation set 1, where divergence was simulated across major axes of variation. In the empirical datasets, DAPC achieved moderate to high ecological separation, whereas alternative methods returned mixed conclusions. PERMANOVA, *PCA-env*, and PCA-space approaches frequently failed to detect divergence across simulations even when ecological separation was strong. Hypervolume and MVNH metrics detected moderate to strong separation, but their effect sizes were sometimes erratic. While p-values from logistic regression detected divergence as reliably as DAPC, its goodness-of-fit measures were less consistent in simulation set 1.

A second advantage is that our permutation test provides estimates of significance via comparison with a null model, while hypervolume, MVNH, or PCA-space methods only return effect sizes. *PCA-env* equivalency and PERMANOVA tests provide p-values, but our simulations show that these do not reliably track divergence. Only logistic regression produced p-values that tracked divergence as consistently as our permutation test. Our results show that our permutation test is highly sensitive, yielding significant p-values already at low to moderate divergence levels. Therefore, significance should be interpreted jointly with divergence metrics and discriminant density curves. Similarly, non-significant results can reflect either true niche similarity or limited statistical power, given the environmental availability and sample sizes.

A third advantage is the ability of DAPC to summarize high-dimensional space to a single discriminant axis. By extending the univariate niche divergence plane of Ascanio et al. (2024) to operate on this multivariate discriminant axis (Figure 1: “Output”), our test identifies patterns that are lost in typical overlap metrics, namely position shifts, breadth differences, density shifts, and nested divergence. These types arise from distinct ecological scenarios, such as differences in specialization versus generalism, niche partitioning, asymmetric environmental tolerance limits, or unequal environmental filtering (Ascanio et al., 2024). The range of niche exclusivity and angle values in our simulations demonstrates that DAPC can recover all these divergence types. Other methods either operate in two-dimensional PCA space (*PCA-env* and PCA-space) or characterize divergence without a single interpretable axis (hypervolumes, MVNH, PERMANOVA, logistic regression). Unlike DAPC-based metrics, MVNH metrics also lack upper bounds, complicating interpretations.

Statistical robustness to multicollinearity is another major benefit of DAPC since environmental datasets typically contain many correlated variables, as in our empirical examples (Supplementary Figure S3). Most divergence tests perform poorly under collinearity, requiring either dimensionality reduction or manual variable selection. PERMANOVA, hypervolumes, and MVNH lack mechanisms to handle redundant structure and become numerically unstable with highly correlated variables. Logistic regression can include all predictors, but coefficient estimates become unstable under strong multicollinearity. *PCA-env* and PCA-space handle collinearity via PCA but are limited to 1–2 PCs. DAPC deals with multicollinearity by decorrelating predictors via PCA prior to discriminant analysis, improving numerical stability while retaining information from many correlated variables. Importantly, decorrelation occurs solely for model fitting while the original variables remain correlated.

DAPC handles high-dimensional multivariate data more effectively than alternative methods, allowing the input of hundreds of environmental variables. Our approach uses cross-validation to derive the optimal number of PCs, thus mitigating overfitting while retaining nearly all informative variation. In our analyses, DAPC routinely retained more than 95% of the environmental variance, far more than PERMANOVA, hypervolumes, MVNH, PCA-env, or PCA-space metrics (Supplementary Table S7). Retaining most of the total variance is important because ecological niches are inherently multivariate and species often diverge along variable combinations that may be lost when only 50–80% of variation is retained (Elith & Leathwick, 2009; this study). PC1–PC2-based approaches are constrained because they operate on one or two unsupervised components and therefore often omit ecologically relevant variation. Logistic regression was the only method that retained PC numbers with comparable or slightly higher variance than DAPC. However, the number of PC predictors in our logistic models was limited by the commonly used events-per-variable rule, which ties dimensionality to sample size rather than discriminatory performance (unlike DAPC’s cross-validated selection).

Linking divergence to environmental variables enables mechanistic hypotheses beyond pattern-only niche comparisons (Hutchinson, 1957; Peterson et al., 2011). DAPC identifies potential drivers of niche divergence by highlighting environmental variables that may be implicated in divergent adaptation among groups. Variable contributions are derived by back-transforming discriminant coefficients to the original variables, yielding scores that indicate which variables contribute most to separation. These scores reflect multivariate, covariance-informed contributions to group separation, so that variable contributions depend on the joint covariance structure rather than independent effects. Because the PCs themselves are combinations of collinear variables, the contributions cannot be interpreted as independent causal effects. High contributions may reflect correlated variable sets rather than single predictors, while low scores do not rule out biological importance if the variable’s signal is absorbed by correlated partners. Furthermore, restricting analyses to analogous environments may further occlude biologically important variables, so that inferred contributions reflect only shared environmental space. PERMANOVA, hypervolumes, PCA-space, and *PCA-env* lack variable-level contributions, preventing such interpretation. Logistic regression and MVNH provide variable-importance measures, but these differ fundamentally from the variable contributions returned by DAPC. Logistic regression coefficients describe how each environmental variable affects group identity conditional on the other predictors in the model. MVNH partitions variance and covariance across the original environmental axes into univariate and correlation components, characterizing how each environmental axis and the joint correlation structure contribute to niche volume and dissimilarity.

Finally, DAPC achieves competitive runtimes across our analyses despite using up to 290 variables and hundreds of permutations and cross-validations, with run times averaging 1.0–4.6 minutes in simulations and 0.17–0.49 minutes in empirical datasets (Supplementary Table S8). DAPC was slower than MVNH, PCA-space, PERMANOVA, and logistic regression, but faster than *PCA-env* and comparable to hypervolumes.

### Ecological divergence in the *Hemileuca maia* complex

In our focal empirical comparison, “*H. latifascia*” and *H. nevadensis* showed near-complete ecological separation within analogous environments, with DAPC yielding a single discriminant axis associated with high niche dissimilarity (Figure 4). Significant permutation and background-corrected tests using the full environmental dataset corroborated strongly divergent niches. These results provide support for ecological divergence between these proposed taxonomic units.

Variable contributions highlighted that seasonal relative humidity is potentially the primary axis of divergence, with “*H. latifascia*” associated with drier conditions in spring and mid-summer and *H. nevadensis* with more humid conditions during early spring and late autumn–winter. Density plots of these predictors showed subtler contrasts, indicating that the ecological signal emerges from coordinated shifts across multiple seasonal humidity variables. This pattern accords with the moths’ natural history, where larval host plants, pupation, and adult emergence are linked to local soil conditions, vegetation, and temporally restricted windows of favorable temperature and moisture, with adults flying primarily in late autumn and winter (Lemaire, 2002). Therefore, our results suggest that monthly moisture-related gradients across the Great Plains–Great Basin transition represent a key axis of niche differentiation between these taxa. Nonetheless, because preprocessing may exclude relevant correlated variables, and humidity may track associated conditions rather than act as direct drivers, experimental studies are needed to confirm the underlying mechanisms.

Ecological divergence is consistent with, but does not by itself demonstrate, ecological speciation. The environmental variables analyzed here describe realized niches under current climatic and land-use regimes, and contemporary distributions may lag behind shifts in fundamental niche space due to dispersal limitations, historical climatic change, and biotic interactions (Araújo & Pearson, 2005; Peterson et al., 2011). Thus, while both taxa have diverged to distinct present-day niches, the timing and processes underlying this divergence cannot be inferred from contemporary niche structure alone and require demographic and phylogeographic analyses.

### Multiple axes of seasonal environmental variables are crucial

Our empirical examples underscore that multiple environmental axes beyond annual climate can be the predominant dimensions of niche divergence. Across all species pairs, the strongest ecological differentiation involved temporally explicit, non-climatic variables rather than annual means or extremes, particularly seasonal humidity, soils, vegetation, disturbance, and land-use variables. This aligns with evidence that species’ realized niches are shaped by seasonal physiological requirements, resource availability, phenological dynamics, habitat structure, and disturbance regimes (Elith & Leathwick, 2009; Kearney & Porter, 2009; Prajzlerová et al., 2025), none of which are captured by static annual averages. Multivariate approaches such as our DAPC-based framework are essential for integrating these numerous predictors.

In our main empirical analysis, divergence was dominated by monthly relative humidity variables. These contributed in opposite directions across the year, reflecting contrasting intra-annual moisture regimes (Figure 5). These differences would collapse into similar values if summarized as annual means, masking true niche structures. The discriminant axis thus reflects within-year heterogeneity that only emerges when multiple seasonal variables are analyzed jointly.

Our environmental extraction function facilitates the automated extraction of a range of environmental variables, including custom layers. By leveraging increasingly available high-resolution global datasets (Batjes et al., 2024; Hufkens et al., 2018; Potapov et al., 2021), it moves niche analyses beyond typical annual bioclimatic variables toward richer seasonal abiotic and biotic descriptors that better reflect phenology, resources, and organismal physiology (Elith & Leathwick, 2009; Kearney & Porter, 2009; Title & Bemmels, 2018). Nonetheless, better representation of aquatic environments, microhabitat partitioning, and direct biotic interactions will require further development of global high-resolution GIS layers.

## Limitations

Despite the outlined advantages, several limitations should be considered. First, *NicheDiv* quantifies niche divergence as realized niche differentiation, not the evolutionary process that generated these patterns. Analogous filtering can reveal that this differentiation occurs under conditions available to both groups (Brown & Carnaval, 2019), but occurrence–environment data alone cannot identify the underlying mechanism (Araújo & Pearson, 2005; Peterson et al., 2011). Second, DAPC identifies the linear combination of variables that maximizes separation between groups. Accordingly, non-linear or interaction-driven niche differences are represented only as linear approximations. In such cases, more complex machine-learning or non-parametric hypervolume methods may better capture these patterns, particularly when species differ along curved or discontinuous boundaries. Nonetheless, robustness to non-linearity is indicated by DAPC capturing imposed divergence reliably in our analyses, and its successful application to high-dimensional genetic data (Jombart et al., 2010; Miller et al., 2020). Third, our test currently includes only continuous or proportional environmental variables. Given its common application to biallelic genetic markers (Miller et al., 2020), our DAPC approach could be extended to incorporate binary or categorical data (e.g., presence/absence for host, habitat, symbiont, or pollinator types). Nevertheless, combining continuous and binary data is challenging due to different variance structures, requiring further development or advanced machine-learning methods. Finally, although DAPC can accommodate multiple groups, our test focuses on pairwise comparisons with the advantage of a single interpretable discriminant axis. Multiple groups can be analyzed through separate pairwise tests.

## Contributions

Conceived study: DS, ZGM, and JRD. Occurrence data compilation: DS, JRD, and BCS. Environmental data extraction: DS supervised by ZGM. Statistical analyses, code development, and figures: DS supervised by ZGM and JRD. First draft: DS supervised by ZGM and JRD. Second draft: DS, with input from all authors. Contribution to final manuscript: All authors.

## Funding

Funding was provided by a USDA-NIFA HATCH grant to JRD (Project KY008091).

## Conflicts of interest

The authors have no conflicts of interest to declare.

## Acknowledgments

We thank Stefan Schönberger (VISUELL Studio für Kommunikation GmbH, Germany) and Adrian Schönberger (Lufthansa Technik, Germany) for assistance with the graphical design of some figures. We are also grateful to Jim P. Tuttle (independent researcher, USA) and Nathan Haan (University of Kentucky, USA) for helpful comments on the manuscript.

## Data availability statement

Data and code supporting this study are available from Zenodo (DOI to be inserted).

## Notes

### Competing Interest Statement

The authors have declared no competing interest.

