## Supplement for "*NicheDiv*: A DAPC framework to quantify niche divergence across highly multivariate environmental space"

**Author affiliations**
[1]Department of Entomology, University of Kentucky, Lexington, Kentucky, USA
[2]Department of Entomology, University of California, Riverside, California, USA
[3]Canadian National Collection of Insects, Arachnids and Nematodes, Agriculture and Agri-Food Canada, Ottawa, Ontario, Canada

**This PDF includes:**
Supplementary Methods: M1–M9
Supplementary Figures: S1–S26
Supplementary Tables: S1–S9
Supplementary References

### Supplementary Methods

#### I) DAPC method implementation (package and analytical framework)

##### M1: Extraction of environmental layers and background points

Our *extract.env.and.background* function automates the extraction of a full set of environmental variables for both occurrence and background data. This generates a comprehensive, multivariate environmental dataset for the DAPC (discriminant analysis of principal components) niche divergence test, including the background data required to account for non-analogous bias (see Supplementary Methods M2.7). The function integrates data acquisition, masking, projection, and raster extraction into a single streamlined workflow, ensuring spatial consistency, reproducibility, and simple handling across multiple environmental data sources.

1. **Input and setup.** As the main input, the function requires an occurrence data frame (*occurrence.data* argument) with unique row names (e.g., specimen or record IDs) and containing longitude and latitude columns. The coordinate reference system (CRS) of the occurrence coordinates is specified via the *CRS.occurrences* argument (default: EPSG:4326). Thus, the function can accept either geographic or projected coordinates based on the specified CRS. This data frame can include two or multiple lineages, taxa, species, subspecies, or populations (hereafter "species"). Environmental datasets to extract can be specified using the *env.datasets* argument (see Step 5). Background data is optionally generated from the accessible area using *generate.background.data = TRUE*; optional parameters include the number of random background points (*N.background.poin*ts argument) and spatial buffer distance of accessible area (*buffer.km* argument). Intermediate files and cached CSVs are stored in a designated subfolder (*intermediate.files.dir* argument) and can be automatically deleted when *delete.intermediate.files.folders = TRUE*. Existing intermediate CSV files are reused unless *overwrite = TRUE*, while previously downloaded environmental files (rasters, ZIP archives, or auxiliary datasets) are reused unless *redownload.rasters = TRUE*.

2. **Study extent and accessible area**. Occurrence coordinates are converted to *sf* objects using the *st_as_sf* function in the *sf v.1.0.21 R* package (Pebesma & Bivand, 2023). If a geographic CRS is detected via the *st_is_longlat* function in *sf*, the function automatically switches to an equal-area projection for geometric operations. Within continental North America, coordinates are projected to EPSG:5070 (NAD83 / Conus Albers) if the study extent falls within the Conus Albers bounding box, as detected using the *relate* function in the *terra v.1.8.7 R* package (Hijmans, 2025). Elsewhere, a locally centered Lambert Azimuthal Equal-Area projection is created dynamically from the centroid of the occurrences using the *st_union* and *st_centroid* functions in *sf*. For polar or extreme regions, EPSG:6933 (World Equal Area) is used as a fallback. If the input data are already in a projected CRS with meter units, geometric operations, including convex-hull construction with the *st_convex_hull* function and buffering with the *st_buffer* function, are performed directly without reprojection. Otherwise, the data are projected to a meter-based equal-area

CRS before these operations. The accessible area (M-space), used to generate background points (see Background Sampling), is defined as the convex hull of all occurrence coordinates buffered by the user-defined distance (*buffer.km* argument) via the *st_buffer* function in *sf*. The study area is defined as this accessible area further buffered by the minimum resolution of the coarsest raster (e.g., one kilometer) plus five kilometers to prevent NA extraction artifacts along raster edges (especially for terrain metrics). The buffered polygon is reprojected back to the input CRS (*CRS.occurrences argument*) via the *st_transform* function in *sf* to ensure distance-preserving buffering and accurate area delineation.

3. **Landmask generation.** If background points are to be created (*generate.background.data = TRUE*; see step 4) and/or terrain metrics are estimated (*env.datasets = "terrain"*; see step 5.2), a land-only spatial mask ("landmask") is generated as follows. Country polygons from GADM version 3.6 (geodata.ucdavis.edu/gadm) are downloaded as RDS files and restricted to those intersecting the study extent (see step 2) via the *relate* and *crop* functions in *terra*. The resulting polygons are disaggregated via the *disagg* function in *terra*, and only the *N* largest pieces (*landmask.largest.N.pieces* argument; default: *5*) are retained to avoid small offshore fragments. Polygon areas are computed via the *expanse* function in *terra* to identify the largest pieces. If *remove.hydrolakes.background = TRUE*, the function optionally removes major lakes from the landmask using the *HydroLAKES v.10* dataset from *HydroSHEDS* (hydrosheds.org) to ensure that background points are restricted to ecologically accessible terrestrial areas rather than aquatic surfaces (N. Barve et al., 2011; Soberón, 2007). The download of the *HydroLAKES* file is facilitated via regional or global versions from Zenodo (doi.org/10.5281/zenodo.17503890). Regional HydroLAKES files are identified using the *relate* function in *terra* to match the study-area extent, downloaded with the *download.file* function in the *base R utils v.4.4.1 package* (R Core Team, 2025), unzipped with the unzip function in *utils*, and read as vectors using the *vect* function in *terra*. Regional bounds are defined using the *ext* function in *terra*. Lake polygons are removed from the landmask using the *erase* function in *terra* to restrict the final mask to terrestrial areas. *HydroLAKES* vectors are saved and loaded as RDS files using the *saveRDS* and *readRDS* functions in *base* to minimize repeated downloads. Users may skip this step (using *remove.hydrolakes.background = FALSE*) if the distributional extent is very large (this operation can take up to hours to run for continental-scale datasets) or to include aquatic environments in the background. The final study-area and accessible-area polygons are returned in the CRS specified by the *CRS.occurrences* argument, but geometric operations are carried out in meter-based equal-area CRSs, and the HydroLAKES sampling mask is rasterized in EPSG:4326. Existing landmask and *HydroLAKES* files are reused, unless *redownload.rasters = TRUE*.

4. **Background sampling.** If *generate.background.data = TRUE*, our function optionally samples background points from the accessible area after terrestrial masking (required for the non-analogous bias correction of the DAPC divergence test: Supplementary Methods

M2.7). When *remove.hydrolakes.background = FALSE*, points are sampled from the intersection of the accessible area and the landmask. When *remove.hydrolakes.background = TRUE*, candidate points are sampled from the accessible area and retained only if they fall on land cells in a HydroLAKES-based raster mask. A uniform random sample (*N.background.points* argument; default: *100,000*) is drawn from within the accessible area (see Step 2) using the *spatSample* function in *terra*. Although smaller values are usually sufficient, this default is intentionally generous to accommodate large geographic extents or analyses involving multiple species followed by later downsampling and subsequent data splitting. Coordinates are exported as a CSV file (*csv.background.out.file* argument). When *remove.hydrolakes.background = TRUE*, background points are sampled exclusively from land raster cells of a *HydroLAKES*-based raster mask. To create this raster mask, *HydroLAKES* polygons overlapping the accessible area are cropped using the *st_crop* function in *sf*, rasterized with the *rasterize* function in *terra* at approximately 8.1-arcsecond spatial resolution, and overlaid on a template raster where lake raster cells are assigned NA and land raster cells are set to 1. The resulting mask is saved as a GeoTIFF file (*HydroLakes_sampling_mask.tif*) and reused to avoid recomputation. Rasterizing the *HydroLAKES* polygons at fine resolution substantially accelerates subsequent sampling because lake–land boundaries are resolved once at the raster level rather than repeatedly through polygon intersections. The function then repeatedly samples candidate batches of up to 50,000 points using the s*patSample* function in *terra* and extracts raster cell values from the raster mask with the *extract* function in *terra*. Points falling on lakes (NA mask values) are discarded, and sampling continues iteratively until the target number of valid background points (*N.background.points* argument) is reached or the maximum batch limit is hit. Valid background points are accumulated across candidate batches in memory and written once to a temporary CSV only when sampling terminates, either after the target number of valid points has been reached or after the maximum number of sampling batches has been exhausted. The final coordinates are written to a CSV file (*csv.background.out.file* argument). During *HydroLAKES* cropping, *s2* spherical geometry is temporarily disabled and restored via the *sf_use_s2* function to avoid topological errors. When HydroLAKES removal is disabled, polygon validity checks are applied when necessary. In this case, *st_intersection* is attempted first with the *s2* geometry. If invalid polygons are detected, the function applies the *st_make_valid* function and retries under planar geometry (*s2* disabled). For both options, the resulting polygons are converted between *sf* and *terra* objects using the *st_as_sf* and *vect* functions for compatibility. Random sampling is reproducible via the *seed* argument.

5. **Environmental data extraction.** The function extracts environmental variables for all occurrence records and (optionally) background points from a series of standardized datasets. For this, occurrence and background coordinates are converted to SpatVector objects in the target CRS using the *vect* function in *terra*. Before extraction, the function checks whether the CRS of the input coordinates matches that of each raster using the *crs* and *same.crs* functions in *terra*. If they differ, the coordinate vectors for occurrence and

background points are temporarily reprojected to the raster's native CRS with the *project* function in *terra*. The rasters themselves are not reprojected, minimizing resampling artifacts and drastically reducing computational time. Environmental values are extracted using the *extract* function in *terra*, ensuring spatially precise correspondence between each point and its native raster cell. Each dataset is processed by a modular internal routine that downloads raster or zip archives with a robust multi-attempt downloader, including file-size checks, unzips, and performs extraction for all points. For high-resolution rasters and large files, regional or continental rather than global files are identified and used automatically using the *relate* function (*relation = "within"*) in *terra* to match the study extent. Incomplete downloads are automatically retried. Intermediate rasters are stored in the *Env_rasters* subdirectory within the output directory (*rasters.dir* argument) and can be deleted automatically at the end of the process (if *delete.intermediate.files.folders* = TRUE) to avoid accumulating large high-resolution files. *Terra* temporary files are cleaned between datasets. The following fifteen environmental datasets can be specified for extracting via the *env.datasets* argument: *"elevation"*, *"ClimateNA"*, *"EVI"*, *"terrain"*, *"ENVIREM"*, *"footprint"*, *"landcover"*, *"soil"*, *"forest_height"*, *"atmosphere"*, *"nightlight"*, *"burned_area"*, *"snow_water_equivalent"*, *"daylength"*, and *"soil_moisture"*.

i. **Elevation.** Elevation (*env.datasets = "elevation"*) is obtained from the Copernicus Global Digital Elevation Model GLO-90 v.1.1 (doi.org/10.5069/G9028PQB). The data were mean-aggregated from the native 90 meters to a 250-meter spatial resolution using the *aggregate* function (*fact = 3, fun = mean*) in *terra*. The function downloads global and continental 250-meter spatial resolution tiles from Zenodo (10.5281/zenodo.17448949; 10.5281/zenodo.17458752; 10.5281/zenodo.17450333; 10.5281/zenodo.17485342). We recommend not retaining elevation in the final environmental dataset, as it is not a direct ecological predictor of species distributions (Dormann et al., 2013; Guisan & Zimmermann, 2000; Title & Bemmels, 2018), but instead using it solely to derive terrain metrics and *ClimateNA* data (see below).

ii. **Terrain variables.** From the elevation raster, five terrain metrics are derived as described below (*env.datasets = "terrain"*; requires *env.datasets = "elevation"*). Following Wilson et al. (2007) and using the *terrain* function in *terra*, we obtain the terrain ruggedness index (TRI; the mean of absolute differences between the elevation of a focal cell and each of the eight surrounding cells), topographic position index (TPI; the difference between the elevation of a focal cell and the mean of the eight surrounding cells), and roughness index (the difference between the maximum and minimum elevation of a focal cell and the eight surrounding cells). A three-by-three cell neighborhood (*neighbors = 8*) is used in all local terrain calculations to capture fine-scale topographic variation while minimizing smoothing effects and boundary artifacts. This represents the standard kernel for geomorphometric derivatives (Conrad et al., 2015; Hengl & Evans, 2009). Next, the heat load index (HLI; predicts solar radiation as a function of the slope and aspect of each cell) is estimated following McCune & Keon (2002) using slope and aspect derived from the elevation raster. Slope and aspect are computed by setting *v = "slope"* and *v = "aspect"* in the *terrain* function, both with units set to radians (*unit = "radians"*). Latitude is obtained via the *init* function (*elevation_raster, fun = "y"*) in *terra* and converted to radians. Based on McCune & Keon (2002), HLI is calculated for each cell as 0.339 plus 0.808 times the cosine of latitude multiplied by the cosine of slope, minus 0.196 times the sine of latitude multiplied by the sine of slope, minus 0.482 times the cosine of aspect minus 225 degrees multiplied by the sine of slope. This formulation approximates relative potential solar radiation as a function of slope, aspect, and latitude and is widely used as a topographic surrogate for heat load and microclimatic energy balance (Kumar et al., 1997; McCune & Keon, 2002). Finally, the topographic wetness index (TWI) is estimated, quantifying soil-moisture potential as a function of slope and upstream contributing area (Beven & Kirby, 1979). TWI is computed in projected, meter-based coordinates to ensure hydrological accuracy by projecting the elevation raster to EPSG:5070 within North America or to a locally centered Lambert Azimuthal Equal-Area CRS elsewhere. TWI is then computed using the *whitebox v.2.4.3 R* package (Lindsay, 2016; Wu & Brown, 2022), which implements the D8 flow algorithm. Following standard hydrological preprocessing procedures and recommendations (Lindsey, 2023; R. Sørensen et al., 2006), the elevation surface is hydrologically corrected using the *wbt_fill_single_cell_pits* and *wbt_breach_depressions_least_cost* functions (with a breach distance threshold of four cells to minimize artificial channel incision; *dist = 4*). Then, the slope in degrees is calculated using the *wbt_slope* function with a z-factor (defined as one divided by 111,320 times the cosine of the mean study-area latitude). The specific contributing area is computed using the *wbt_d8_flow_accumulation* function (*out_type = "sca"*), and the wetness index is generated using the *wbt_wetness_index* function (following the classical formulation in which the index equals the natural logarithm of the upslope contributing area divided by the tangent of the slope). Before extraction, occurrence and background coordinates are reprojected to match the TWI raster CRS to ensure spatial alignment between points and hydrologically derived terrain metrics. Overall, these five terrain metrics describe complementary aspects of landscape structure that jointly influence local energy balance, hydrology, and habitat heterogeneity, thereby affecting microclimate, soil development, and vegetation distribution (Bennie et al., 2008; Beven & Kirby, 1979; Dobrowski, 2011; Wilson et al., 2007). Computations are performed with the *terraOptions* function (*tempdir = terrain_dir, todisk = TRUE, memfrac = 0.6, progress = 1*) in *terra* to optimize memory use during large raster operations.

iii. **Climatic and phenological variables.** Available for North America, thirty-eight monthly and annual *ClimateNA* climatic variables are extracted (Wang et al., 2016) (*env.datasets = "ClimateNA"*; requires *env.datasets = "elevation"*). Multi-year mean values for 2011–2020 are obtained using the *ClimateNAr* function (*varList = "YM",* *periodList = "Decade_2011_2020.dcd")* in the *ClimateNAr v.3.1.0 R* package (climatena.ca). This function dynamically generates scale-free climate data based on input latitude, longitude, and elevation by combining bilinear interpolation with local elevation adjustment to downscale Parameter-elevation Regressions on Independent Slopes Model climate averages ("normals") to point-level resolution (Wang et al., 2016). This approach provides higher topographic precision than other gridded climatic datasets such as *WorldClim* (MacDonald et al., 2025; Wang et al., 2016). All monthly and annual formulations result in 205 variables (25 annual, 180 monthly) (Supplementary Table S9). The *ClimateNAr* function is internally replicated to suppress console messages and progress output for reproducibility. The *ClimateNAr* package (only available for Windows systems) is downloaded via Zenodo to facilitate access (doi.org/10.5281/zenodo.17401569). In addition to *ClimateNA*, sixteen *ENVIREM* (Environmental Rasters for Ecological Modeling; envirem.github.io) variables are extracted (*env.datasets = "ENVIREM"*). These are calculated as long-term yearly averages spanning 1960–1990, representing bioclimatic variables relevant to species' ecology and physiology, thus improving niche modeling performance relative to other climatic datasets (Title & Bemmels, 2018). The sixteen rasters are downloaded from Zenodo (doi.org/10.5281/zenodo.17507776) as continent-, region-specific or global archives. Each layer is provided at 30-arcsecond spatial resolution, corresponding to approximately one kilometer at the equator. *ClimateNA* and *ENVIREM* variables are complemented by three *WorldClim v.2.1* climatic metrics at 30-arcsecond spatial resolution (Fick & Hijmans, 2017; Hijmans et al., 2005): solar radiation (srad), wind speed (wind), and vapor pressure (vapr) (env.datasets = "atmosphere"). Data is extracted from Zenodo (doi.org/10.5281/zenodo.17429723) as continent-, region-specific, or global archives to facilitate downloading (six layers per variable), representing mean conditions for paired months across 1970–2000. These variables capture atmospheric energy input (srad), mechanical mixing and desiccation potential (wind), and ambient vapor pressure (vapr), which influence evapotranspiration and physiological water stress (Allen et al., 1998; Novick et al., 2016). The median, minimum, and maximum values across the six bimonthly layers are also computed for each metric. Finally, *Daymet v.4* daily surfaces (earthdata.nasa.gov/data/catalog/ornl-cloud-daymet-daily-v4r1-2129-4.1) for daylength (dayl) and snow water equivalent (SWE) are retrieved (*env.datasets = "daylength"* and *"snow_water_equivalent"*; available only for North America). *Daymet* is a research product of the Environmental Sciences Division at Oak Ridge National Laboratory (daymet.ornl.gov). Both variables are derived from monthly mean composites at 30-arcsecond spatial resolution in the native Daymet Lambert Conformal Conic projection and downloaded via Zenodo (daylength: doi.org/10.5281/zenodo.17468681; SWE: doi.org/10.5281/zenodo.17495169). For daylength and SWE, monthly means from 2024 and 2023 are extracted, respectively. Each raster comprises twelve layers representing calendar months. These monthly means can be further aggregated within two-month windows or averaged across years to generate stable bimonthly climatologies when multiple years are available. Daylength provides a direct measure of photoperiod duration as a fundamental driver of circannual phenology. Snow water equivalent reflects accumulated snowpack and, hence, winter habitat accessibility and surface moisture regimes. Together, these variables capture seasonal light and cryospheric constraints on ecological and physiological processes (Hufkens et al., 2018).

iv. **Vegetation variables.** Bimonthly *EVI* (Enhanced Vegetation Index) data at 250-meter spatial resolution for 2020 are derived from *OpenLandMap* (stac.openlandmap.org/evi_mod13q1.tmwm.inpaint/collection.json) (*env.datasets = "EVI"*). *EVI* is derived from Moderate Resolution Imaging Spectroradiometer *MOD13Q1* products with values aggregated into two-month composites using the 90th percentile, quantifying vegetation greenness and canopy structure while reducing saturation in high biomass areas compared to the commonly used Normalized Difference Vegetation Index (Huete et al., 2002). The median, minimum, and maximum EVI values are also calculated across the year. Download is facilitated via regional, or global zip files from Zenodo (doi.org/10.5281/zenodo.17449850). Next, global land-cover fractions for 2020 were downloaded from the *European Space Agency WorldCover Project* (esa-worldcover.org) using the *landcover* function in the *geodata v.0.6.6 R* package (Hijmans et al., 2024) (*env.datasets = "landcover"*). These layers quantify the proportional coverage of eleven land-cover classes (trees, grassland, shrubs, cropland, built, bare, snow, water, wetland, mangroves, and moss) within each grid cell. The *WorldCover* product represents a single-year composite classification derived from Sentinel-1 and -2 imagery, providing a high-resolution characterization of heterogeneous landscapes.

v. Finally, forest height (*env.datasets = "forest_height"*) is extracted from derived canopy-height rasters based on the ETH Global Canopy Height 2020 product. This is a global 10-m canopy top height model estimated from Sentinel-2 imagery with *NASA* Global Ecosystem Dynamics Investigation waveform Light Detection and Ranging supervision (Lang et al., 2023). Forest height provides spatially explicit estimates of canopy height as a structural measure of vegetation complexity, vertical habitat structure, and shade availability (Potapov et al., 2021). We downloaded the original ten-meter source tiles (https://langnico.github.io/globalcanopyheight/; research-collection.ethz.ch/handle/20.500.11850/609802) using custom Python scripts (available at Zenodo: doi.org/10.5281/zenodo.19796077) that first generated continent-specific download lists from predefined geographic extents and the ETH Global Canopy Height tile naming scheme, then validated the generated URLs and downloaded only missing tiles via the command-line download manager *aria2c v.1.37.0* (aria2.github.io/) with retry logic and resumable transfer support. The downloaded files were then processed with custom *R* scripts (available at Zenodo:

doi.org/10.5281/zenodo.19796134). For each continent, the downloaded GeoTIFF tiles were checked for readability and CRS consistency, assembled into a virtual raster mosaic, and used to derive a densified source-footprint polygon. This footprint was projected to a continent-specific equal-area CRS and used to define an output template that reduced clipping along curved projection edges. Each continental virtual raster mosaic was then projected and aggregated directly to a compressed GeoTIFF at 250-meter spatial resolution. Realm-scale products were generated in a second step from the continental rasters by projecting them to shared equal-area templates derived from projected non-NA footprints and mosaicking them into combined outputs. The global product was produced similarly, but written at 500-meter spatial resolution in a world-scale equal-area projection. Download is facilitated via regional, continental, or global Geotiff files from Zenodo (doi.org/10.5281/zenodo.19616887).

vi. **Soil variables.** Nine soil variables are derived from *SoilGrids* (Poggio et al., 2021; Turek et al., 2023) at 30-arcsecond spatial resolution using the *soil_world* function in *geodata* (*env.datasets = "soil"*). *SoilGrids* represents baseline soil properties around 2017 predicted based on 2015–2019 covariates and estimated by machine-learning models trained on World Soil Information Service soil profiles and remote-sensing predictors. The topsoil layer (zero to five centimeters) is used, which is most relevant to vegetation–soil interactions and surface ecological processes (Weil. R. & Brady, 2016). Variables included bulk density of the fine earth fraction (bdod), volumetric fraction of coarse fragments larger than two millimeters (cfvo), clay fraction (clay), sand fraction (sand), silt fraction (silt), total nitrogen content (nitrogen), soil pH in $H_2O$ (phh2o), soil organic carbon content (soc), and organic carbon density (ocd). These properties influence water availability, nutrient cycling, and vegetation growth, thus directly shaping species' ecological niches (Hillel, 1998; R. B. Jackson et al., 1996). Download is facilitated via regional or global zip files from Zenodo (doi.org/10.5281/zenodo.19614206). Furthermore, (volumetric) soil moisture is extracted from the European Space Agency Climate Change Initiative Soil Moisture *COMBINED v09.1* product (climate.esa.int/en/projects/soil-moisture) (Dorigo et al., 2017; Gruber et al., 2019; Preimesberger et al., 2021) at 900-arcsecond spatial resolution corresponding to approximately 25 km at the equator (*env.datasets = "soil_moisture"*). The product integrates multiple active and passive microwave retrievals (e.g., ASCAT, SMOS, AMSR-E, AMSR2, SSM/I, TMI) into a harmonized global time series using a triple-collocation-based scaling and merging framework. Six global layers are used, representing climatological bimonthly means (*soil_moisture_01 – soil_moisture_06*), averaged across 2018–2023 from daily Level-3S *COMBINED* NetCDF data. These are downloaded from Zenodo (https://doi.org/10.5281/zenodo.17496279).

vii. **Urbanization variables.** Human footprint data for 2009 are retrieved at 30-arcsecond spatial resolution using the *footprint* function in *geodata*. We use the Human Footprint Index as a proxy for urbanization and broader human landscape modification. Specifically, the index is a cumulative measure of human pressure on the environment that integrates eight anthropogenic factors: built environments, population density, electric infrastructure, croplands, pasturelands, roads, railways, and navigable waterways (Venter et al., 2016). Higher values represent greater intensity of human landscape modification, providing a proxy for habitat alteration and anthropogenic disturbance. Next, nighttime lights data are extracted as an additional proxy for urbanization from the Defense Meteorological Satellite Program – Operational Linescan System sensor (eogdata.mines.edu/products/dmsp/#rad_cal). The F18 2013 v4c intercalibrated stable lights composite is used (annual 2013 average visible radiance cleaned of ephemeral sources such as fires, gas flares, and lightning). This dataset captures spatial patterns of urbanization, infrastructure, and anthropogenic activity as reflected by persistent artificial light at night (Baugh et al., 2010; Elvidge et al., 1997). The global raster is accessed via Zenodo (doi.org/10.5281/zenodo.17416838).

viii. **Burned area variable.** Global burned area (*env.datasets = "burned_area"*) is extracted using the European Space Agency Fire Disturbance Climate Change Initiative Sentinel-3 SYN Burned Area Grid Product version 1.1 derived from the NERC EDS Centre for Environmental Data Analysis platform (catalogue.ceda.ac.uk/uuid/da8e669a74334c82a56e0b470bc4ef04). This dataset provides monthly burned-area estimates in square kilometers at 900-arcsecond spatial resolution (corresponding to approximately 25 kilometers at the equator) derived from Sentinel-3 Ocean and Land Colour Instrument and Sea and Land Surface Temperature Radiometer imagery using a hybrid spectral-temporal change-detection algorithm. To generate a consistent multi-year climatology suitable for niche modeling, the mean burned area is computed per calendar month across 2019–2024. Each month's average raster represents the mean surface area burned per grid cell during that month over the 6-year period. The twelve resulting global rasters are archived together as a zip file and are available via Zenodo (doi.org/10.5281/zenodo.17487468). These layers capture broad-scale spatiotemporal patterns in fire frequency and intensity, providing an indicator of disturbance regimes and vegetation turnover potentially influencing species distributions and habitat dynamics (Andela et al., 2017; Chuvieco et al., 2016).

ix. **Custom environmental rasters.** Additional user-defined environmental rasters can be incorporated to extend or replace the implemented default variable set (custom.env.rasters argument). Inputs may be supplied as local file paths or URLs and can be a single GeoTIFF (.tif/.tiff), a directory containing one or more GeoTIFF files, or a compressed archive (.zip) containing one or more GeoTIFF files. Each input is processed sequentially. The argument custom.env.rasters.names specifies short descriptive identifiers (e.g., "soil_moisture") that are used to name the resulting folders and output files. The argument *custom.env.rasters.variable.names* provides variable names for individual raster layers (e.g., "NDVI_median", "NDVI_min"). If no variable names are provided, the function automatically generates sequential names based on the raster's layer count (e.g., "EVI_1", "EVI_2", …). Multiple rasters can be provided simultaneously by supplying vectors of equal length to these arguments. Each position across the vectors (*custom.env.rasters*, *custom.env.rasters.names*, *custom.env.rasters.crs*) is treated as one complete dataset, allowing several independent rasters or raster stacks to be processed in a single run. The workflow supports flexible naming conventions and multi-layer rasters (assigning sequential variable suffixes when layer-specific names are not provided). This modular approach enables seamless integration of external predictors. A minimum file size threshold in MB can be set to prevent incomplete downloads (*custom.raster.min.size.mb* argument). It is recommended to set a valid CRS (*custom.env.rasters.crs* argument) only when needed: the function reads the CRS directly from the custom raster itself, whereas a user-supplied CRS is required if the raster lacks embedded CRS metadata.

6. **Output.** The function returns a combined data frame containing occurrence identifiers, coordinates, and all extracted environmental variables, written to the *output.dir* as a CSV file (*csv.occurrence.out.file* argument). If background data were generated, a separate CSV (*csv.background.out.file* argument) is produced, containing environmental values for random background points.

**M2: Pre-DAPC preprocessing pipeline**

Before applying our DAPC niche divergence test, we provide a standardized seven-step preprocessing pipeline to harmonize occurrence and background data and reduce known sources of bias in niche divergence testing (N. Barve et al., 2011; Broennimann et al., 2012; Brown & Carnaval, 2019; Dormann et al., 2007; Elith & Leathwick, 2009; Guisan et al., 2014; Kramer-Schadt et al., 2013; Mesgaran et al., 2014; Phillips et al., 2009). All package functions and their dependencies are summarized in Supplementary Table S1. Below, we describe our preprocessing pipeline in detail:

1. **Prepare data.** Occurrence records and background points are imported and harmonized. If integer columns are present, they are transformed to numeric (using our *convert.integer.to.numeric* function).

2. **Accessible background.** For each species, the accessible area M (sensu Soberon & Peterson, 2005) is approximated by buffering a convex hull polygon around occurrences (via our *crop.background.buffered* function). The buffer distance is ideally in reference to the inferred dispersal capacity of focal species, with more mobile species requiring larger buffer distances (N. Barve et al., 2011). Coordinates are converted to simple features using the *st_as_sf* function in *sf*. If the input CRS is geographic, coordinate values are checked against valid longitude and

latitude ranges, and the data are projected into a metric CRS so that distances are computed in meters. By default, the function heuristically chooses between a local Universal Transverse Mercator zone for compact extents (centroid-based zone; EPSG 326## in the Northern Hemisphere or 327## in the Southern, constrained to UTM zones 1–60) and a locally centered Lambert Azimuthal Equal-Area projection for large, multi-zone, or polar-proximate extents. If the input coordinates are already supplied in a projected CRS with meter units, geometric operations are performed directly in that CRS. Datasets spanning more than 180 degrees of longitude are flagged because antimeridian-crossing ranges may require manual CRS handling. These steps ensure that buffering and spatial intersections are performed in distance-preserving meter units while retaining compatibility with both geographic and projected input coordinates (Hijmans, 2012; Snyder, 1987). Base geometry is constructed as a convex hull by uniting points with the *st_union* function in *sf* and wrapping them with the *st_convex_hull* function in *sf*, then buffered by the user-supplied distance with the *st_buffer* function in *sf*. We set the convex hull (*buffer.method = "hull"*) as the default geometry because it provides a robust and biologically meaningful approximation of the accessible area (N. Barve et al., 2011; Owens et al., 2013; Soberón & Peterson, 2005). The convex hull traces the outer extent of occurrences and provides a realistic approximation of the accessible area while avoiding overinflated areas and unnecessary parameter dependence, especially when buffered by an ecologically informed dispersal distance. In addition to the convex hull, three alternative geometries are available in our function: concave hulls (*buffer.method = "alpha"*), per-point buffers (*buffer.method = "points"*), and bounding boxes (*buffer.method = "bbox"*). Concave alpha hulls are a viable alternative that follow the detailed shape of distributions and may approximate complex ranges more accurately, but are sensitive to the concavity parameter α (Edelsbrunner & Mücke, 1994; Fourcade, 2016; Pateiro-López & Rodríguez-Casal, 2010). Lower α values yield more concave outlines that closely fit the points but risk creating holes and fragmentation, whereas higher values produce smoother outlines that approach the convex hull. We set $\alpha = 3$ as the default (*alpha* argument) as a robust compromise capturing range complexity while avoiding over-fragmentation (Fourcade, 2016; Pateiro-López & Rodríguez-Casal, 2010). Concave hulls are built with the *concaveman* function in the *concaveman v.1.1.0 R* package (Gombin et al., 2020), with an internal safeguard requiring at least three unique coordinates to ensure valid geometry. Per-point buffers generate local circles around each record and capture fragmented clusters but can produce patchy outlines (R. P. Anderson & Raza, 2010; Fourcade et al., 2014). Per-point buffering is performed by applying the *st_buffer* function in *sf* to each point and uniting buffers with the *st_union* function in *sf*. Although realistic for disjunct or fragmented distributions, this method is only feasible for dense background data and large species distributions. Bounding boxes provide the most inclusive geometry by enclosing all occurrences within a rectangle defined by the minimum and maximum coordinates (Peterson et al., 2007, 2011; Phillips et al., 2006; VanDerWal et al., 2009). Because bounding boxes represent the least realistic approximation and often greatly overestimate accessible environments (VanDerWal et al., 2009), we recommend avoiding bounding boxes except in cases of small species distributions where the other methods cannot retain sufficient sample sizes. Bounding boxes are generated with the *st_bbox* and *st_as_sfc* functions in *sf*, then buffered with the *st_buffer* function in *sf*. After geometry creation, invalid polygons are repaired with the *st_make_valid* function in *sf*, and the background is retained using memory-efficient sparse intersections via the *st_intersects* function in *sf* to retain only background rows inside the buffered accessible area. Finally, background points are down-sampled to a standardized maximum number (N.rows argument; default: 10,000) using our *sample.down* function. If fewer points are available than requested, all available rows are retained. The function probabilistically prioritizes records with fewer missing values (after dropping geometry with the *st_drop_geometry* function in *sf*, when present). For each row, the number of missing values is counted and used to assign Poisson-tail weights. The weight for a row is defined as the probability that a Poisson-distributed random variable with mean lambda takes a value greater than or equal to the number of missing values in that row. Therefore, rows with fewer missing values receive a higher sampling probability. If all weights are non-finite or zero, the function falls back to uniform sampling without replacement. By default, lambda is set to adaptively use the median number of missing values across rows (*poisson.lambda = NULL*), but user-specified values can be provided. Sampling is performed without replacement to ensure unique rows. This approach preserves randomness while favoring more complete records for downstream analyses. Ties are handled stochastically through the weighting scheme rather than deterministic ranking. This is consistent with guidance on balanced background sampling and data quality control (Elith & Leathwick, 2009; Merow et al., 2013).

3. **Presence thinning & autocorrelation check.** A minimum nearest-neighbor distance is enforced among occurrences using our *thin.occurrence* function to mitigate sampling bias and spatial autocorrelation (Aiello-Lammens et al., 2015; Boria et al., 2014; Dormann et al., 2007; Fourcade et al., 2014; Inman et al., 2021; Kramer-Schadt et al., 2013; Lamboley & Fourcade, 2024; Veloz, 2009). For datasets with 5000 or fewer valid coordinate records, pairwise great-circle distances are computed using the *rdist.earth* function (*setting miles = FALSE*) in the *fields v.16.3.1 R* package (Nychka et al., 2021). For datasets with more than 5000 valid coordinate records, we use a sparse distance-neighbour search with the *st_is_within_distance* function in *sf* instead of constructing a full pairwise distance matrix, because the full matrix scales quadratically with sample size. For example, 10,000 records require 100 million entries. This reduces memory use and speeds up thinning for large datasets (by around 30% in our benchmark with 13 thousand records). Nonetheless, runtime can still be substantial when records are highly clustered, the thinning distance is large, or many thinning replicates are requested. Records violating the minimum nearest-neighbor distance are identified from either the full pairwise distance matrix or the sparse neighbor list, depending on sample size, and the most spatially redundant record is iteratively removed until all remaining pairs exceed the threshold. In cases of ties, the record with more missing environmental values is removed first, while any remaining ties are resolved at random. Thinning is repeated for the requested number of replicates (*N.thinning.replicates* argument), and the replicate yielding the largest retained subset of records is selected. By default, we set the thinning threshold (*thinning.dist.km* argument) to one kilometer because it approximates the resolution of many fine-scale

environmental GIS layers that are commonly used in ecological niche modeling. However, no single thinning distance is universally recommended in niche modeling, so the appropriate value depends on predictor resolution, the spatial structure of sampling bias, and the biology and geographic extent of the study system (Aiello-Lammens et al., 2015; Boria et al., 2014; Fourcade et al., 2014; Inman et al., 2021; Kramer-Schadt et al., 2013; Lamboley & Fourcade, 2024). Therefore, users should increase this parameter when occurrence clustering reflects broader-scale sampling bias, or decrease it for narrowly distributed taxa, fine-resolution predictors, or sparse datasets where stronger thinning would remove too many records (Aiello-Lammens et al., 2015; Boria et al., 2014; Kramer-Schadt et al., 2013; Veloz, 2009). Residual autocorrelation in environmental predictors is quantified using Moran's I (Moran, 1950) via the *moran.test* function in the *spdep R* package *v.1.3.13* (Bivand, 2022; Bivand et al., 2013; Bivand & Wong, 2018; Pebesma & Bivand, 2023). Spatial weights are constructed using projected metric coordinates (Universal Transverse Mercator zones selected automatically from the dataset centroid) by progressively increasing the number of nearest neighbors until the graph becomes connected. The maximum number of nearest neighbors is capped at one-third of the sample size to avoid overly dense graphs and suppress connectivity warnings in small datasets. Neighbors are obtained with the *knearneig*h and *knn2nb* functions (symmetrized) in *spdep*, and connectivity is verified with the *n.comp.nb* function in *spdep*. The resulting graph is converted to a row-standardized *listw* object with the *nb2listw* function in *spdep* (*setting style = "W", zero.policy = TRUE*) (Bivand et al., 2013; Dormann et al., 2007). Latitude and longitude columns are automatically excluded from autocorrelation testing. Results are summarized by reporting the median Moran's I and visualized as histograms of its distribution across variables using the *hist* function in the *base R v.4.4.1* package (R Core Team, 2025). When sample sizes allow, we suggest thinning data until Moran's I values fall below 0.5, corresponding to moderate spatial autocorrelation (Fortin & Dale, 2005; Legendre, 1993). Although this cutoff is conservative, it ensures datasets remain tractable while reducing residual spatial dependence to biologically acceptable levels (Dormann et al., 2007).

4. **Equal sample sizes.** After thinning, the two occurrence datasets are down-sampled to equal sample sizes via our *sample.down* function using the same NA-aware ranking as in step 2. This avoids unequal sample sizes that can bias discriminant functions toward the species with more samples and inflate apparent divergence because unequal group sizes change the pooled within-group covariance matrix and classification thresholds in discriminant analysis (Lachenbruch, 1975; Lachenbruch & Goldstein, 1979; McLachlan, 1992).

5. **Transformation of skewed variables.** Skewed variables are transformed to stabilize variance and reduce the influence of extreme values. Otherwise, strongly right- or left-skewed predictors can dominate the first few PCA (principal component analysis) axes and bias discriminant functions toward variables with heavy tails or restricted ranges (Bartlett, 1947; Box & Cox, 1964; Osborne, 2010). This is done via our *transform.skewed.variables* function, which computes unbiased Fisher–Pearson sample skewness for each numeric predictor after ignoring user-specified non-environmental columns (*exclude.cols* argument). Fisher–Pearson skewness

measures the asymmetry of a distribution by comparing the relative weight of its left and right tails (Fisher, 1930; Joanes & Gill, 1998). A value near zero indicates approximate symmetry, positive values indicate a long right tail, and negative values indicate a long left tail. Variables with absolute skewness higher than a threshold (*skewness.threshold* argument) are evaluated against a candidate set of transformations, and the transformation is chosen by comparing the reduction in absolute skewness relative to the untransformed variable. If no candidate reduces absolute skewness, the variable is left unchanged. We set 1.0 as the default for absolute skewness, indicating substantial asymmetry (Bulmer, 1979; Doane & Seward, 2011). Candidate transformations are generated according to the observed data range and type of variable; when background data are supplied, the occurrence and background values are evaluated jointly to ensure that both datasets are transformed on the same scale. Continuous proportion variables bounded between zero and one are evaluated using a shrinkage logit transformation (rescales proportions while preserving mid-range differences) and an arcsine square-root transformation (stabilizes variance when data are concentrated near the boundaries or include many zeros) (Warton & Hui, 2011). The option producing the larger reduction in absolute skewness is selected. For other strictly positive variables, natural-log and square-root transformations are evaluated. Both reduce right-skewed distributions but differ in strength, with the log transformation preferred for long right tails (Bartlett, 1947; Box & Cox, 1964; Emerson & Stoto, 1983). For non-negative variables that include zeros, a square-root transformation and a natural-log transformation after adding one to all values are evaluated (Cleveland, 1984; Emerson & Stoto, 1983). Variables containing negative values are transformed using approaches that preserve both the sign and relative order of values. In these cases, the data are first shifted so that the minimum value becomes slightly greater than zero, allowing logarithmic and square-root transformations to be applied safely (John & Draper, 1980; Manly, 1976; Tukey, 1977). A signed cube-root transformation is also tested, which can handle both positive and negative values directly while maintaining proportional scaling across the range (John & Draper, 1980; Manly, 1976; Tukey, 1977). For strongly right-skewed positive variables (skewness higher than three), the natural-log transformation is preferred directly as a special-case rule for heavy right tails. Variables with fewer than three unique finite values, binary 0/1 variables, and other two-level variables are not transformed. For each variable, the function reports the skewness before and after transformation, the transformation method applied, and the selection rationale, producing a detailed summary table. To keep scaling consistent between occurrence and background data, the same transformations are automatically applied to the background dataset (*background.dataframe* argument), ensuring that both datasets share a uniform set of transformed environmental variables. Transformed variables are labeled with suffixes indicating the applied transformation (replacing the original column in both datasets), while untransformed variables retain their original names.

6. **Low-information filter:** Predictors with no or low variability are removed based on the coefficient of variation (Pearson, 1896) in either species' occurrences using our function *remove.low.CV.vars*. These predictors provide negligible discriminatory power because they contribute little to between-group separation in discriminant analysis. Including such near-

constant variables increases the dimensionality of the dataset without adding information, increases the condition number of the covariance matrix, and can lead to numerical instability or singular matrices during PCA or discriminant analysis (Dormann et al., 2013; Greenacre & Primicerio, 2014). Variables with fewer than five finite observations per species are also excluded because such sparse data cannot produce a reliable coefficient of variation estimate. After dropping geometry with the *st_drop_geometry* function in *sf* where needed, numeric columns in each occurrence table are identified, user-specified non-environmental fields are excluded, and the coefficient of variation is computed as the standard deviation of a variable divided by the absolute value of its mean (both calculated after removing missing values) (Pearson, 1896; Sokal & Rohlf, 2012; Zar, 2010). We prefer the coefficient of variation over variance because it is unitless and scale-invariant, making it directly comparable across environmental predictors measured in different units and magnitudes. In contrast, variance is unit-bearing and depends on measurement scale, so variables with larger numeric units appear more variable even when their relative dispersion is negligible. Therefore, using the coefficient of variation flags predictors that are effectively constant relative to their typical level, independent of unit choice and prior to any standardization, which improves numerical conditioning in downstream multivariate analyses (Pearson, 1896; Sokal & Rohlf, 2012; Zar, 2010). We implement a coefficient of variation threshold of 0.01 as the default (*CV.threshold* argument) corresponding to variables with less than 1% relative variability compared to their mean, meaning that they are effectively constant across occurrences. The same variables are also excluded from the backgrounds to ensure synchronized inputs across species. For variables with means close to zero, where the coefficient of variation is unstable or undefined, the relative variability is calculated instead using a scale-free standard deviation–to–median absolute deviation ratio to avoid spurious inflation (Maronna et al., 2006; Zar, 2010). This fallback measure is evaluated against the same threshold used for the coefficient of variation.

7. **Environmental analogy screening.** To reduce bias from non-analogous environments (Brown & Carnaval, 2019; Guisan et al., 2014), we remove variables with insufficient univariate and bivariate overlap between species' accessible background spaces using our *filter.analogous.variables* function. This function first discards low-information predictors, then quantifies environmental overlap using univariate kernel-density estimates and bivariate probability grids, retaining only variables that are sufficiently variable and environmentally comparable between the two taxa. First, variables with fewer than the minimum required complete observations per species (*min.rows* argument; default: *15*) are removed, ensuring stable estimates of variation and overlap. Background variables with low variability are removed based on a default 0.01 CV threshold (*CV.threshold* argument; similar to the low-information filter in step 6). Univariate overlap is then quantified using kernel-density estimates of Schoener's D via the *density* function in the *base R stats v.4.4.1* package (R Core Team, 2025), retaining variables above the specified overlap threshold (*overlap.threshold* argument; see below). Bandwidths are estimated from pooled values of both species using the *bw.SJ* function in *base R stats* (*method = "dpi"*), falling back to the *bw.nrd0* function if needed. This ensures consistent kernel smoothing across species without bias from unequal sample

sizes. After this univariate overlap filtering, bivariate overlap is assessed among the remaining variable pairs using quantile-binned two-dimensional histograms. Each pair's overlap is calculated as the sum of the element-wise minima between the two normalized joint-probability grids, equivalent to a two-dimensional extension of Schoener's D. Variables that do not appear in any high-overlap pairs (overlap ≥ threshold) are considered non-analogous and removed. To reduce bias from unequal sample sizes, the quantile breaks are determined from the pooled background values of both species, which equalizes occupancy across bins and stabilizes estimates under skewed distributions (Mesgaran et al., 2014; Scott, 2015; Silverman, 1986). The bin number is chosen automatically as the square root of the effective sample size divided by twenty, rounded down to the nearest whole number, and clamped to 5–20 bins, to ensure resolution without sparse-bin noise. Before overlap calculations, non-finite values are treated as missing, and background rows with a missing-value proportion greater than *max.NA.prop* are removed. Remaining missing values are then either imputed by the variable median when *impute.NA.median = TRUE* or removed by complete-case filtering when *impute.NA.median = FALSE*. Median imputation is used for these overlap calculations when the within-variable missing value proportion ≤ 0.2 (*max.NA.prop* argument). We use a default threshold of 0.2 consistent with general recommendations for minimizing imputation bias (Harrell, 2015; van Buuren, 2018). Environmental variables exceeding this threshold are removed. Variables with few complete observations per group are excluded (*min.rows* argument; default: *15*), ensuring kernel densities and histogram estimates are based on sufficient data for stability (Silverman, 1986). Because the number of pairs can be large, a coverage-aware sampler is employed to ensure each predictor participates in multiple pairs (*max.pairs* argument). Parallelized computation can be used (if *use.parallel = TRUE*) with *N* cores (*N.cores* argument) to significantly speed up bivariate overlap computation using the *makeCluste*r, *parSapply*, *clusterExport*, and *stopCluster* functions in the *parallel v.4.4.1 R* package (R Core Team, 2025). The *clusterSetRNGStream* function in *parallel* is used to ensure reproducible parallel RNG streams. For both univariate and bivariate overlap filtering, a commonly used value of 0.7 is implemented as the default overlap threshold (*overlap.threshold* argument): values below indicate poor analogy and high risk of extrapolation, while stricter thresholds may eliminate informative predictors (N. Barve et al., 2011; Peterson et al., 2011; Rödder & Engler, 2011; Warren et al., 2008). We restrict screening to univariate and bivariate projections rather than higher-dimensional spaces because density estimation in higher dimensions demands exponentially larger sample sizes and rapidly becomes computationally infeasible (Scott, 2015; Silverman, 1986). Most extrapolation risk is already detectable in marginal or pairwise dimensions (Mesgaran et al., 2014; Peterson et al., 2011), so univariate and bivariate checks provide an efficient compromise between computational tractability and effective bias detection (Scott, 2015; Silverman, 1986).

The resulting occurrence matrix produced by our pipeline constitutes the input to the main DAPC niche divergence test. Unlike ENM-based niche divergence tests (Warren et al., 2008, 2010), our divergence test is conducted entirely in environmental space (E-space) rather than geographic space (G-space). This helps reduce G-space bias, where niche similarity estimates are confounded by the geographic co-occurrence of taxa and biased toward common habitats (Brown & Carnaval, 2019; Di Cola et al., 2017). By framing divergence tests explicitly within accessible background environments, restricting comparisons to analogous environmental conditions, and incorporating seasonal and monthly predictors (see Empirical test: Supplementary Methods M7–M9), we reduce bias from non-equilibrium distributions, where realized occurrences capture only a subset of the tolerable niche (Brown & Carnaval, 2019; Peterson et al., 2011; Soberón, 2007). Together, these steps provide a standardized pipeline for robust divergence estimation that reduces the main sources of bias in niche divergence testing: sampling bias, G-space bias, non-analogy, and non-equilibrium.

In addition to our Environmental Analogy Screening (step 7; *filter.analogous.variables*), which removes non-analogous environmental predictors based on univariate and bivariate overlap thresholds, we also implement a complementary Humboldt-style analogous trimming to restrict occurrence records to environmentally shared regions between species for comparison and methodological consistency. This trimming follows the *Humboldt*'s *humboldt.g2e* function (Brown & Carnaval, 2019), enabling parallel evaluation of both predictor- and occurrence-based analogy corrections within our framework. To do so, our *trim.to.analogous.environments* function projects both species' background datasets into a shared environmental PCA space (E-space) using the *dudi.pca* function in the *ade4 v.1.7.23 R* package (Bougeard & Dray, 2018; Chessel et al., 2004; Dray & Dufour, 2007; Thioulouse et al., 2018), based on the intersection of shared numeric environmental variables. Similar to step 6, low-information predictors with excessive missing data or near-zero variance are removed, and remaining values are optionally imputed by the column median if their missing proportion does not exceed 0.2 (*max.NA.prop* argument). The combined background matrix of both species is then used to derive PC1 and PC2 (principal components) that define the shared environmental space. Following *Humboldt*'s *nae.window* trimming logic, each species' background is iteratively clipped within a moving window (*analogous.window.size* argument; default: *5*) along a PCA grid defined by *grid.resolution = 100*. For every band along PC1, species 2's background is restricted to the PC2 range occupied by species 1 within that band, extended on both sides by the window size. The process is then repeated reciprocally along PC2, trimming species 1 relative to species 2, followed by a final reciprocal pass (species 1 then species 2 then species 1) to ensure symmetry of the retained analogous space. This bidirectional filtering ensures that both species are represented only in environmentally analogous regions of PCA space, equivalent to the reciprocal reduction trimming when *REDUC = 5* in *Humboldt's humboldt.g2e* function. After the reciprocal trimming, the function defines the final shared environmental extent by computing the intersection of the PC1–PC2 ranges occupied by both species. Occurrence records for each species are projected as supplementary points into this same PCA model using the *suprow* function in *ade4*, and retained only if their coordinates fall within the shared environmental bounds. This results in a pair of trimmed occurrence datasets restricted to environmentally comparable regions, eliminating extrapolated or non-analogous occurrences that would otherwise bias downstream niche-divergence analyses.

##### M3: DAPC permutation procedure

Our DAPC method identifies a single discriminant axis that maximizes between-group variation relative to within-group variation, summarizing multivariate differences between groups in one-dimensional space (see Methods). However, the statistical significance of this separation cannot be inferred directly from this discriminant function. Group differentiation may arise by chance, especially in high-dimensional data. To test whether the observed group separation exceeds random expectation, we implement a permutation procedure that generates a null distribution of group assignment under the hypothesis of a single shared niche (k = 1), effectively emulating one shared environmental space.

First, the PCA space of the environmental variables for all occurrence records is computed with centered and scaled data using the *dudi.pca* function in *ade4*. The number of retained PCs is fixed to the cross-validated optimum or optionally to a user-specified override. Group labels are reshuffled globally without replacement while preserving overall group sample sizes. Discriminant analysis is re-fit on the fixed PC scores using the *lda* function in the *MASS v.7.3.65 R* package (Venables & Ripley, 2002) with empirical class priors computed from the labels used in each fit (observed or permuted). The overall assignment accuracy (proportion correctly classified) is recomputed for every replicate. Repeating this procedure many times (*N.permutations* argument; default 1,000) yields a null distribution of assignment accuracy. One-sided p-values are calculated via a right-tailed test by comparing observed statistics to their null distributions. To avoid p-values equal to zero when the observed statistic is more extreme than all permutation replicates, a standard +1 correction is applied to the numerator and denominator (North et al., 2002; Phipson & Smyth, 2010). Numerical stability of the discriminant analysis is controlled via the *tol* parameter (tol = $1e^{-8}$) in the *lda* function. Failed permutation replicates due to singular covariance are discarded, and the p-value denominator reflects the number of successful permutations.

Random number generation uses L'Ecuyer–CMRG streams (L'Ecuyer, 1999) via the *RNGkind("L'Ecuyer-CMRG")* function in *parallel*, ensuring reproducibility across serial and parallel executions. Parallelization is supported through *parallel*: on Unix systems via the *mclapply* function, and on Windows via PSOCK clusters created with the *makeCluster* function and distributed using the *parLapplyLB* function. To ensure reproducible parallel RNG, we set a seed and, for PSOCK clusters, initialize substreams with the *clusterSetRNGStream* function in *base R*. In practice, parallelization typically accelerates the permutation procedure by a factor of roughly five to fifteen, depending on dataset size and the number of cores used. However, due to overhead associated with PSOCK clusters and the relatively light computational load of individual permutation iterations, optimal performance is generally achieved with a modest number of workers (2–4 cores). Using larger numbers of cores often provides diminishing or negative returns because the communication overhead dominates the per-iteration compute time.

**M4: Niche divergence metrics and visualization**

For pairwise comparisons, DAPC returns a single discriminant function that collapses the multivariate niche into a one-dimensional discriminant axis (see Methods for details). This axis represents the linear combination of predictors that maximizes between-group separation relative to within-group variation, summarizing the multivariate niche structure into a single interpretable dimension. The resulting discriminant scores for each taxon correspond to their positions along this univariate axis, allowing visualization and quantification of niche overlap, exclusivity, and divergence. To extract, evaluate, and visualize these DAPC results, we implemented several functions (Supplementary Table S1) in our package:

- Our *plot.DAPC.niche.divergence* function produces kernel density estimates of the DAPC discriminant scores for the two taxa on a shared axis, with individual scores visualized as rug lines below the axis (e.g., Fig. 4A, Supplementary Figs. S12, S18). This provides a clear view of multivariate niche overlap in one dimension.

- Our *plot.DAPC.permutation function* visualizes the permutation test results by displaying a histogram of a null distribution for assignment accuracy (e.g., Fig. 4B, Supplementary Figs. S12, S18). Observed values are added as vertical red lines. This visualization shows whether observed separation exceeds chance expectations under permutations simulating a single shared niche.

- Our *calc.niche.divergence.metrics* function provides five metrics that quantify and characterize niche divergence. To do so, it estimates kernel densities for both taxa on a shared grid (default resolution: 1024; *density.grid.resolution* argument) using a pooled bandwidth to ensure comparability. Densities are normalized to integrate to one and max-normalized by dividing through their peak value, following equation 1 in Ascanio et al. (2024). This max-normalization ensures that each group's density peaks are on the same scale, allowing proportional comparisons of divergence. When *weight.background = TRUE*, a Humboldt-like background correction (Brown & Carnaval, 2019) is applied to reduce bias from unequal frequencies of available environments along the discriminant axis by up-weighting rare and down-weighting common environments. Each species' background dataset is projected into discriminant space using the *dudi.pca* function in *ade4* to obtain PCA loadings, the *lda* function in *MASS* to extract discriminant scaling vectors, and the *scale* function in *base* to standardize variables. These components are combined through matrix multiplication to compute background LD1 scores. Kernel densities for both background and occurrence LD1 values are estimated using the *density* function in *stats* and averaged to represent the combined available environmental space. The function then up-weights rare environments and down-weights common ones through weighted density scaling, followed by normalization of the corrected densities. This correction reproduces the *humboldt.espace.correction* function (*correct.env = TRUE*) in the *humboldt* package (Brown & Carnaval, 2019) in DAPC-derived discriminant space. As the first metric, the function calculates Schoener's D (Schoener, 1968), which measures the proportion of shared density from zero (no overlap) to one (complete overlap). Four additional metrics are based on the niche divergence plane of Ascanio et al. (2024). We extend their framework from single environmental variables to the multivariate discriminant axis returned by DAPC. Niche dissimilarity (NDS) is based on one-sided dissimilarities, where for each taxon the proportion of its normalized density that does not overlap with the other is obtained by dividing the non-overlapping area of the density by the total area under its curve across its support (equation 3 of Ascanio et al. 2024). The two one-sided values are then averaged to yield the final NDS, which represents the mean fraction of density unique to each taxon (equation 2 of Ascanio et al. 2024). Niche breadth exclusivity (NE) is quantified by first identifying the lower and upper bounds of the discriminant axis where each taxon's normalized density exceeds a minimal support threshold of $1e^{-6}$. The shared portion of these ranges is divided by the total union, and exclusivity is then defined as one minus this shared proportion (equation 4 in Ascanio et al. 2024). Niche divergence magnitude (ND) combines the two previous metrics by calculating the square root of the sum of squared niche dissimilarity and niche exclusivity values (equation 5 in Ascanio et al. 2024). In other words, the niche divergence magnitude increases as either dissimilarity or exclusivity rises, providing a single index that summarizes the overall strength of niche divergence. Niche divergence angle (θ) is calculated as the arctangent of niche dissimilarity relative to niche exclusivity, expressed in degrees (equation 6 in Ascanio et al. 2024). Angles near zero degrees indicate that divergence is explained mainly by differences in range limits (i.e., exclusivity), angles near ninety degrees indicate divergence driven mainly by differences in density within shared space, and intermediate values reflect mixed contributions of both.

- Our *plot.DAPC.var.contributions* function identifies and displays the potential ecological drivers of divergence by back-transforming the discriminant axis into the original predictor space (e.g., Fig. 5A). Each variable receives a loading that reflects its weight, which is squared and normalized to sum to one. The direction of effect is indicated by the sign of the loading relative to group centroids. The function produces a bar plot colored by the group with higher values. Optional arguments allow limiting the plot to the top predictors (*top.N* argument) or excluding those below a contribution threshold (*min.contribution.threshold* argument).

- Our *plot.top.DAPC.predictors* function visualizes univariate density distributions of top environmental variables (e.g., Fig. 5B). Variables are ranked by their contribution values and displayed as kernel density plots for each taxon. This approach links the discriminant axis separation back to familiar ecological predictors and highlights whether divergence reflects shifts in means, differences in spread, or separation in distribution tails.

- Our *plot.occurrences.map* function visualizes the geographic basis of divergence analyses (i.e., G-space or geographic space) by displaying the occurrence and optionally background points on a political basemap (e.g., Fig. 2). The function expects coordinates in decimal degrees. Map extents are buffered. For the United States, optional state and county boundaries can be added (*add.USA.states* and *add.USA.counties* arguments). The function relies on the *maps v.3.4.3 R* package (Becker et al., 2025) for basemaps and *base R graphics* (R Core Team, 2025) for rendering.

All plotting functions allow users to adjust key parameters such as line widths, colors, text, and legend placement. By default, group colors are dark blue and grey (e.g., Supplementary Fig. S13), chosen for high contrast on white backgrounds, robustness to common color-vision deficiencies, clarity in grayscale reproduction, and a neutral tone that reduces semantic bias while keeping densities and rugs distinguishable under transparency. Non-map plots use the *ggplot2 v.4 R* package (Wickham, 2016) and can be exported via the *ggsave* function as PNG, JPG, or SVG with user-defined size and resolution. Maps are saved via *base R graphics* devices as SVG, PNG, or JPG.

#### II) Method validation and benchmarking (simulations and comparisons)

##### M5: Simulations of multivariate niches

To evaluate the robustness and accuracy of our DAPC niche divergence framework, we simulated virtual species under controlled conditions to create a gradient of ecological divergence scenarios. Virtual species are simulated entities that provide a rigorous way to test the statistical behavior of new ecological methods such as our niche divergence test under controlled occurrence–environment relationships and sampling strategies (Leroy et al., 2016; Meynard et al., 2019; Zurell et al., 2010). Hence, these simulations allowed us to assess whether the test correctly detects true divergence, maintains an appropriate type I error rate, and performs consistently across varying degrees of environmental overlap. They also allowed verification that the method correctly infers no niche divergence when both taxa share the same simulated niche, ensuring that apparent separation does not arise spuriously from sampling noise or model structure.

Synthetic species were generated using the *virtualspecies v.1.6 R* package (Leroy et al., 2016) following general workflow recommendations (Leroy, 2018; Leroy et al., 2016; Meynard et al., 2019). This was done by generating virtual suitability rasters from our empirical background data (see Empirical test: Supplementary Methods M7–M9), followed by sampling presence-only occurrences to generate species distributions, and extracting environmental values at those locations. Synthetic environmental suitability surfaces were generated along the first sixteen PC axes of variation (explaining 86.7% of the total variance) derived from the background data, representing North America. Our aim was to first establish a null model of species and then simulate niches that increasingly diverge stepwise in multivariate suitability space from this baseline. This allowed us to test how our method and alternative approaches detect increasing levels of multivariate niche differentiation.

All numeric background variables were rasterized at 30-arcsecond spatial resolution within the convex hull of all background points buffered by 80 kilometers to avoid edge cuttings using the *rasterize* function (*fun = mean*) in *terra*. All environmental rasters were combined into a SpatRaster stack. 500,000 random raster cell values were sampled from the stacked layers via repeated random cell-ID sampling using the *sample* function in *base R*, restricted to eligible raster cells identified by computing a fraction-valid raster with the *app* function in *terra* and retaining only those where at least 70% of layers contained valid (non-NA) values. This was done in 20,000-cell chunks to overcome memory limitations. Conversion to coordinates and value extraction were done with the *xyFromCell* and *extract* functions in *terra*, respectively. Samples with more than 30% missing data were discarded, and the remaining missing values were imputed with the mean of each variable. Columns with zero variance were dropped. A PCA was performed with sixteen axes on the centered and scaled sampled matrix using the *dudi.pca* function (*center = TRUE, scale = TRUE, nf = 16*) in *ade4*. For downstream raster computations, each raster cell was projected into PCA space using a custom wrapper around the *predict* function in *terra*, which applied the PCA centering, scaling, and loading-matrix rotation directly to raster cell values, with column-wise mean imputation for missing values. This projection was computed once (rather than separately for each simulation as done in *virtualspecies*). Species-specific suitability layers were then computed by applying a custom Gaussian wrapper to the saved PCA scores using the *app* function in *terra*.

**Simulation set 1.** We defined Gaussian response functions along the first two PCs (explaining 60.8% of the total variance together), with fixed standard deviations (PC1: 5, PC2: 7, all other PCs: 4). The null species *Sp00* was placed at the lower end of the PC1–PC2 space. To avoid edge effects, species means were kept at least two standard deviations inside the PCA range, ensuring that nearly the entire niche distribution (95.4%) remained within the considered environmental space. We created 100 simulated species (Sp01–Sp100) with increasingly divergent niches by incrementally increasing the distance between the species' means in PC1–PC2 space. Specifically, we built two sequences of length 100 for PC1 and PC2: species *i* used the *i*-th mean from each sequence, so that means move in lock-step across PC1 and PC2 (PC1 step size: 0.61, PC2 step size: 0.10; PC1 admissible mean range: –19.24 to 41.48, PC2 admissible mean range: –6.28 to 3.97). For each step, a suitability raster was created and min-max rescaled to 0 and 1. This computation reused the single PCA projection and applied the Gaussian wrapper with species-specific means and shared standard deviations via the *app* function in *terra*.

**Simulation set 2.** This set-up was repeated in PC7–PC8 space instead of PC1–PC2 (explaining 3.4% of the total variance together), generating more subtle divergence along multivariate axes that explain approximately 18 times less environmental variation compared to simulation set 1. Gaussian response functions were used with fixed standard deviations (PC7: 31, PC8: 18, all other PCs: 6). Divergence was introduced by shifting the species' means in increments of 1.64 along PC7 in a descending sequence (reversed order), with matched step counts along PC8 in increasing sequence (PC7 admissible mean range: –48.97 to 112.98, PC8 admissible mean range: 0.80 to –15.65, step size PC8: –0.17).

Suitability rasters were converted to probability of occurrence based on a logistic function using the *convertToPA* function in *virtualspecies* (*α = −0.04, prob.method = "logistic"*; for simulation set 1: *β = 0.35*, for set 2: *β = 0.5*). Following recommendations (Meynard et al., 2019), these alpha and beta values were chosen after exploring several settings in prior runs to optimize the probability surface and species distributions. From this probability surface, we drew 4,000 presence-only occurrences with a 90% detection rate and replacement, using the *sampleOccurrences* function in *virtualspecies* (*detection.probability = 0.9, replacement = TRUE*). Only detections (*Observed == 1*) were retained. To create the null species *Sp00*, we duplicated the suitability of *Sp01* representing no shift in mean. Then, we used a different random seed to draw occurrences, yielding identical suitability and environmental niche (i.e., no niche divergence) to Sp01 but independent stochastic occurrence samples. Simulated presence-absence and occurrence records were evaluated to ensure realistic species distributions (Supplementary Figs. S1–S2). Finally, we extracted the environmental values at the sampled coordinates from the original SpatRaster stack for each occurrence file using the *extract* function (*ID = FALSE*) in *terra*.

The *setThreadOptions* function (*numThreads = 1*) in the *RcppParallel v.5.1.11.1 R* package (Allaire et al., 2025) and *terraOptions* function (*memfrac = 0.6, todisk = TRUE*) in *terra* were used for all raster operations. Rasters were written using the *writeRaster* function in *terra* with LZW compression and FLT4S datatype (*wopt = list(datatype = "FLT4S", gdal = "COMPRESS=LZW")*). Intermediate and final products were saved as files to avoid recomputation on subsequent runs.

**M6: Comparisons with other niche divergence methods**

To benchmark our DAPC-based divergence test, we compared it with six alternative niche divergence tests. These comparisons allowed us to assess the relative sensitivity, robustness, and interpretability of our DAPC divergence framework in relation to other commonly used niche divergence approaches. For comparability, all methods were applied to datasets processed through steps 1–7 of our standardized pre-processing pipeline (see Supplementary Methods M2): 1) data preparation and harmonization, 2) delimitation of accessible backgrounds, 3) spatial thinning, 4) equalized sample sizes, 5) transformation of skewed variables, 6) removal of low-information predictors, and 7) environmental analogy screening. Missing values were imputed with the column mean prior to PCA in all alternative niche tests. All analyses were conducted with a fixed random seed for reproducibility. P-values below 0.05 were interpreted as statistically significant across all hypothesis tests, including the DAPC permutation test.

1. ***ecospat PCA-env***. We used environmental PCA with equivalency and similarity tests (Broennimann et al., 2012) as implemented in the *ecospat v.4.1.2 R* package (Broennimann et al., 2025). The group-1 and group-2 background samples were combined into a pooled background matrix, which was mean-centered and scaled, and used to fit the PCA with the *dudi.pca* function in *ade4*. Analyses were limited to PC1–PC2. Group occurrences and group-specific backgrounds were then projected into this PCA space with the *suprow* function in *ade4*. Projected scores were clipped to the PC1–PC2 extent of the pooled background to avoid extrapolation. The pooled background was randomly down-sampled to a maximum of 10,000 rows. Within this common PCA space, smoothed occurrence densities were estimated on a regular grid using the *ecospat.grid.clim.dyn* function with a grid size *R* of 100 and presence threshold *th.sp* of 0.01. The package's default Gaussian kernel smoothing was applied. We built one grid for group 1 using the pooled background as *glob*, the group-1 background as *glob1*, and its occurrence scores as *sp*. A second analogous grid was created for group 2. Niche overlap was quantified with the *ecospat.niche.overlap* function (*cor = TRUE*) using Schoener's D (Schoener, 1968), defined as one minus one-half of the sum, over all grid cells, of the absolute difference between the two normalized density values. This index ranges from zero (no overlap) to one (complete overlap). We tested for niche divergence with the equivalency test using the *ecospat.niche.equivalency.test* function (*rep = 1000, intersection = 0, overlap.alternative = "lower", expansion.alternative = "lower", stability.alternative = "higher", unfilling.alternative = "lower"*). In this test, occurrences from both species were pooled and randomly reassigned to species labels while preserving sample sizes, producing a null distribution of overlap against which the observed overlap is compared. To account for differences in available environments, we also performed background similarity tests in both

directions with the *ecospat.niche.similarity.test* function (*rand.type = 1, rep = 1000, intersection = 0*). The first test randomized occurrences of group 1 within the background of group 2, and the second randomized occurrences of group 2 within the background of group 1. After each randomization, niche overlap was recalculated, producing a null distribution of expected overlaps given the available environment. The observed overlap was then compared to this distribution to evaluate whether the niches are more different than expected by chance. Running the test in both directions evaluates whether the observed overlap is lower than expected given each group's accessible environment. The niche similarity (background) test and the niche dynamics were not computed for the simulations due to computational restraints. Runtime was recorded from PCA through density estimation, overlap computation, and permutation tests. Finally, we visualized test results with permutation histograms for Schoener's D from the equivalency test and both background-similarity directions using *ggplot*.

2. **PERMANOVA.** We performed stratified PERMANOVA (permutational multivariate analysis of variance), which is a distance-based multivariate method (M. J. Anderson, 2001; M. J. Anderson & Walsh, 2013). Environmental predictors were scaled and centered, and then transformed into orthogonal PCs via PCA using the *prcomp* function in *base R*. We retained PCs using the broken-stick criterion (Frontier, 1976), which compares each PC's proportion of explained variance to expectations under a random stick-breaking model, retaining only axes that exceed this baseline. The broken-stick method is a conservative and effective rule for identifying interpretable components in ecological multivariate analysis (D. A. Jackson, 1993; Legendre & Legendre, 2012). By excluding noise-dominated axes, this approach reduces the risk of diluting distance measures and losing statistical power, providing an objective and transparent cutoff prior to stratified PERMANOVA tests. Euclidean distances were computed on the PC scores using the *vegdist* function in the *vegan v.2.7.1 R* package (Oksanen et al., 2025). PERMANOVA partitions the sums of squares of these distances into between-group and within-group components and calculates a pseudo-F statistic as the mean square between species divided by the mean square within species (M. J. Anderson, 2001). PERMANOVA was run using the *adonis2* function (*by = "margin"*) in *vegan*. Significance was assessed by permutations of sample labels constrained within spatial blocks, ensuring that local autocorrelation was preserved (H. H. Wagner & Dray, 2015). This was done as follows. Coordinates were transformed to an appropriate Universal Transverse Mercator zone chosen by the dataset centroid. An effective spatial range was estimated from an empirical variogram of PC1 (Cressie, 1993; Legendre & Fortin, 1989) and searched over rotated (0°, 22.5°, 45°, 67.5°) and jittered rectangular grids to select a block layout with three or more blocks, a median of ten or more points per block, and mean block purity (dominance of the majority group within a block) of 0.7–0.9. Up to ten iterations and ten jitters per angle were evaluated, with block cell size capped at 90% of the shorter study-area extent. The chosen block factor was then supplied to the *how* function in the *permute v.0.9-8 R* package (Simpson, 2025) to generate 1000 blocked permutations. Because PERMANOVA is sensitive to heterogeneity of multivariate dispersion (M. J. Anderson, 2006), we conducted betadisper tests using the *betadisper* function in *vegan* and only interpreted PERMANOVA results when dispersion was not significantly different

between species. Runtime was recorded from PCA and distance calculation through dispersion testing and the blocked PERMANOVA.

3. **Hypervolume.** We estimated kernel hypervolume overlap (Blonder et al., 2014, 2018) using the *hypervolume R v.3.1.6* package (Blonder et al., 2025). Predictors were centered and scaled, and transformed into orthogonal PCs by PCA using the *prcomp* function in *base R*. Following guidance for stable hypervolume estimation (Blonder et al., 2018), we retained a PC-number equal to the floor of the natural logarithm of the sample size. Gaussian kernel-density estimation was performed with a fixed Silverman bandwidth (Silverman, 1986) estimated from the pooled PCA scores using the *estimate_bandwidth* function (*method = "silverman"*). Hypervolumes were constructed with this pooled bandwidth for each taxon using the *hypervolume_gaussian* function. To reduce sensitivity to low-density tails and outliers, hypervolumes were restricted to the 0.95 probability isosurface using the *hypervolume_threshold* function (*quantile.requested = 0.95, quantile.requested.type = "probability"*). Pairwise overlap was computed using the *hypervolume_se*t function, and similarity indices were extracted with the *hypervolume_overlap_statistics* function. We report two standard similarity indices (Mammola, 2019): Jaccard similarity defined as the intersection volume divided by the union volume of the two thresholded hypervolumes (Jaccard, 1901), and Sørensen–Dice similarity defined as two times the intersection volume divided by the sum of the two hypervolume volumes (Dice, 1945; T. Sørensen, 1948). Both indices range from zero (no overlap) to one (identical hypervolumes), with Sørensen–Dice always greater than or equal to Jaccard for a given pair because it weights the intersection more strongly (Mammola, 2019). We also report the fraction of the union unique to each taxon, which corresponds to the proportion of the combined hypervolume occupied exclusively by each group. In addition to overlap indices, we also computed two distance-based metrics (Mammola, 2019) using the *hypervolume_distance* function: centroid distance (the Euclidean distance between hypervolume centroids) and minimum inter-hypervolume distance. The latter is estimated from 2,000 random samples drawn by taking the smallest cross-distance between sampled points from each group's hypervolume. Both metrics quantify how far apart the hypervolumes are in multivariate space, even when overlap is low or absent (Mammola, 2019). Runtime was recorded from PCA through overlap and distance calculations.

4. **MVNH.** We applied the multivariate normal hypervolume framework (MVNH; Lu et al., 2021) using the *MVNH v.0.1.0 R* package (Lu, 2025). Environmental predictors (scaled and centered) were transformed into orthogonal PCs via PCA using the *prcomp* function in *base R*. We retained the minimum number of PCs that explained at least 90% of variance to retain nearly all variation while avoiding overfitting (Jolliffe & Cadima, 2016; Legendre & Legendre, 2012). In this PCA space, the niches of the species were modeled as multivariate normal distributions defined by mean vectors and covariance matrices. Niche size was estimated as the log determinant of the covariance matrix using the *MVNH_det* function (*log = TRUE*). Divergence was partitioned into three components using the *MVNH_dissimilarity* function: 1) Mahalanobis distance (MD), which measures centroid separation and bounded at zero with no fixed upper limit (MD = 0 when centroids coincide, larger MD values indicate increasing separation); 2) determinant ratio

(DR), which compares the log determinants of the covariance matrices as a measure of relative niche breadth (DR = 0 when breadths are equal, larger values indicate greater breadth disparity); and Bhattacharyya distance (BD), which combines centroid separation and breadth differences into a single measure of total divergence (BD ≥ 0, larger values indicate stronger divergence). Numerical stability of covariance matrices was confirmed with the *base R* diagnostic functions *eigen*, *qr*, *determinant*, and *kappa*. Analyses were skipped if conditions for positive definiteness or finite log-determinants were not met. Runtime was recorded from PCA through the calculation of all MVNH metrics.

5. **PCA-space.** We estimated Schoener's D (Schoener, 1968) along PC1–PC2 and the niche divergence plane (Ascanio et al., 2024) along PC1. Environmental predictors (scaled and centered) were transformed into orthogonal PCs via PCA using the *prcomp* function in *base R*. For the niche divergence plane (Ascanio et al., 2024), kernel densities were estimated for each group along PC1 with the *density* function in *base R* using Silverman's bandwidth rule (Silverman, 1986). Densities were normalized to a maximum of one, and supports were defined as the range of grid values with density greater than $10^{-6}$. We then calculated the four niche divergence metrics (Ascanio et al., 2024). Niche dissimilarity (NDS) was calculated as the average of the two one-sided dissimilarities, each defined as one minus the integral of the minimum density over a group's support divided by the integral of its own density. Niche breadth exclusivity (NE) was calculated as one minus the ratio of the length of the overlap of the two supports to the length of their union. Niche divergence magnitude (ND) was calculated as the square root of the sum of squared NDS and squared NE. Niche divergence angle (θ) was defined as the arctangent of NDS divided by NE. NDS and NE both range from zero (complete similarity or overlap) to one (no density overlap or disjoint supports), ND ranges from zero (no divergence) to around 1.41 (square root of two; maximum divergence when both NDS and NE equal 1), and θ ranges from zero degree (purely exclusivity-driven divergence) to 90° (purely dissimilarity-driven divergence), with intermediate values indicating mixed contributions. Schoener's D as calculated in PC1–PC2 space by estimating two-dimensional Gaussian kernel densities for each group via the *kde2d* function in *MASS* with normal-reference bandwidths, evaluated on a shared 100 × 100 grid. Densities were normalized to sum to one, and Schoener's D was computed as the sum of the minimum of the two densities across all grid cells (equivalent to one minus one-half times the sum of absolute differences between the densities). Schoener's D ranges from zero (no overlap) to one (identical niches). Runtime was recorded from PCA through kernel density estimation and metric calculations. The results were visualized via *ggplot* as PC1–PC2 point clouds with overlaid density contours via the *stat_density_2d* function, and by plotting normalized PC1 kernel-density curves for both species estimated with the *density* function in *base R* stats.

6. **Logistic regression.** We tested for niche divergence using logistic regression of species identity on PCs of environmental predictors (Hosmer & Lemeshow, 2000; Huang et al., 2017; McCullagh & Nelder, 1989). The number of retained PCs was determined by the events-per-variable guideline (Peduzzi et al., 1996; Vittinghoff & McCulloch, 2007). Under this rule, at least ten outcome events in the smaller class are required per predictor to ensure stable

coefficient estimation and valid inference. In our case, this corresponds to limiting the number of retained PCs to no more than one-tenth of the per-group sample size. This conservative criterion reduces the risk of overfitting, biased regression coefficients, and undercoverage of confidence intervals (Austin & Steyerberg, 2017; Peduzzi et al., 1996). Occurrences of the two species were coded as a binary response variable (*0 = species 1*, *1 = species 2*) and modeled with a generalized linear model using the *glm* function in *base R* with a binomial error distribution and logit link (*family = stats::binomial()*). In this formulation, the logit of the probability that an observation belongs to species 2 is equal to an intercept plus the sum of regression coefficients multiplied by the retained PC scores. Model significance was evaluated by comparing the full model to an intercept-only null model using a likelihood-ratio test (McCullagh & Nelder, 1989) implemented with the *anova* function in *base R stats*. This test compares the log-likelihoods of the two models, with the difference asymptotically following a $\chi^2$ distribution. It uses the null hypothesis that environmental predictors do not improve model fit (Huang et al., 2017). A significant likelihood-ratio test was interpreted as evidence that environmental predictors significantly distinguish species identity. Fitted probabilities from the model range from zero (corresponding to a certain assignment to species 1) to one (corresponding to a certain assignment to species 2). Runtime was recorded from predictor preprocessing through model fitting and likelihood-ratio testing. The results were visualized by plotting the fitted probability distributions by species with a rug overlay in *ggplot*. Additionally, partial-dependence style curves of fitted probability versus each retained PC were plotted with *ggplot*.

We initially also considered the HUMBOLDT niche divergence test (Brown & Carnaval, 2019) implemented in the *humboldt v.2.0.0.90825 R* package (Brown, 2025). This method provides an established framework for quantifying ecological niche divergence while explicitly correcting for background environmental availability, non-analogous environmental space, and potential niche truncation. However, pilot trials showed that this method did not complete within twelve hours for a single simulated dataset under default settings on our Windows system. This rendered the approach computationally impractical for two sets of 100 simulated species, in contrast to the other methods, which finished within minutes (Supplementary Table S8).

#### III) Empirical analyses (data, study extent, variables)

##### M7: *Hemileuca* taxonomic background and additional empirical examples

The main empirical niche-divergence analysis presented in the main text focuses on western versus Great Plains populations traditionally assigned to a single species, *Hemileuca nevadensis* Stretch, 1872. This taxonomic concept has long been applied to all *maia*-group populations ranging from western North America into the Great Plains (Ferguson, 1971; Lemaire, 2002). The northern Great Plains populations were historically described as a separate taxon, *Hemileuca lucina latifascia* Barnes and McDunnough, 1916 and later synonymized with *H. nevadensis* by Ferguson (1971). Based on disjunct distributions and mitochondrial DNA patterns, Schmidt (2022; preprint) proposed that the western and Great Plains populations should be treated as separate species-level taxa, namely *H. nevadensis* and "*H. latifascia*". However, the eastern limits of the "*H. latifascia*" remain unclear because mitochondrial haplotypes from Great Plains populations are shared with the "Great Lakes complex" populations to the east (Schmidt, 2022; Tuskes et al., 1996) and apparent distribution gaps, morphological clines, and generalized host-use acceptance of willows and cottonwoods obscure clear taxonomic assignment. Thus, most authors have treated these populations as part of a species complex rather than clearly delimited taxa (Dupuis et al., 2018; Schmidt, 2022; Tuskes et al., 1996). Likewise, a recent biogeographic study did not recognize "*H. latifascia*" and grouped the Great Plains and Great Lakes populations as a single operational taxonomic unit deserving of more study (Tuttle et al. 2026, in review). We therefore use "*H. latifascia*" and *H. nevadensis* as operational labels for Great Plains–Great Lakes and western populations, respectively, explicitly following Schmidt's (2022) illustrated distributions. This name usage is not meant to propose new taxonomic changes, and we maintain current species concepts (St Laurent, 2023). Instead, this pairing provides a geographically broad and taxonomically relevant case study for evaluating our niche divergence framework. Our analyses are intended to highlight patterns of niche differentiation that may motivate future taxonomic and field-based work on unresolved transitions, particularly between the Great Plains and Great Lakes.

In addition to the focal comparison presented in the main text (*Hemileuca nevadensis* and "*H. latifascia*"), we conducted two further empirical analyses within the *H. maia* species group to evaluate whether our DAPC niche divergence framework consistently identifies ecological differentiation across independently evolved lineages.

The first comparison examined *H. slosseri* Peigler & Stone, 1989 and *H. peigleri* Lemaire, 1981. These two taxa were historically included within a broad interpretation of *H. maia* but later recognized as distinct species. *Hemileuca peigleri* was originally described as the subspecies *H. maia peigleri* based on material from central Texas, where populations occupy oak-dominated uplands (Lemaire, 1981). Peigler & Stone (1989) then elevated this taxon to species rank. Peigler & Stone (1989) also described *H. slosseri* as a new species from populations in west Texas and southeastern New Mexico occurring in shinnery-oak (*Quercus havardii*) sand-plain habitats. Some syntheses treated these taxa within *H. maia sensu lato* (Lemaire, 2002; Tuskes et al., 1996), but recent ecological and genomic evidence support their recognition as distinct taxa within the *maia* group (Dupuis et al., 2018, 2020). Given their well-characterized differences in geographic distribution and habitat type, this pair provides an independent test of whether our DAPC framework detects corresponding multivariate environmental differentiation.

The second supplementary comparison evaluated ecological divergence between regional oak-associated populations of the *H. maia* species group, namely "*H. m. sandra*" Pavulaan, 2020 and "*H. m. orleans*" Pavulaan, 2020. These two geographically structured regional populations were originally described as subspecies of *H. maia* based on adult phenotypes, distributional discontinuities, and habitat associations (Pavulaan, 2020). "*Hemileuca m. sandra*" represents inland and northern populations associated with sandy oak–pine barrens and prairie–savanna mosaics of the upper Midwest and interior Northeastern United States, whereas "*H. m. orleans*" corresponds to Gulf Coast and Atlantic Coastal Plain populations inhabiting coastal pine–oak savannas, flatwoods, and sandy maritime-influenced ridges (Pavulaan, 2020). Earlier treatments of *H. maia* describe potential ecological distinctiveness of these regional habitats and document variation in host associations and phenology across the species' range (Lemaire, 2002; Tuskes et al., 1996). However, Tuttle et al. (2026, in review) recently found no support for recognizing both subspecies based on expanded occurrence data, overlapping regional distributions, broad morphological variation, shared host associations, and a lack of evidence for discrete geographic, ecological, or reproductive boundaries among oak-associated populations. Therefore, we use both names here only as operational labels to test the performance of our DAPC framework rather than as a test of taxonomic hypotheses. Considering the habitat and distributional differences proposed by Pavulaan (2020), we use "*H. m. sandra*" and "*H. m. orleans*" as an additional test of whether DAPC detects niche separation among these two lineages recently recognized as subspecies.

**M8: Compiling of occurrence data**

We compiled a comprehensive dataset of occurrence records for all *H. maia* group taxa by integrating three online databases: GBIF (Global Biodiversity Information Facility; gbif.org), iNaturalist (inaturalist.org), and Butterflies and Moths of North America (butterfliesandmoths.org). Records were supplemented by expert-curated datasets by the authors (JRD and CS) and Jim P. Tuttle (personal communication) containing specimen-based occurrences and expert-verified locality information. Occurrence records were carefully evaluated and manually reassigned at the subspecies level where necessary based on morphology, host associations, and distributions following Lemaire (2002), Pavulaan (2020) and Schmidt (2022). Considering that these pairings include taxonomically contentious taxa, we acknowledge that alternative interpretations based on expanded occurrence, morphological, ecological, and biogeographic evidence have been presented elsewhere (Tuttle et al., 2026, in review).

Research-grade iNaturalist records were downloaded using the *get_inat_obs* function in the *rinat v.0.1.1 R* package (V. Barve & Hart, 2025). Butterflies and Moths of North America records were manually downloaded as a CSV file from butterfliesandmoths.org. GBIF records were extracted using the *occ_search* function in the *rgbif v.3.8.2 R* package (Chamberlain et al., 2025). We excluded fossil records and coordinates flagged by automated cleaning routines (country centroids, duplicate centroids, capitals, institutions, GBIF headquarters, and zero coordinates) using the *clean_coordinates* function in the *CoordinateCleaner v.3.01.1 R* package (Zizka et al., 2019). Legacy coordinates reported in degrees–minutes–seconds were converted to decimal degrees for consistency. All downloaded records were extracted on the 4th of November 2025. Records with location uncertainty greater than ten kilometers were excluded. To balance sample size with spatial accuracy, we created two datasets with accuracy of no greater than 700 meters and no greater than ten kilometers. County-level records were assigned to the ten-kilometer dataset. Here, we only present results based on the ten- kilometer dataset due to its larger sample sizes. We repeated the empirical analyses with the much scarcer 700-meter dataset, which mostly yielded qualitatively similar results (results not shown).

The final combined data are available at Zenodo (Link to be inserted) and contain 4425 and 6890 records for the 700-meter and ten-kilometer datasets, respectively. Finally, interactive maps showing the most updated distributions for each taxon based on these occurrences are available at Zenodo (Link to be inserted). These maps were created with the *leaflet* function in the *leaflet v.2.2.2 R* package (Cheng et al., 2024) and exported with the *saveWidget* function in the *htmlwidgets v.1.6.4 R* package (Vaidyanathan et al., 2023).

**M9: Extraction of environmental variables and background sampling**

We assembled a suite of abiotic and biotic environmental axes (Supplementary Fig. S3) from fifteen independent sources. For this, we used our *extract.env.and.background* function (see Supplementary Methods M1). We generated one million background points, erased lakes from the accessible area and land mask, and used a five-kilometer buffer distance to delimit the accessible area for background sampling. This buffer distance represents a conservative estimate of dispersal distances in saturniid moths (Franzén & Nilsson, 2007; Stevens et al., 2010; D. L. Wagner, 2010). Otherwise, default function parameters were used.

In addition to the implemented *"env.datasets"*, we added one custom dataset via the *custom.env.rasters* argument as a broad-scale proxy of predation pressure. For this, we used a global terrestrial bird species richness layer from the International Union for Conservation of Nature Red List spatial data release 2025. This raster represents total avian species richness per ten times ten kilometers grid cell, compiled from BirdLife International (iucnredlist.org/resources/other-spatial-downloads) range maps and distributed in an equal-area Mollweide projection (EPSG:54009).

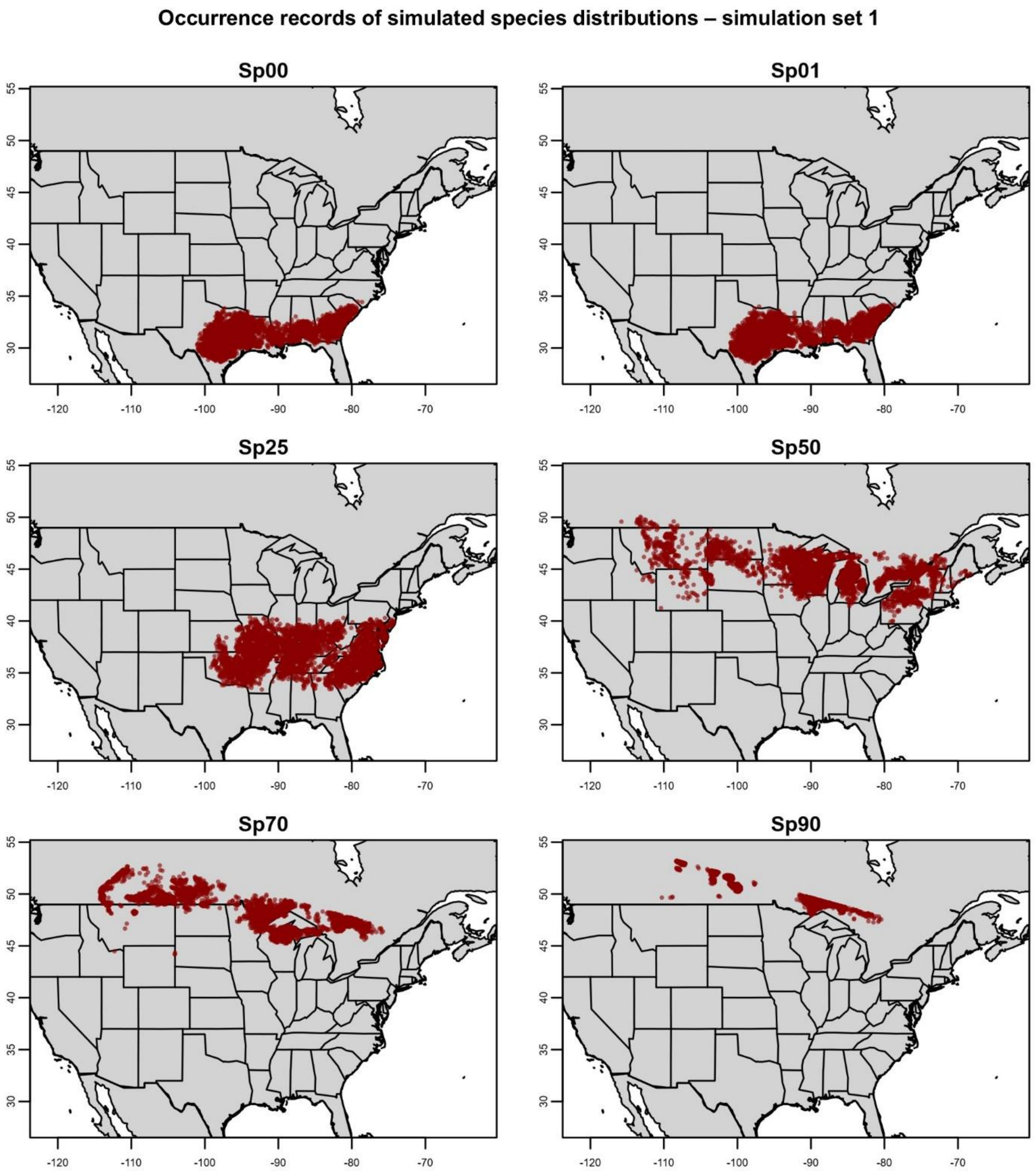


**Supplementary Figure S1: Distributions of simulated synthetic species across North America for simulation set 1 (multivariate divergence gradient generated along PC1–PC2 space).** Red dots represent simulated occurrence records used to extract environmental variables for our niche divergence test. Panels include the null species (Sp00) and five focal species. Abbreviation: PC = principal component.

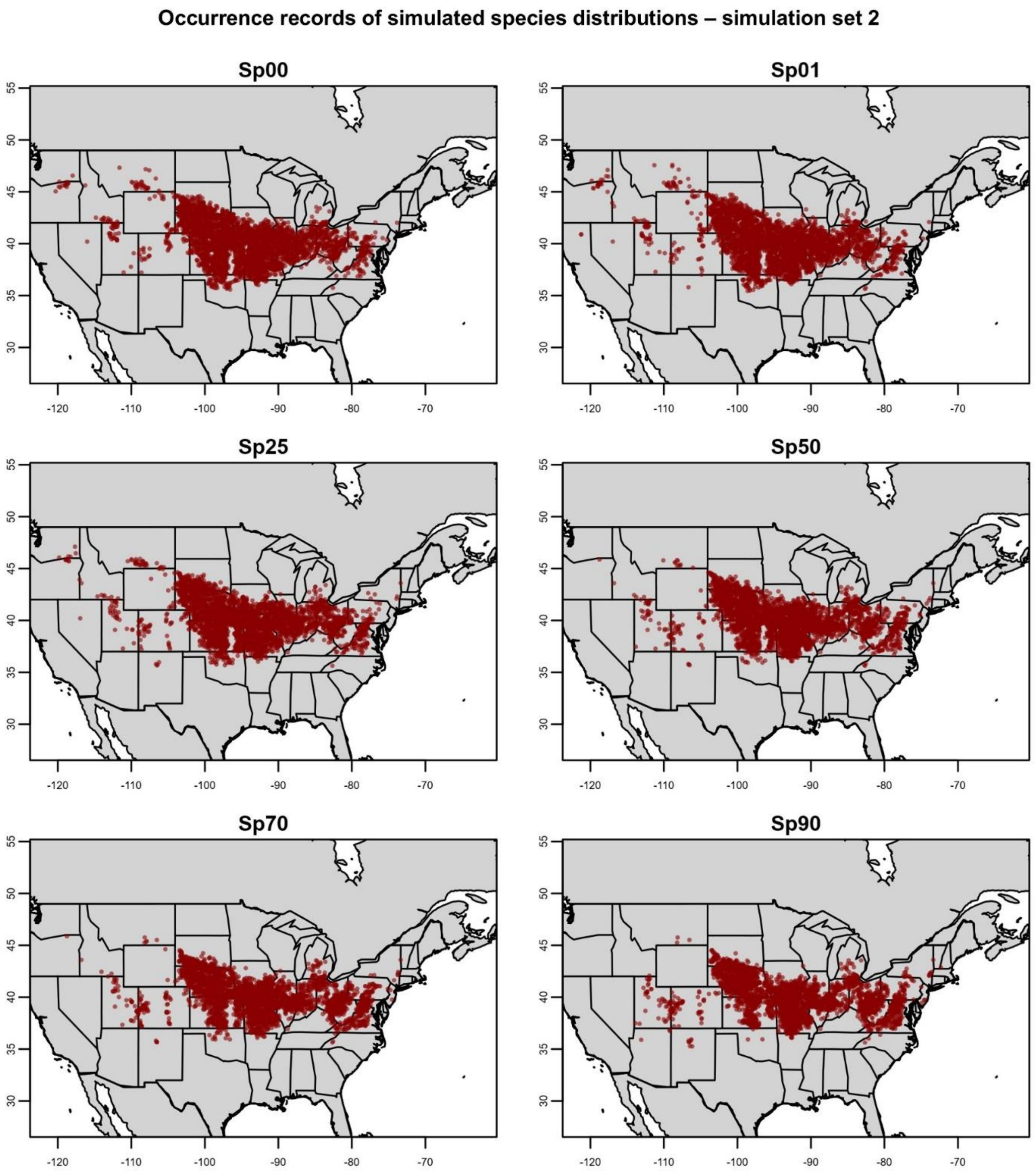


**Supplementary Figure S2: Distributions of simulated synthetic species across North America for simulation set 2 (multivariate divergence gradient generated along PC7–PC8 space).** Red dots represent simulated occurrence records used to extract environmental variables for our niche divergence test. Panels include the null species (Sp00) and five focal species. Abbreviation: PC = principal component.

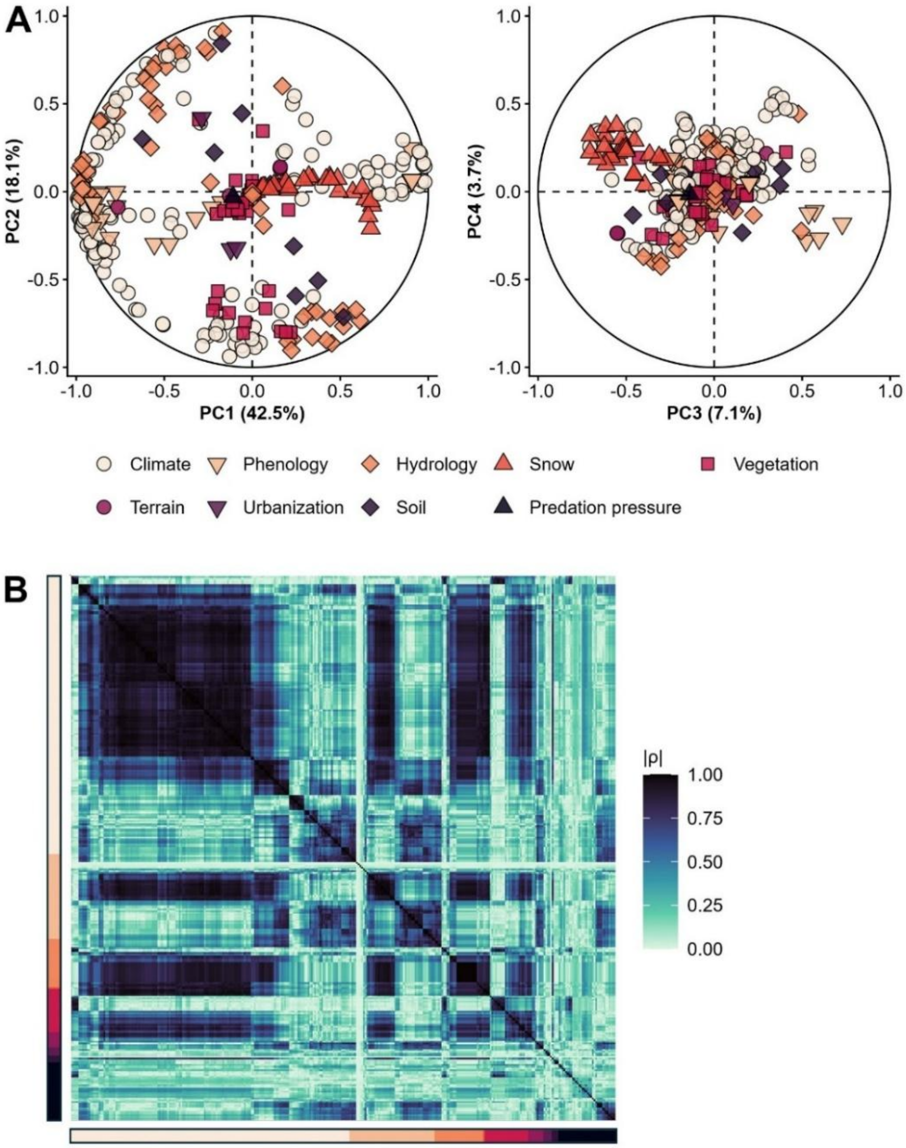


**Supplementary Figure S3: Correlations among all 328 environmental variables representing nine different categories. A)** PCA correlation circles for PC1–PC2 (left) and PC3–PC4 (right), with points representing variables colored by category. Points farther from the origin have stronger loadings and points near the center have weaker loadings on the two respective PCs. Variables in similar directions from the origin are positively associated, variables arranged roughly orthogonally show weak or no association, and variables in opposite directions are negatively associated. Clusters highlight categories that co-vary. Axis labels report variance explained for each PC. Variables were centered and scaled and missing values were imputed by column means prior to PCA. **B)** Heatmap of absolute pairwise Spearman correlations (computed on ranked values with pairwise completeness). Variables are first ordered by category and clustered within category using average-linkage hierarchical clustering on the distance one minus the absolute Spearman correlation. Light to dark colors indicate low to high absolute pairwise correlation. Side strips encode categories of variables as in panel A. Mean absolute pairwise correlation coefficient across all variables: 0.41.

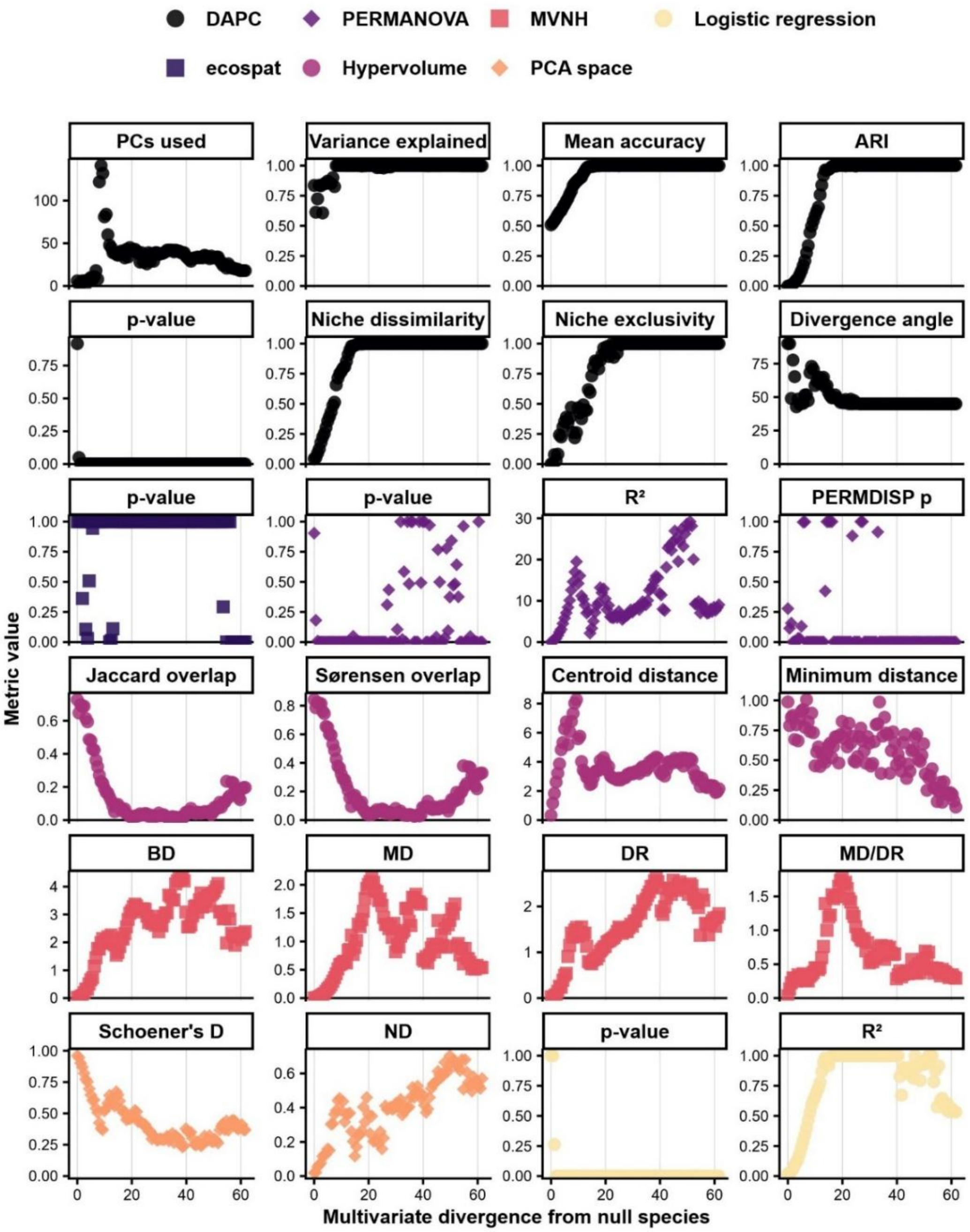


**Supplementary Figure S4: Niche divergence test results for simulation set 1 across 100 simulations across multivariate PC1–PC2 suitability space.** Abbreviations: ARI = Adjusted Rand Index, BD = Bhattacharyya distance, DAPC = discriminant analysis of principal components, DR = determinant ratio, MD = Mahalanobis distance, MD/DR = Mahalanobis distance divided by determinant ratio, MVNH = multivariate normal hypervolume, ND = niche divergence magnitude, PCA = principal component analysis, PCs = principal components, PERMANOVA = permutational multivariate analysis of variance, PERMDISP p = p-value from permutation test of homogeneity of multivariate dispersions, $R^2$ = proportion of explained model variance (for logistic regression: McFadden pseudo-$R^2$ goodness-of-fit measure), theta = niche divergence angle.

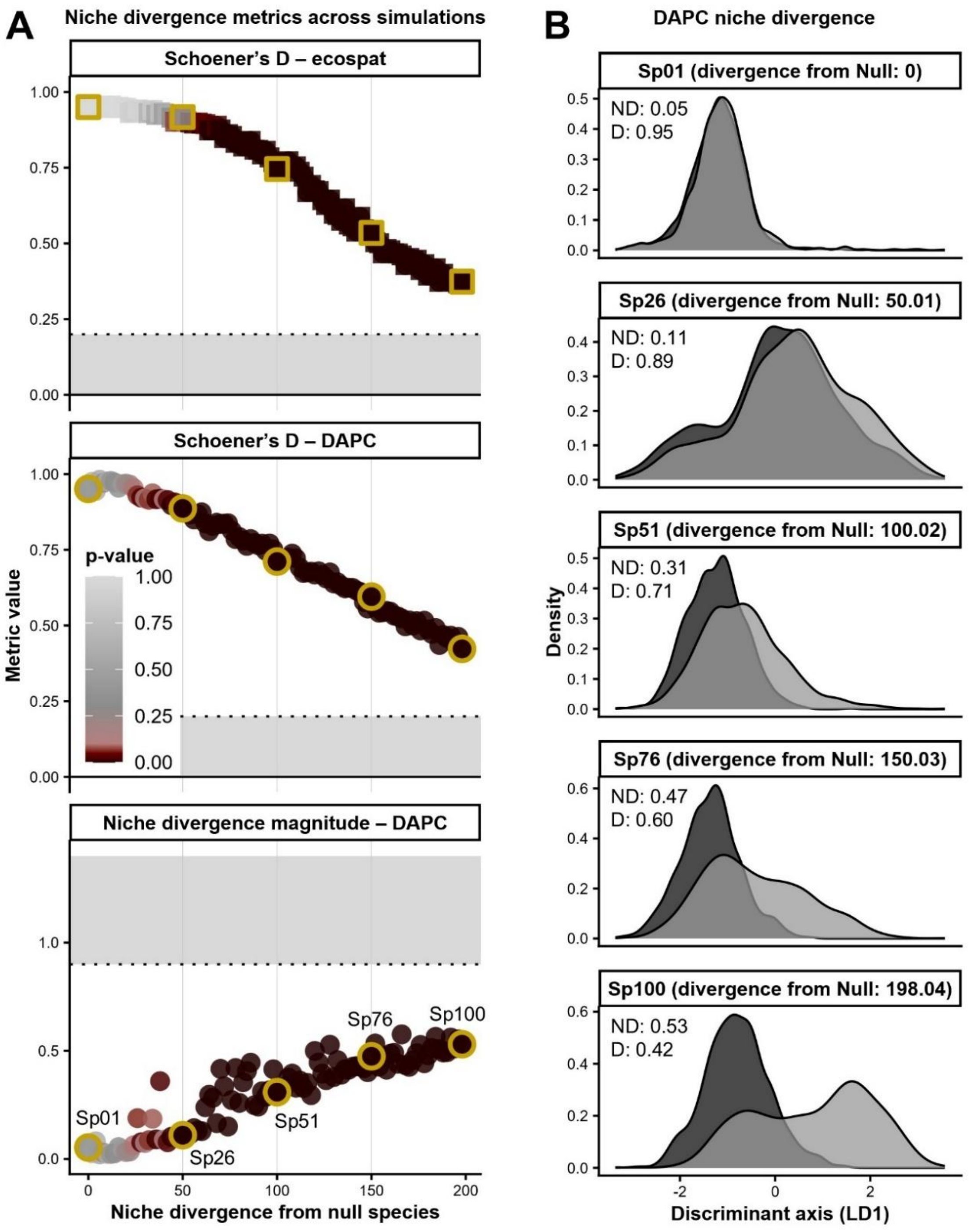


**Supplementary Figure S5: Multivariate niche divergence across 100 virtual species simulated with step-wise increasing levels of divergence in PC7–PC8 suitability space (simulation set 2).** **A)** Divergence from the Null species Sp00 vs. Schoener's D and ND (niche divergence magnitude) from *ecospat* and our DAPC method, with points as simulations, focal species outlined in golden, point color encoding p-value, and grey area representing strong niche divergence ($D \leq 0.2$; $ND \geq 0.9$). **B)** LD1 density curves of DAPC for the five focal species with increasing divergence from the Null species. Sp01 (top) represents simulated niche fully shared relative to null species ($k = 1$).

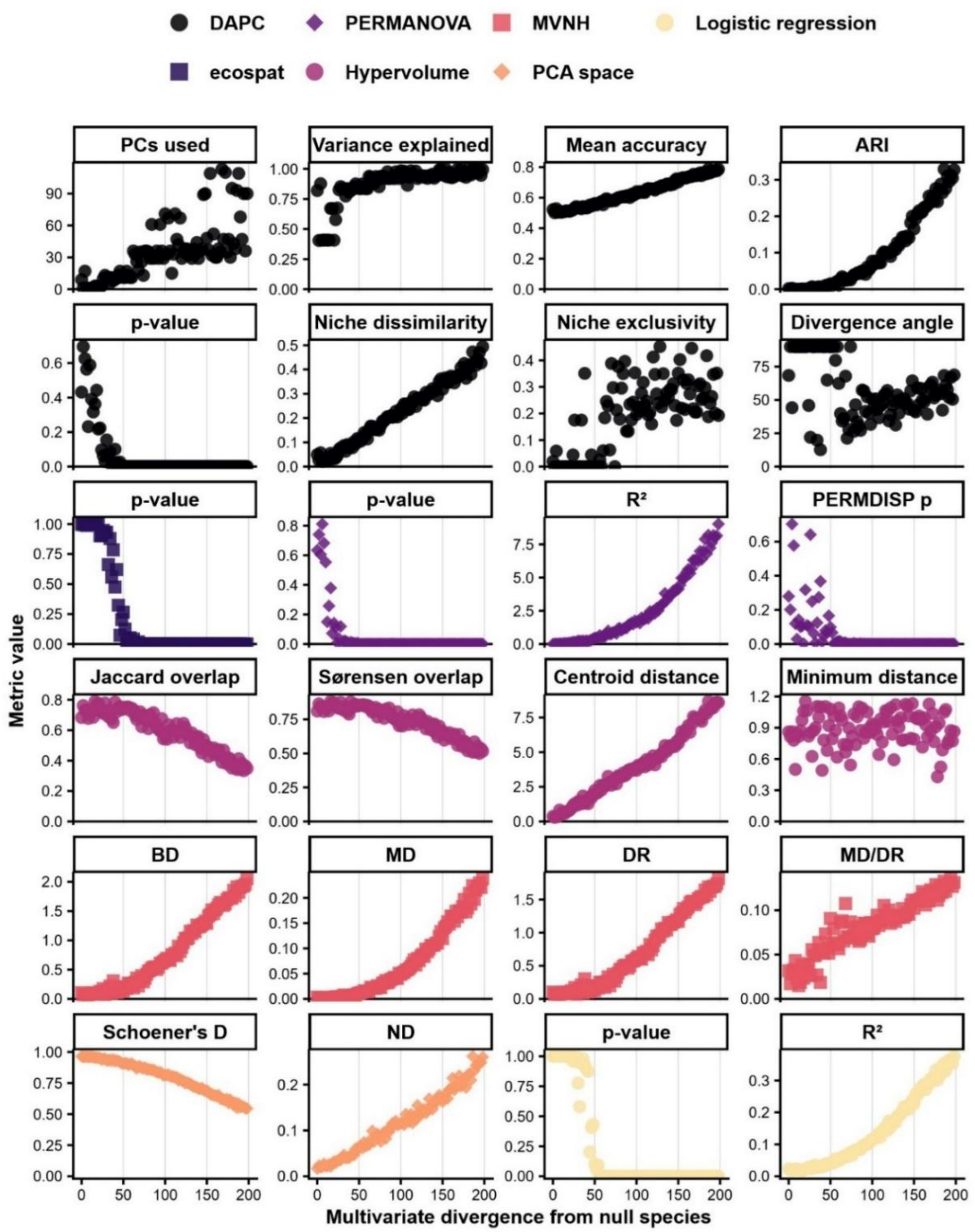


**Supplementary Figure S6: Niche divergence test results for simulation set 2 across 100 simulations across multivariate PC7–PC8 suitability space.** Abbreviations: ARI = Adjusted Rand Index, BD = Bhattacharyya distance, DAPC = discriminant analysis of principal components, DR = determinant ratio, MD = Mahalanobis distance, MD/DR = Mahalanobis distance divided by determinant ratio, MVNH = multivariate normal hypervolume, ND = niche divergence magnitude, PCA = principal component analysis, PCs = principal components, PERMANOVA = permutational multivariate analysis of variance, PERMDISP p = p-value from permutation test of homogeneity of multivariate dispersions, $R^2$ = proportion of explained model variance (for logistic regression: McFadden pseudo-$R^2$ goodness-of-fit measure), theta = niche divergence angle.

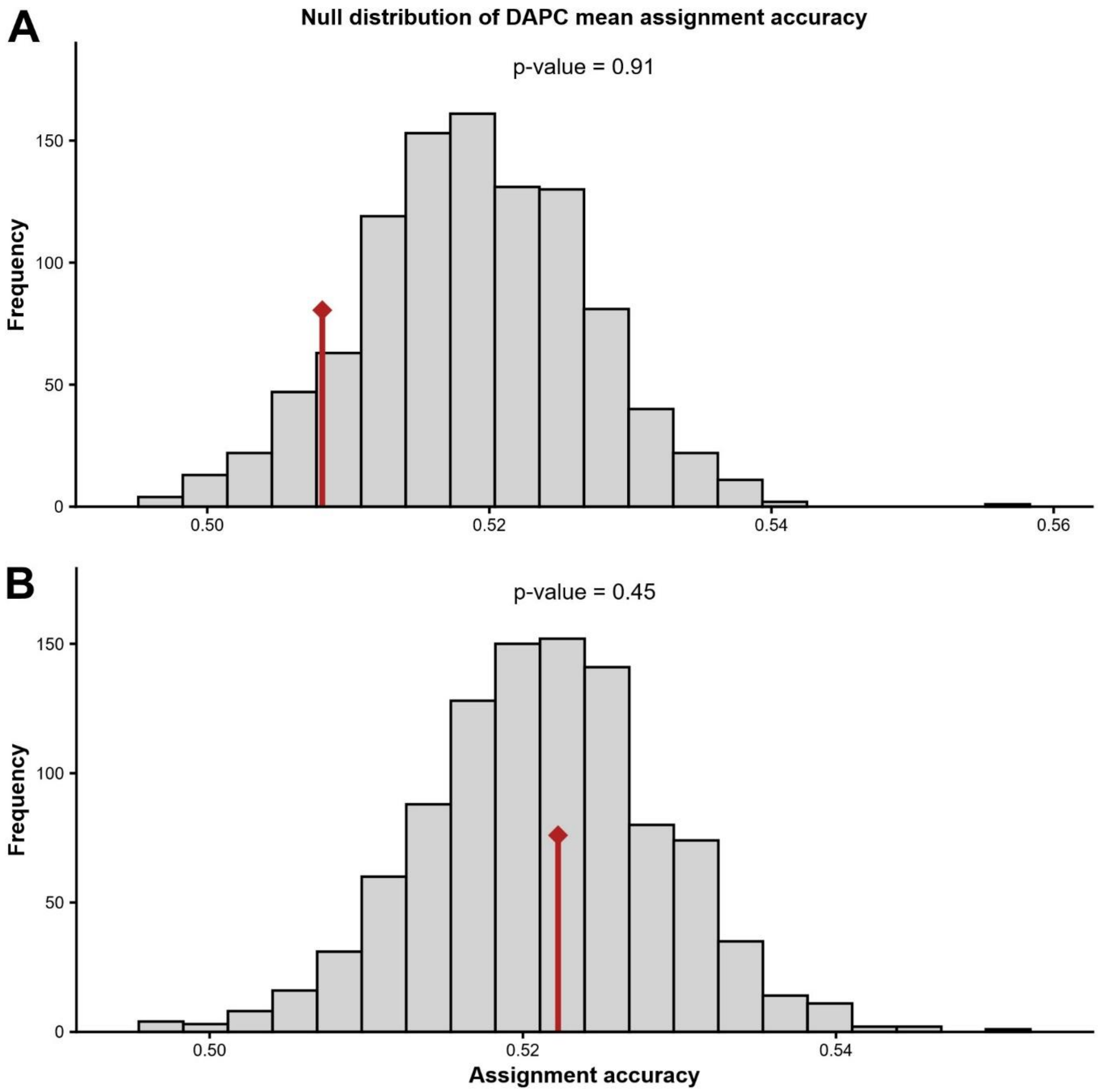


**Supplementary Figure S7: DAPC permutation results for Sp00 vs. Sp01 corresponding to no niche divergence for simulation set 1 (A) and set 2 (B).** Grey bars show the null distribution as permutation values while the red line the shows observed value.

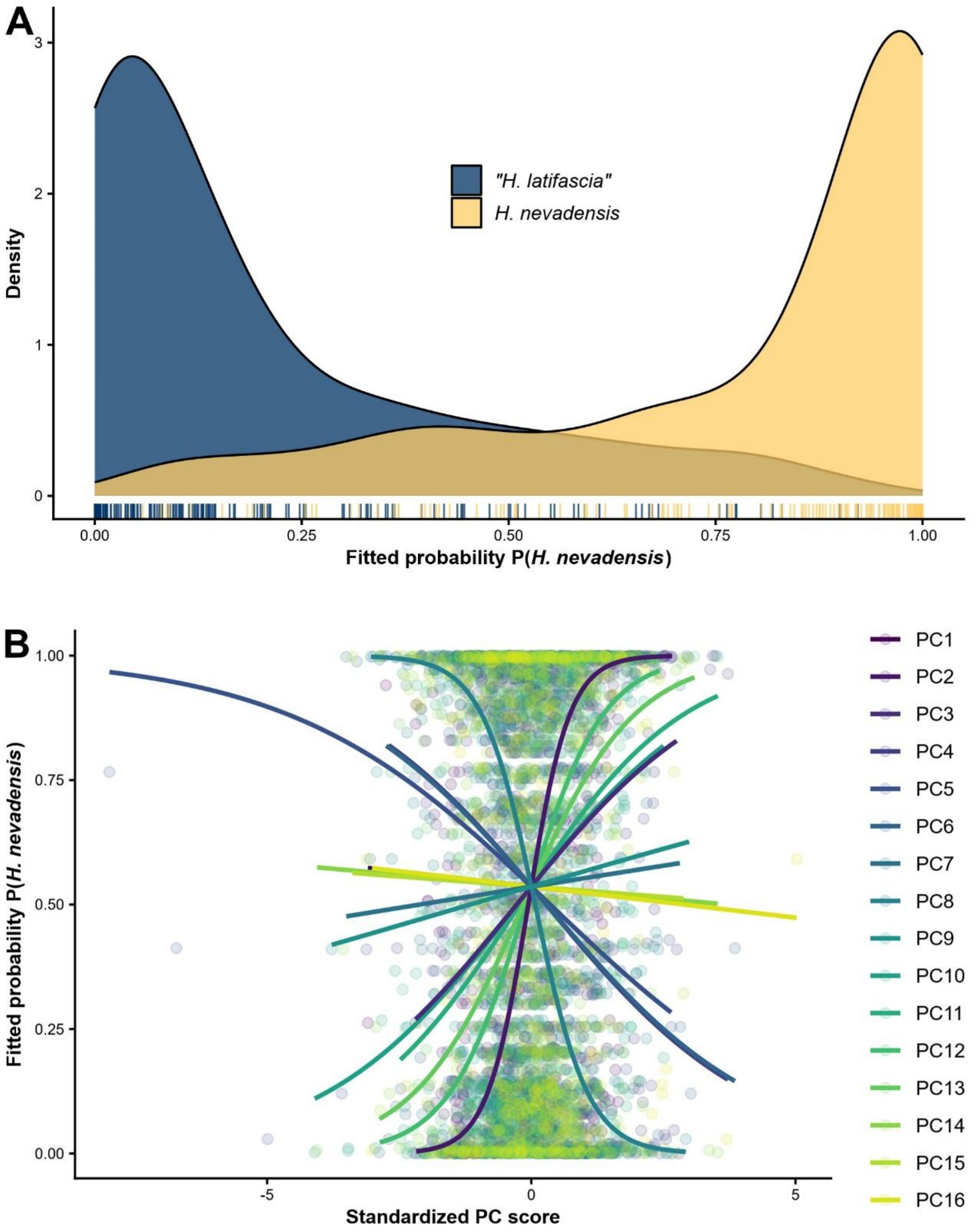


**Supplementary Figure S8: Logistic regression classification based on binomial generalized linear model (logit link) of "*Hemileuca latifascia*" vs. *H. nevadensis* (empirical dataset 1) using sixteen principal components (PCs) of analogous environmental predictors. A)** X-axis shows fitted model probability *P* that a record belongs to *H. nevadensis*, curves show kernel density estimates of fitted probabilities for each species, and rug ticks mark individual predictions. **B)** Partial dependence curves showing fitted probabilities based on model predictions along standardized PC scores, with points marking observed values. Steep curves indicate stronger niche divergence.

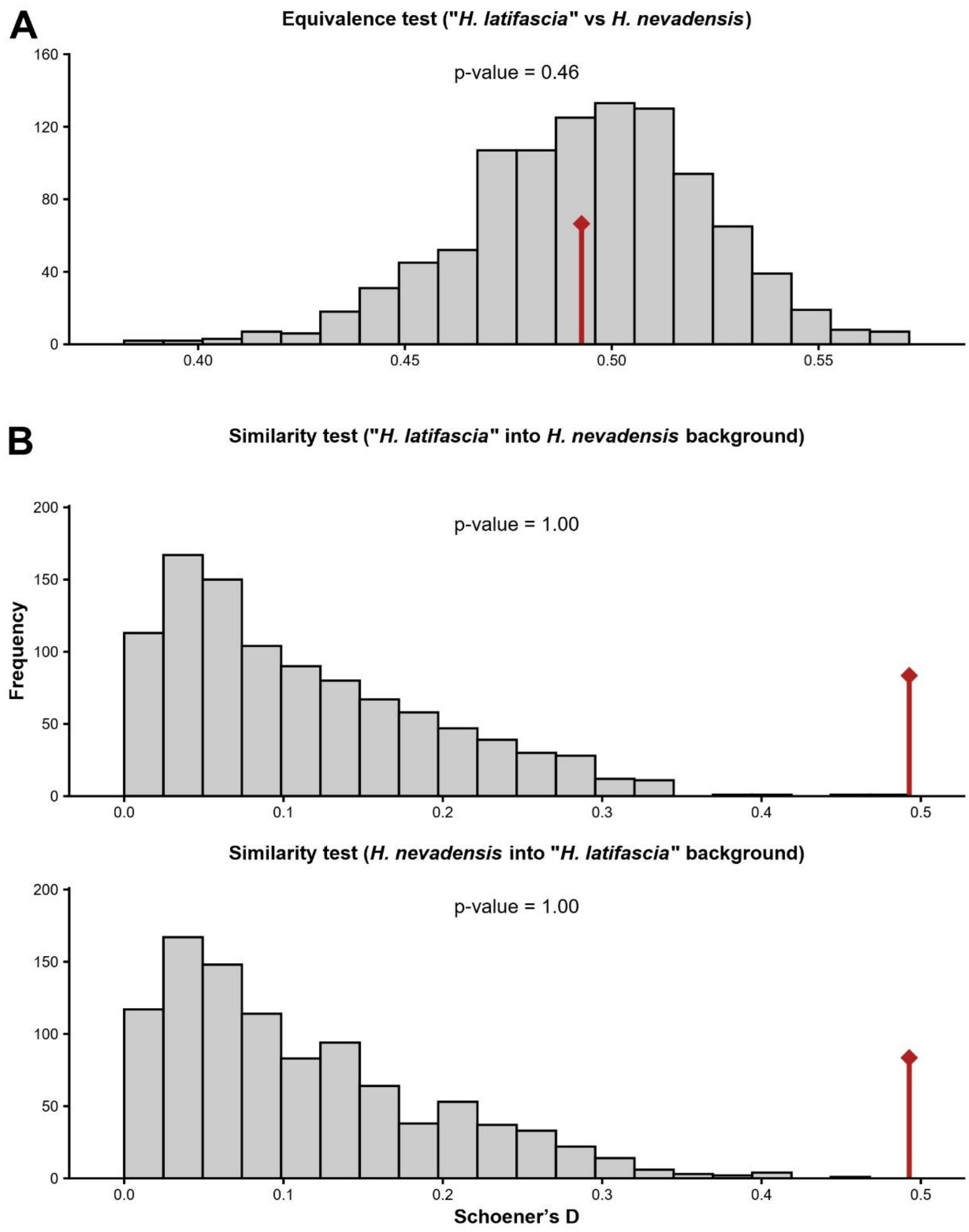


**Supplementary Figure S9: Permutation histograms for *ecospat* niche overlap tests between "*Hemileuca latifascia*" and *H. nevadensis* (empirical dataset 1).** Grey bars show null distributions generated by randomizing species occurrences within background space and red lines mark observed values. **A)** Niche equivalence test: evaluates whether the environmental niches of "*H. latifascia*" and *H. nevadensis* are statistically indistinguishable. **B)** Niche similarity test: examines whether the niche of "*H. latifascia*" is more similar to the available background of *H. nevadensis* than expected by chance (upper row), and vice versa (lower row). Observed values in the null tails indicate rejection of equivalence or similarity, consistent with niche divergence or asymmetry.

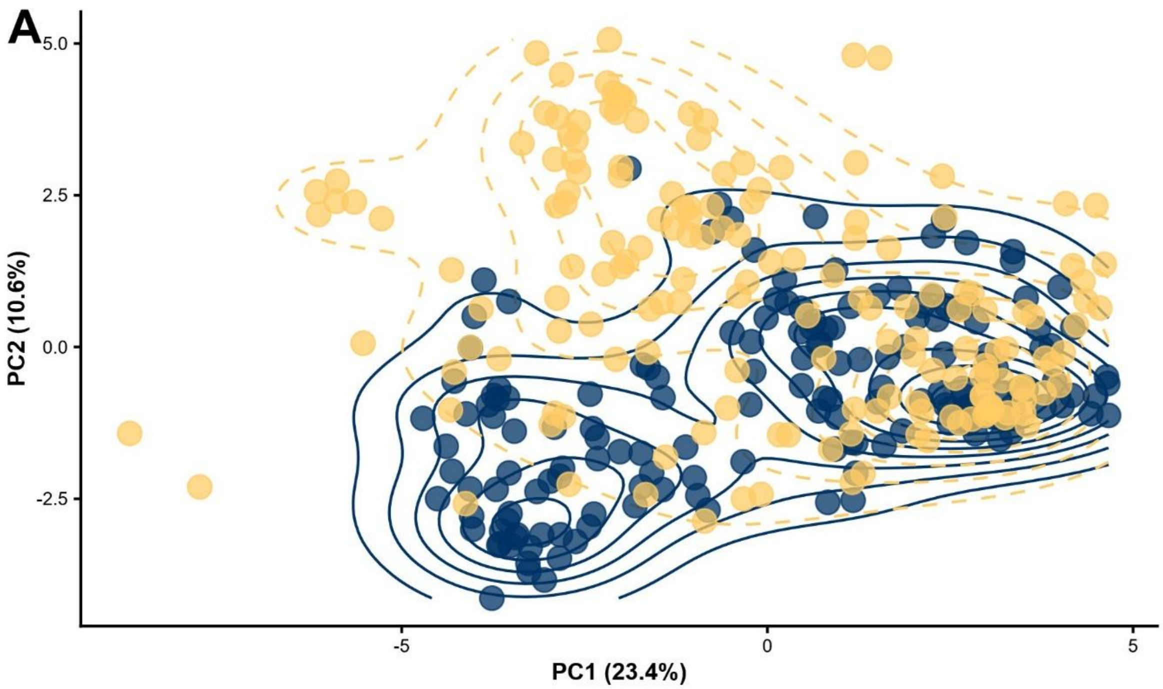


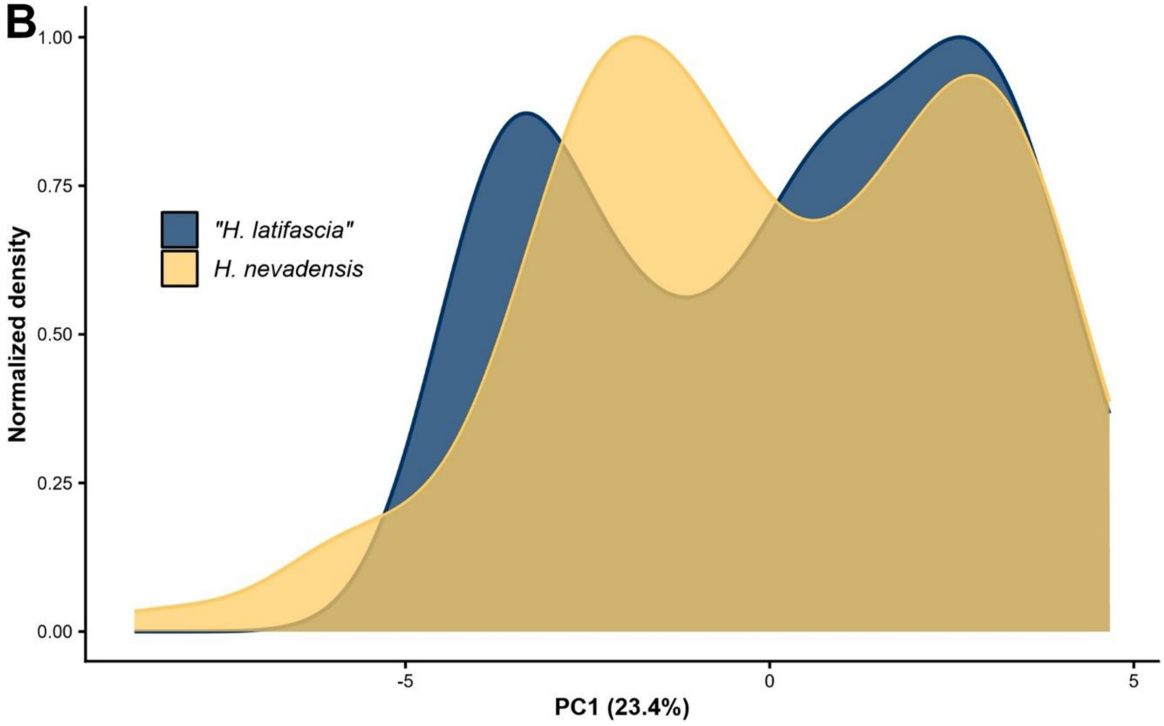


**Supplementary Figure S10: Niche divergence in analogous environmental principal component analysis space between "*H. latifascia*" and *H. nevadensis* (empirical dataset 1). A)** Ordination of principal components (PC) 1 and 2 with occurrences plotted as points and occurrence densities (kernel-density isopleths) as contours. **B)** Density curves along PC1 (normalized to unit). Axis labels show the variance explained by each PC in parenthesis.

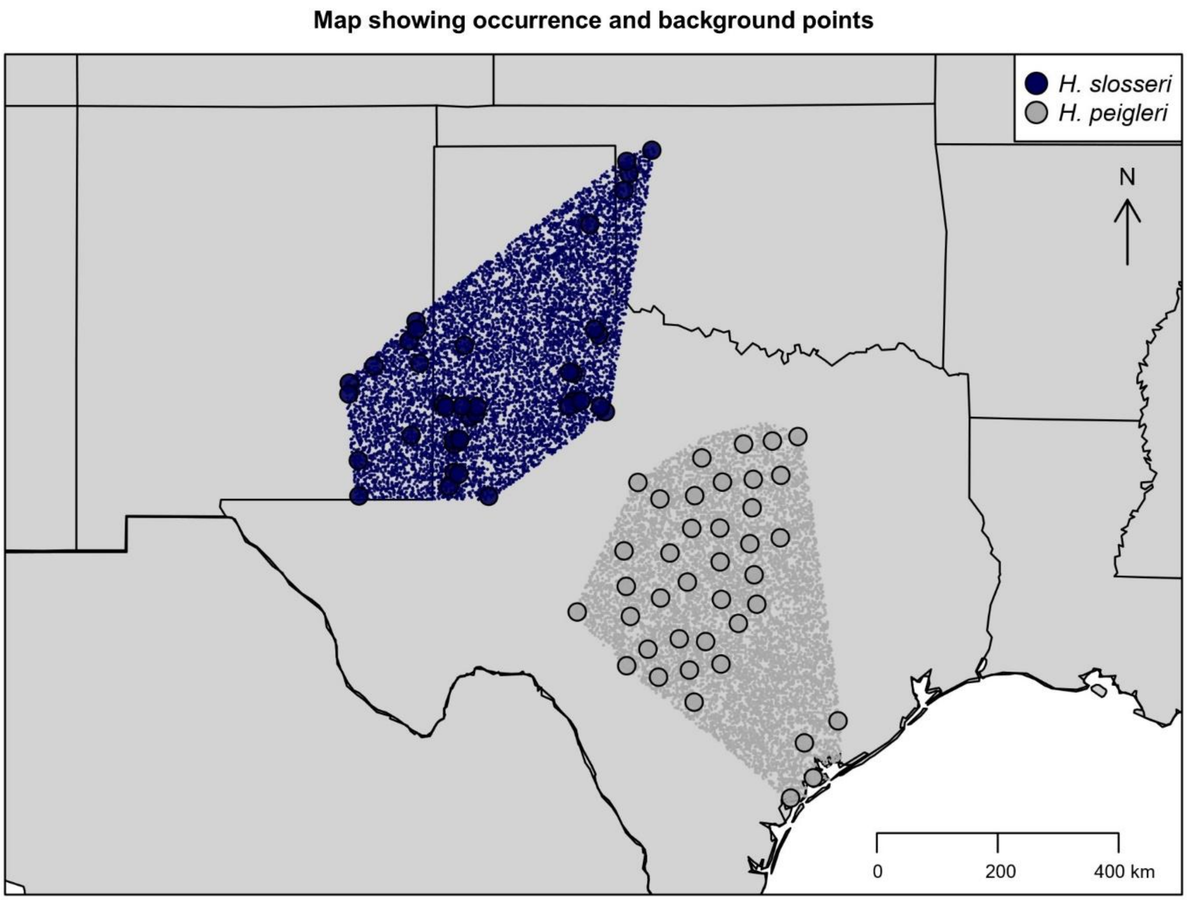


**Supplementary Figure S11: Occurrence records (large points) and background points (small points) for *Hemileuca slosseri* and *H. peigleri* (empirical dataset 2) across the United States.** Shown are thinned and down-sampled records used as input for our DAPC (discriminant analysis of principal components) niche divergence test.

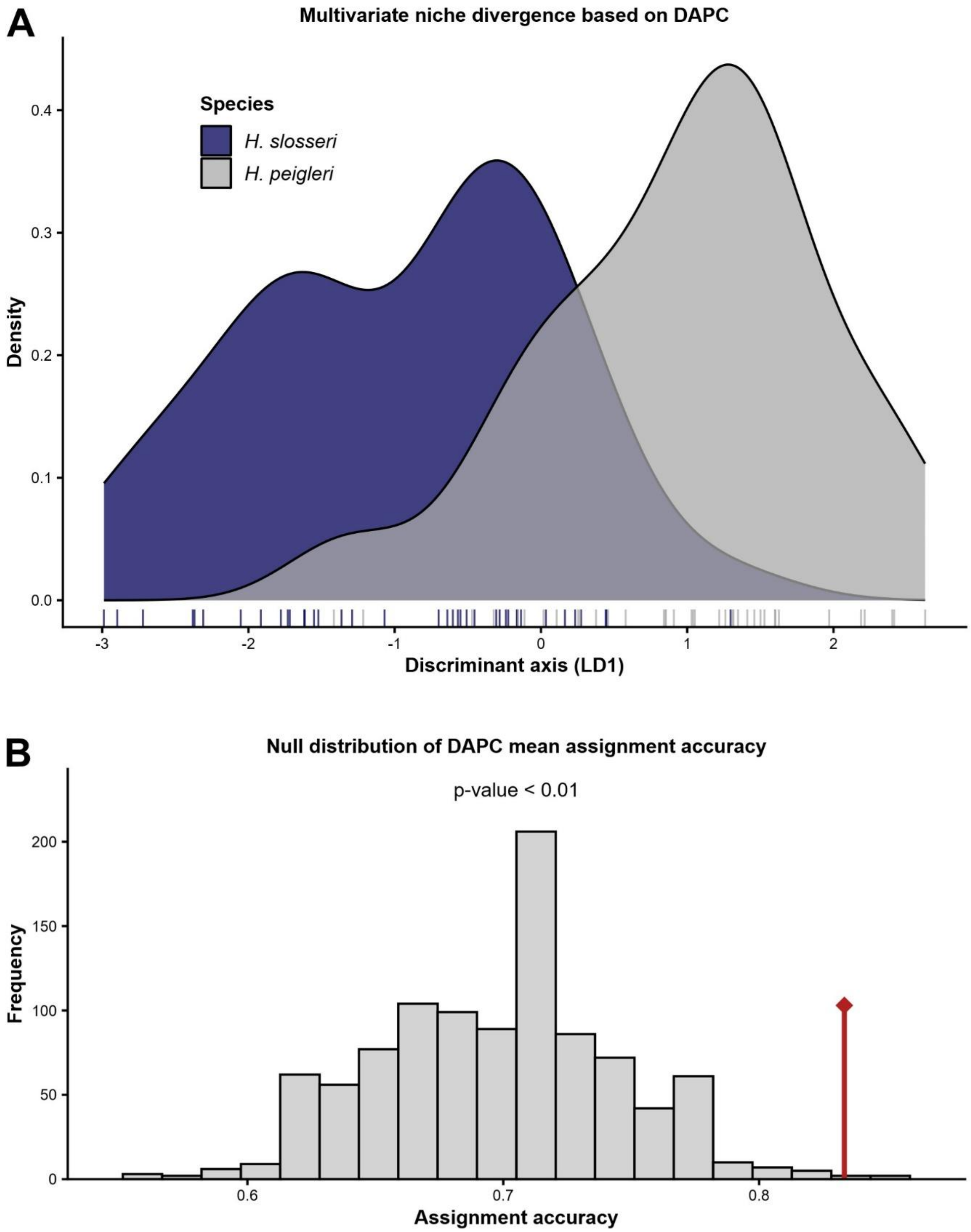


**Supplementary Figure S12: Discriminant analysis of principal components (DAPC) approach revealed moderate and significant multivariate niche divergence between *Hemileuca slosseri* and *H. peigleri* (empirical dataset 2). A)** Divergence along the discriminant axis (LD1). **B)** Null distributions of DAPC classification accuracy and misclassification rates from permutation tests (grey bars) with observed values shown as red lines.

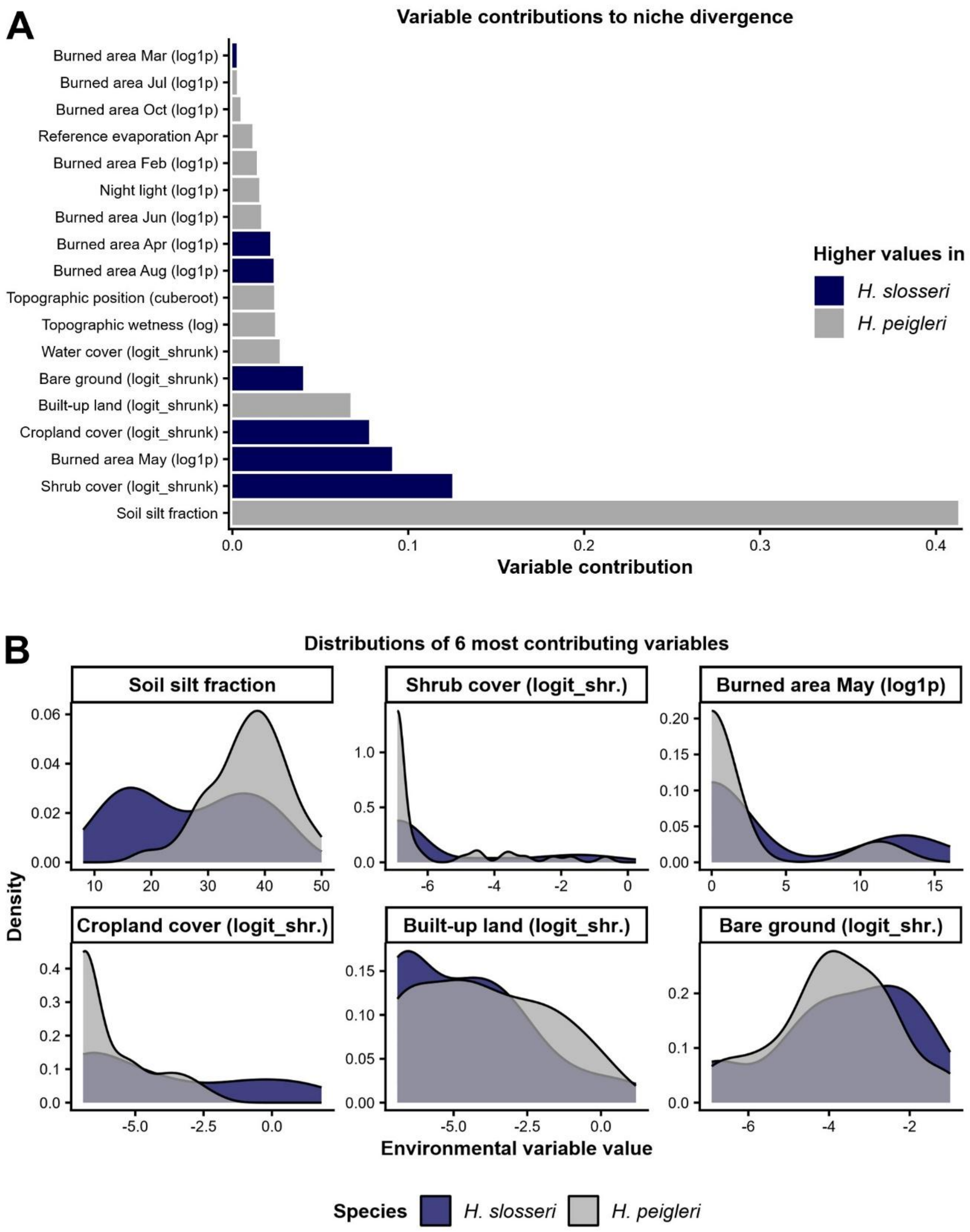


**Supplementary Figure S13: Contributions of environmental variables to multivariate niche divergence based on discriminant analysis of principal components (DAPC) niche divergence test between *Hemileuca slosseri* and *H. peigleri* (empirical dataset 2). A)** DAPC variable loadings to discriminant axis, with bar colors indicating the species with higher raw variable values. **B)** Kernel density plots of the raw distributions of the six top-contributing variables for both species. Months are abbreviated as three letter codes. Data transformations are abbreviated in parentheses.

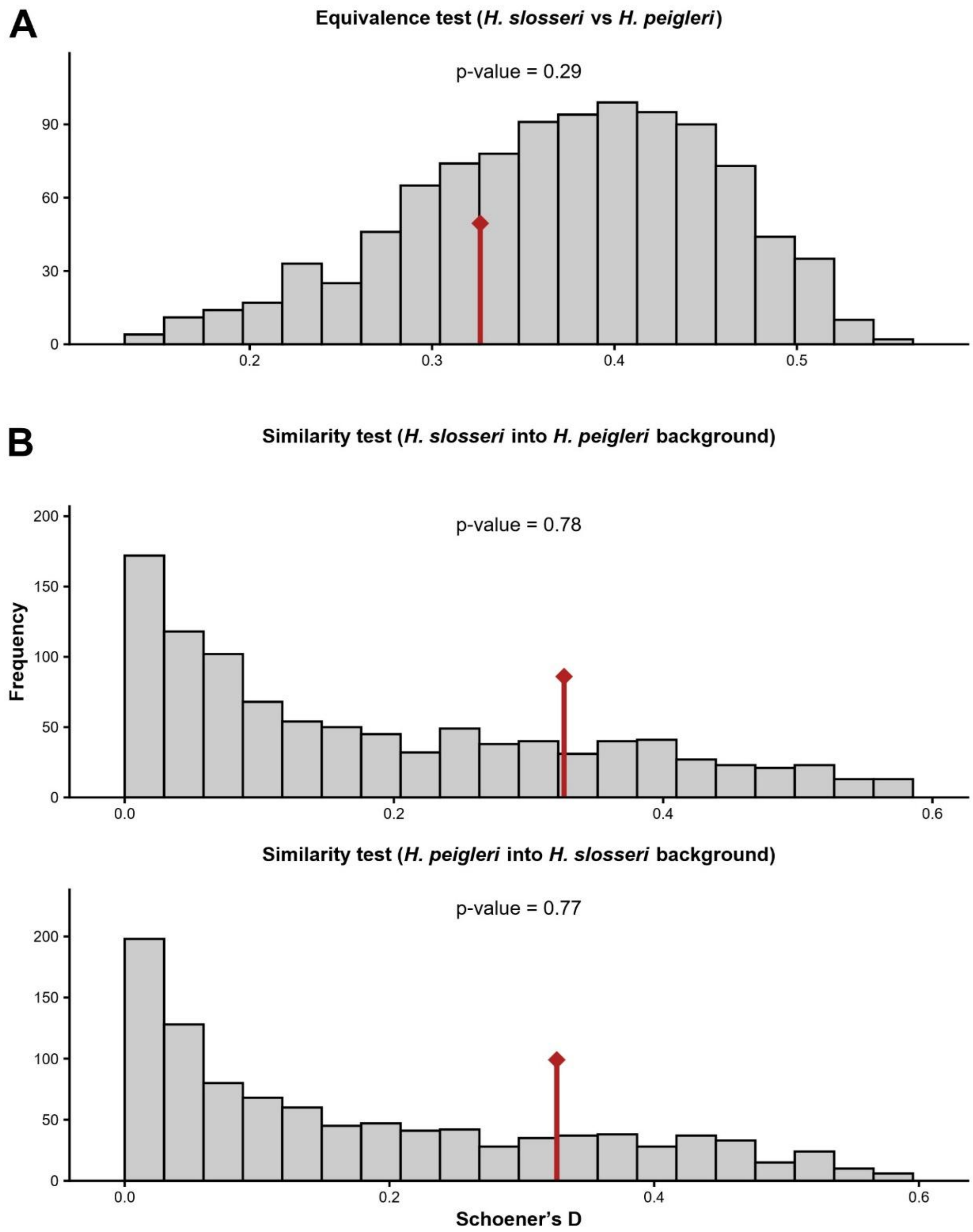


**Supplementary Figure S14: Permutation histograms for *ecospat* niche overlap tests between *Hemileuca slosseri* and *H. peigleri* (empirical dataset 2).** Grey bars show null distributions generated by randomizing species occurrences within background space and red lines mark observed values. **A)** Niche equivalence test: evaluates whether the environmental niches of *H. slosseri* and *H. peigleri* are statistically indistinguishable. **B)** Niche similarity test: examines whether the niche of *H. slosseri* is more similar to the available background of *H. peigleri* than expected by chance (upper row), and vice versa (lower row). Overlap was quantified using Schoener's D. Observed values in the null tails indicate rejection of equivalence or similarity, consistent with niche divergence or asymmetry.

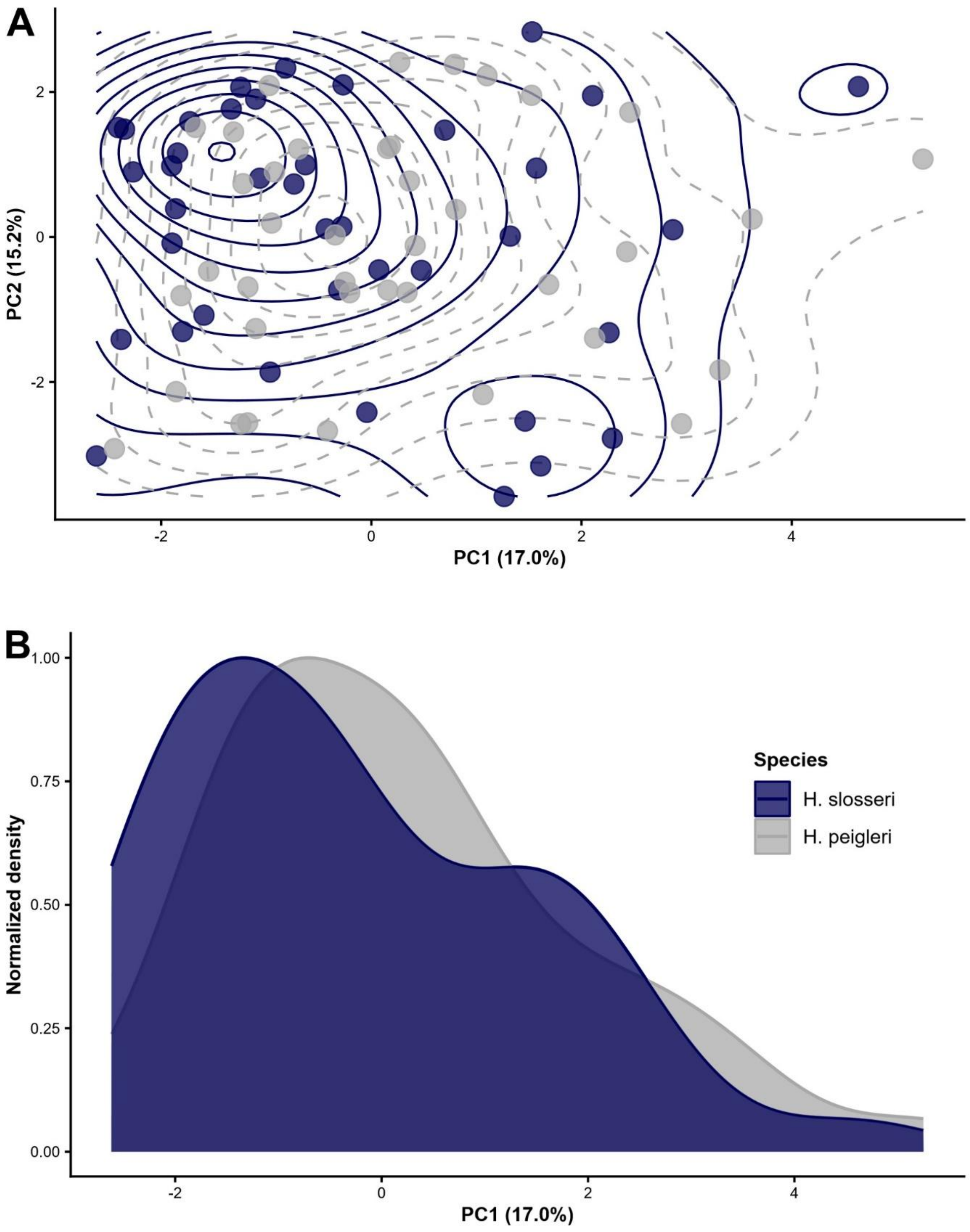


**Supplementary Figure S15: Niche divergence in analogous environmental principal component analysis pace between *H. slosseri* and *H. peigleri* (empirical dataset 2). A)** Ordination of principal components (PC) 1 and 2 with occurrences plotted as points and occurrence densities (kernel-density isopleths) as contours. **B)** Density curves along PC1 (normalized to unit). Axis labels show the variance explained by each PC.

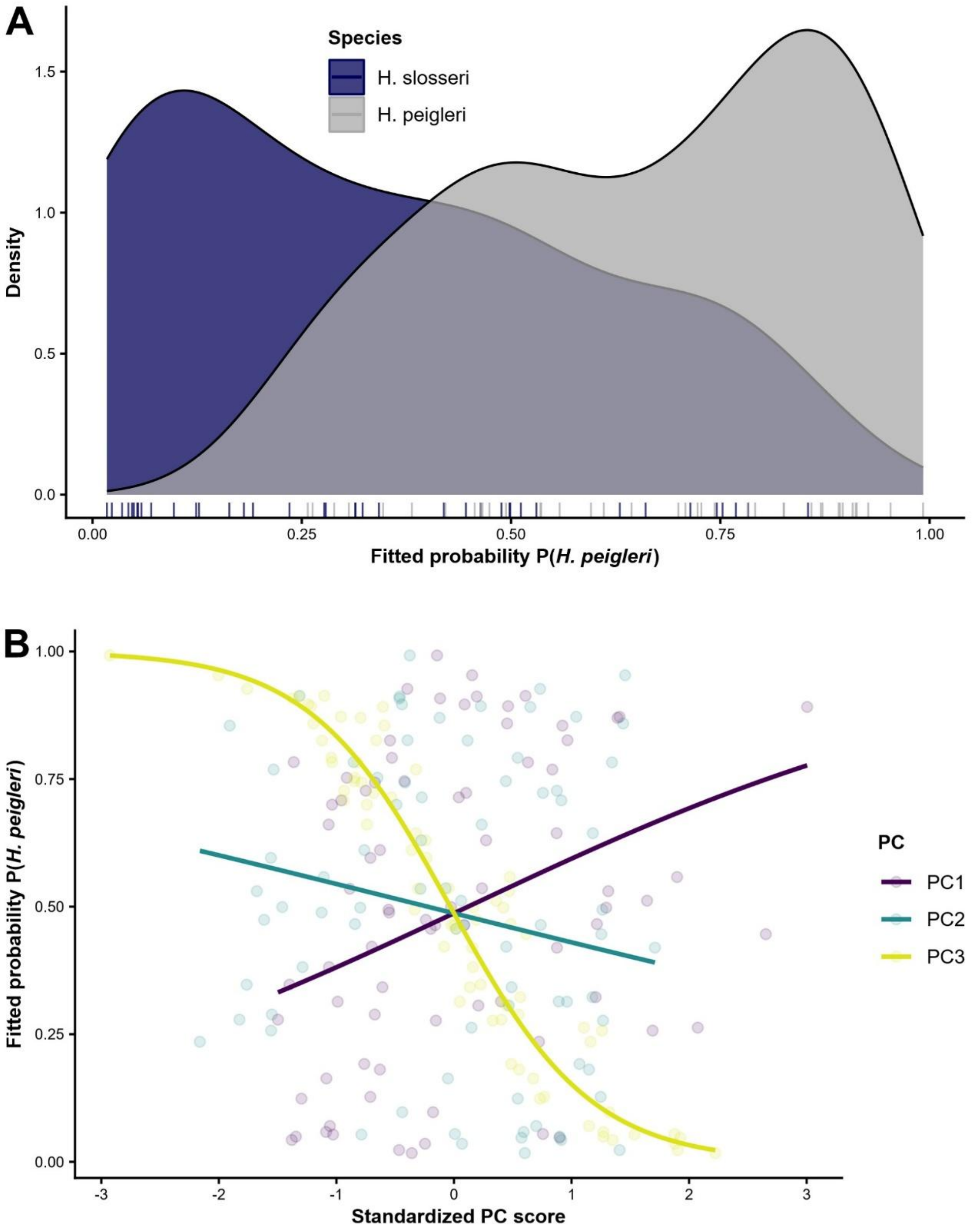


**Supplementary Figure S16: Logistic regression classification based on binomial generalized linear model (logit link) of *Hemileuca slosseri* vs. *H. peigleri* (empirical dataset 2) using three principal components (PCs) of analogous environmental predictors. A)** X-axis shows fitted model probability *P* that a record belongs to *H. slosseri*, curves show kernel density estimates of fitted probabilities for each species, and rug ticks mark individual predictions. **B)** Partial dependence curves showing fitted probabilities based on model predictions along standardized PC scores, with points marking observed values. Steep curves indicate stronger niche divergence.

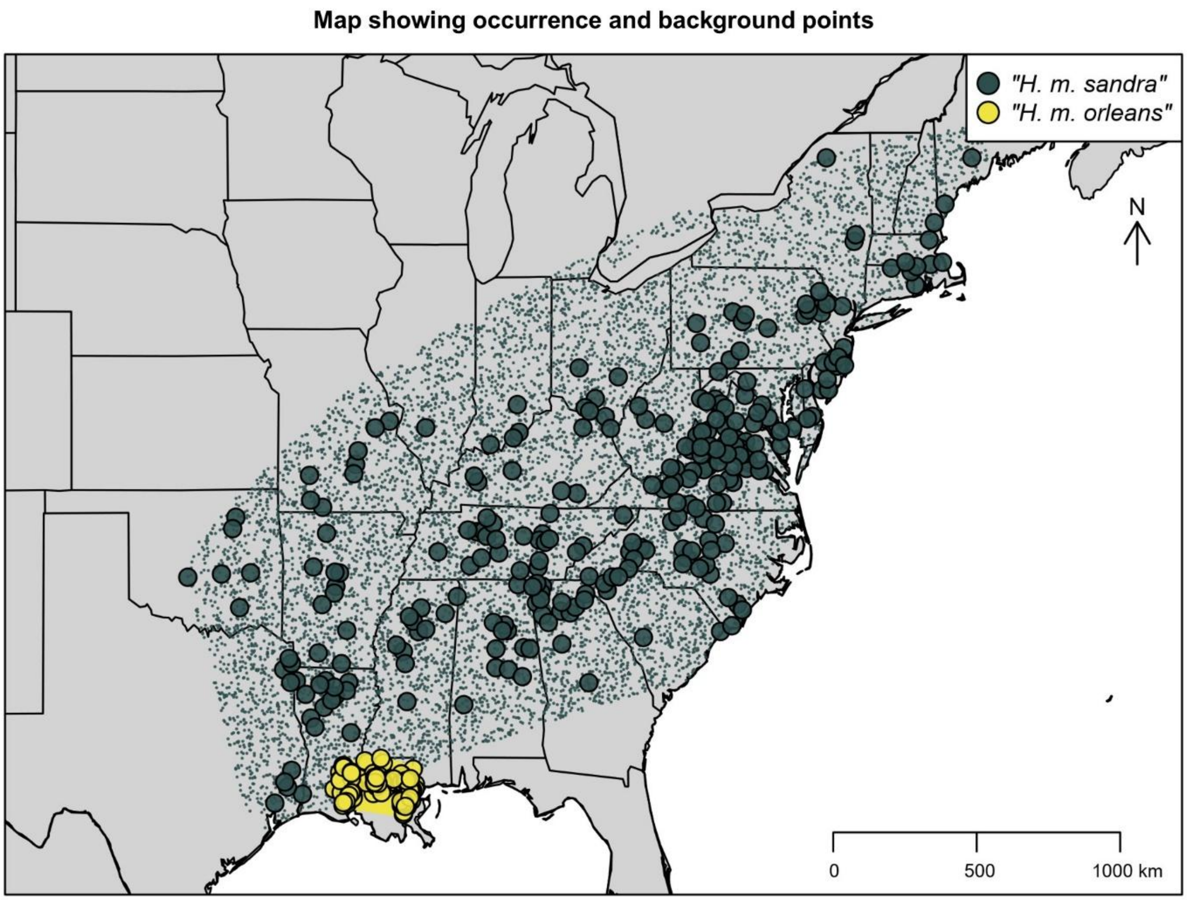


**Supplementary Figure S17: Occurrence records (large points) and background points (small points) for "*Hemileuca maia sandra*" and "*H. m. orleans*" (empirical dataset 3) across Eastern United States.** Shown are thinned and down-sampled records used as input for our DAPC (discriminant analysis of principal components) niche divergence test.

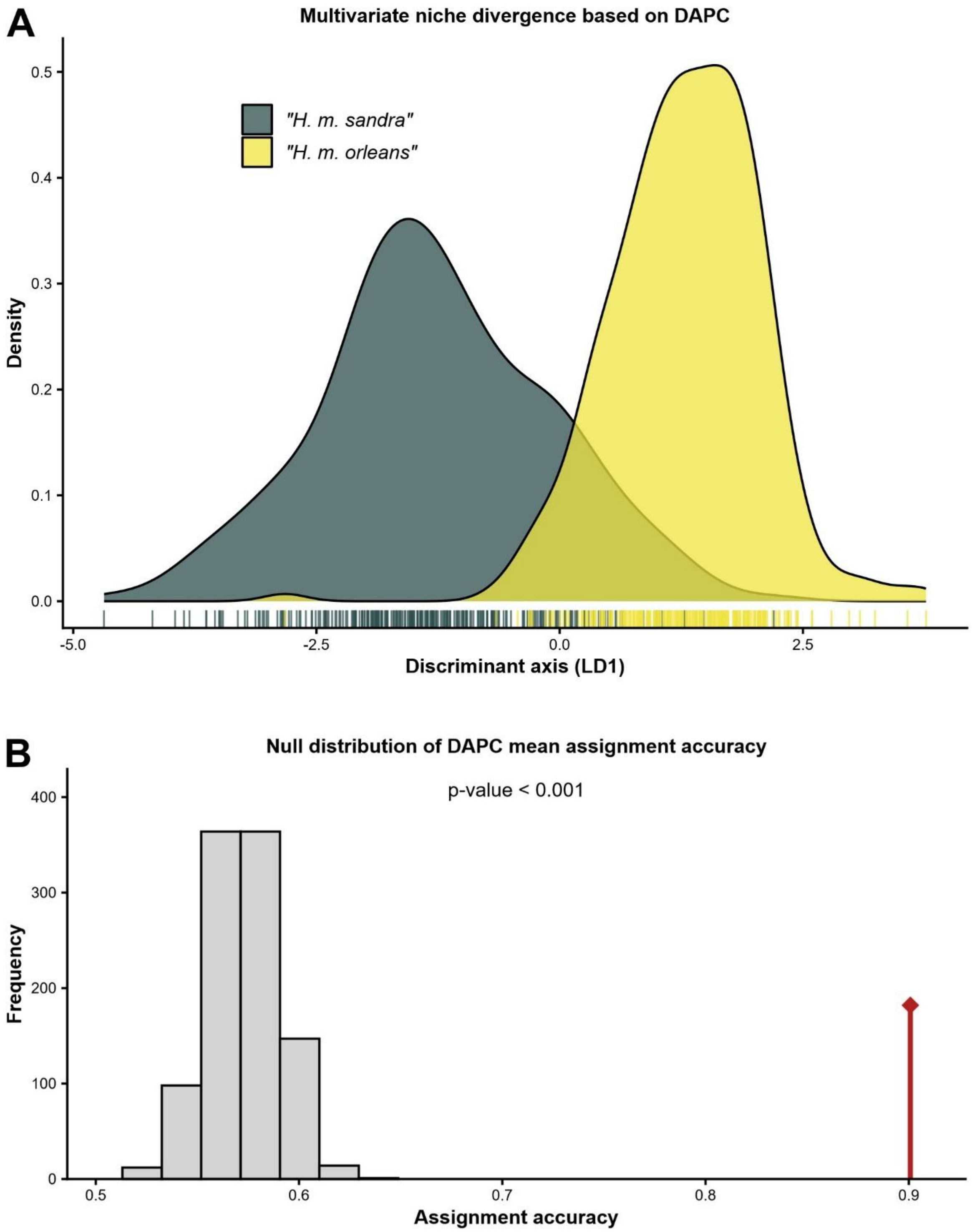


**Supplementary Figure S18: Discriminant analysis of principal components (DAPC) approach revealed moderate, significant multivariate niche divergence between "*Hemileuca maia sandra*" and "*H. m. orleans*" (empirical dataset 3). A)** Divergence along the discriminant axis (LD1). **B)** Null distribution of DAPC classification accuracy from permutations (grey bars) with observed value shown as red line.

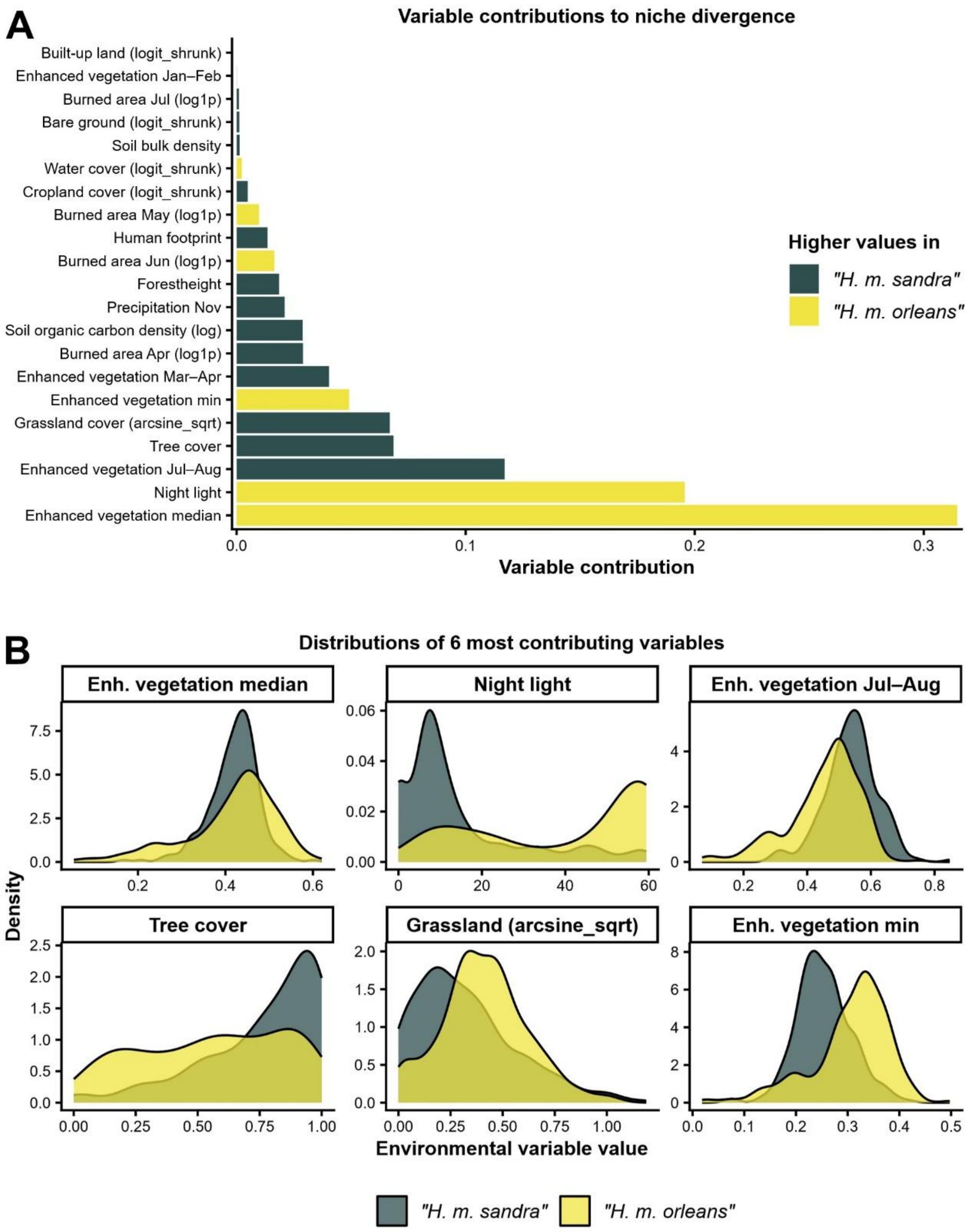


**Supplementary Figure S19: Contributions of environmental variables to multivariate niche divergence based on discriminant analysis of principal components (DAPC) niche divergence test between "*Hemileuca maia sandra*" and "*H. m. orleans*" (empirical dataset 3). A)** DAPC variable loadings to discriminant axis, with bar colors indicating the species with higher raw variable values. **B)** Kernel density plots of the raw distributions of the five top-contributing variables for both species. Months are abbreviated as three letter codes. Months are abbreviated as three letter codes. Data transformations are abbreviated in parentheses.

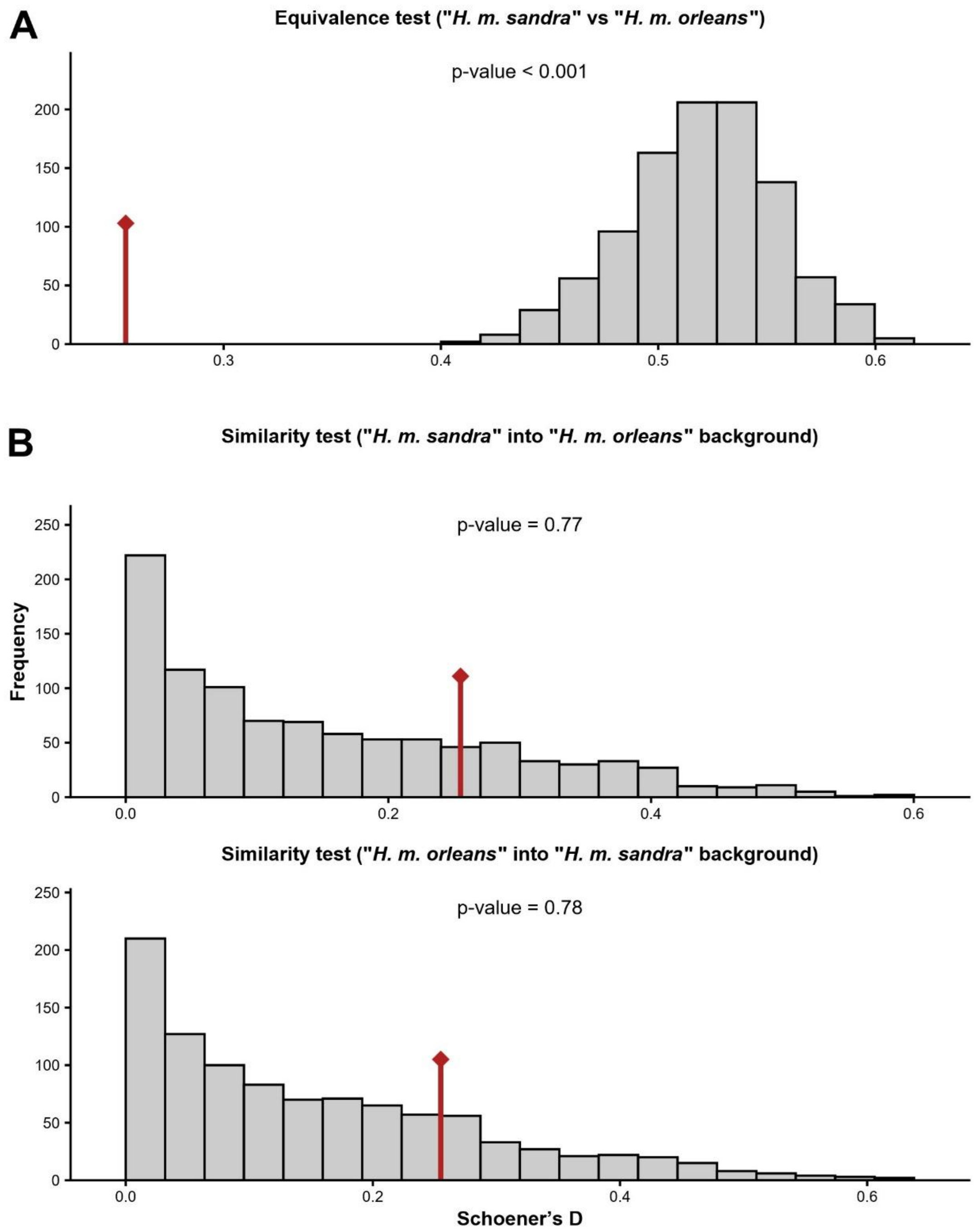


**Supplementary Figure S20: Permutation histograms for *ecospat* niche overlap tests between "*Hemileuca maia sandra*" and "*H. m. orleans*" (empirical dataset 3).** Grey bars show null distributions generated by randomizing species occurrences within background space and red lines mark observed values. **A)** Niche equivalence test: evaluates whether the environmental niches of "*H. m. sandra*" and "*H. m. orleans*" are statistically indistinguishable. **B)** Niche similarity test: examines whether the niche of "*H. m. sandra*" is more similar to the available background of "*H. m. orleans*" than expected by chance (upper row), and vice versa (lower row). Observed values in the null tails indicate rejection of equivalence or similarity, consistent with niche divergence or asymmetry.

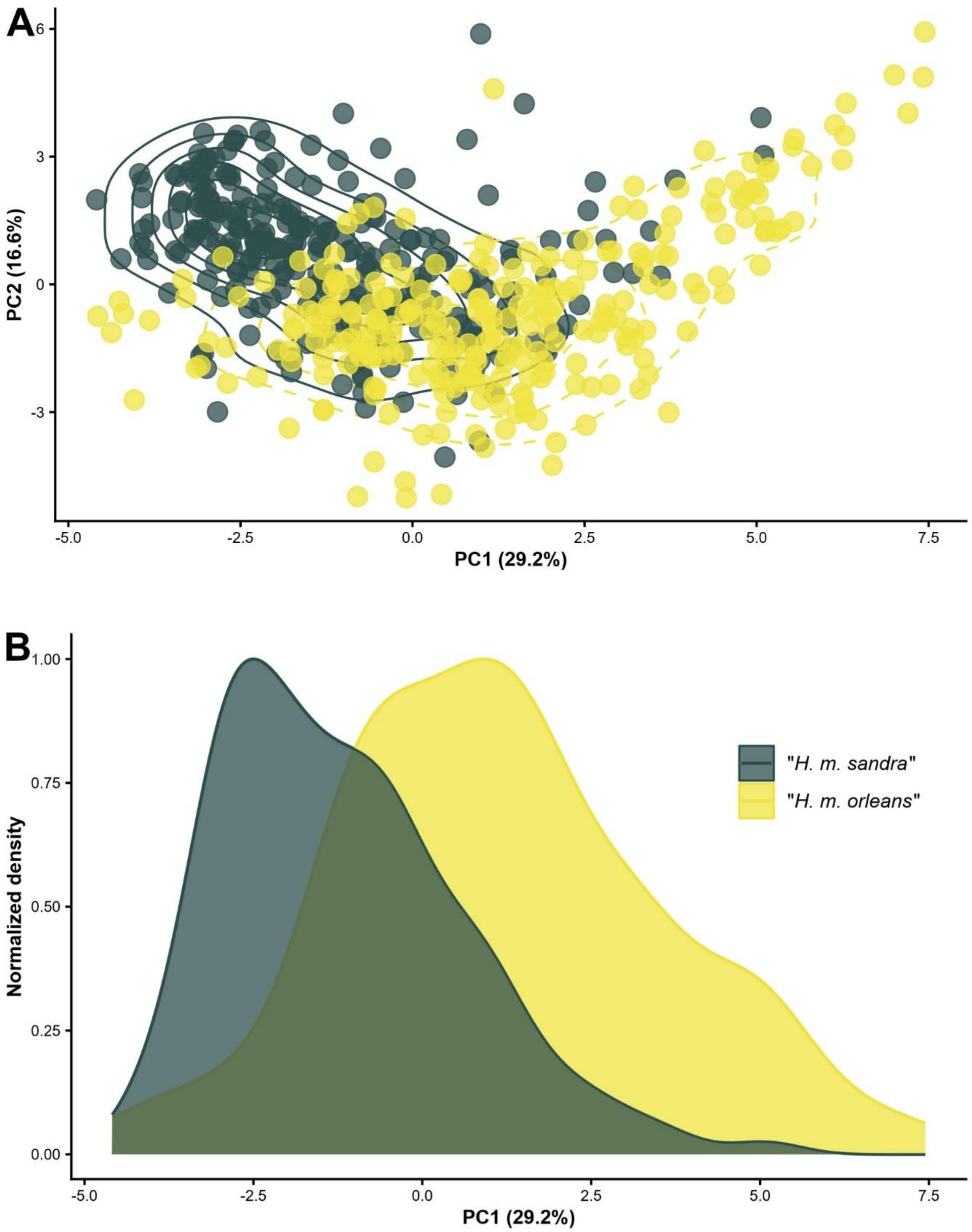


**Supplementary Figure S21: Niche divergence in analogous environmental principal component analysis space between "*H. maia sandra*" and "*H. m. orleans*" (empirical dataset 3). A)** Ordination of principal components (PC) 1 and 2 with occurrences plotted as points and occurrence densities (kernel-density isopleths) as contours. **B)** Density curves along PC1 (normalized to unit). Axis labels show the variance explained by each PC.

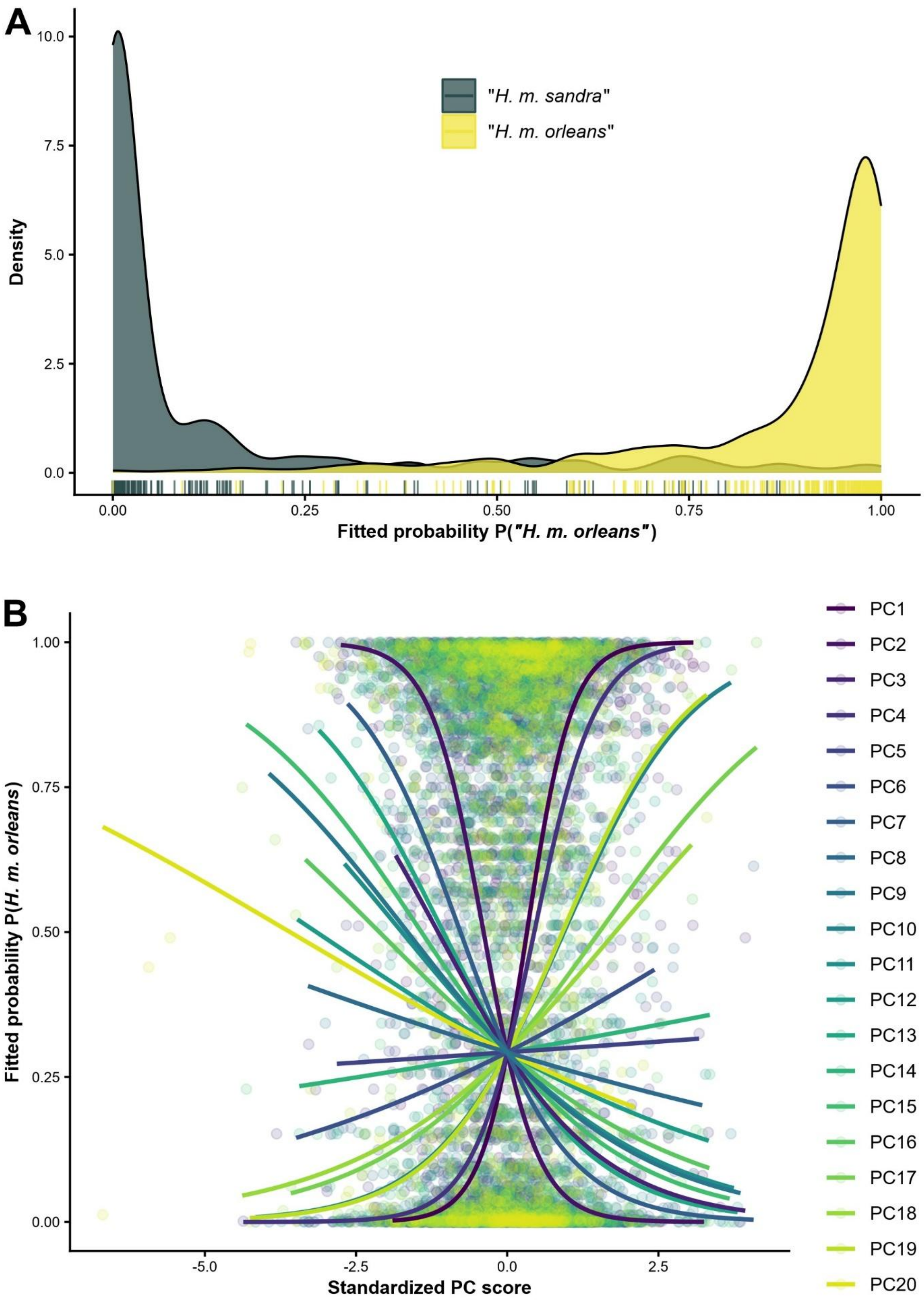
A
B
"H. m. sandra"
"H. m. orleans"
Density
10.0
7.5
5.0
2.5
0.0
0.00
0.25
0.50
0.75
1.00
Fitted probability P("H. m. orleans")
Fitted probability P(H. m. orleans)
Standardized PC score
-5.0
-2.5
0.0
2.5
PC1
PC2
PC3
PC4
PC5
PC6
PC7
PC8
PC9
PC10
PC11
PC12
PC13
PC14
PC15
PC16
PC17
PC18
PC19
PC20

**Supplementary Figure S22: Logistic regression classification based on binomial generalized linear model (logit link) of "*Hemileuca maia sandra*" vs. "*H. m. orleans*" (empirical dataset 3) using twenty principal components (PCs) of analogous environmental predictors. A)** X-axis shows fitted model probability *P* that a record belongs to "*H. m. orleans*", curves show kernel density estimates of fitted probabilities for each species, and rug ticks mark individual predictions. **B)** Partial dependence curves showing fitted probabilities based on model predictions along standardized PC scores, with points marking observed values. Steep curves indicate stronger niche divergence.

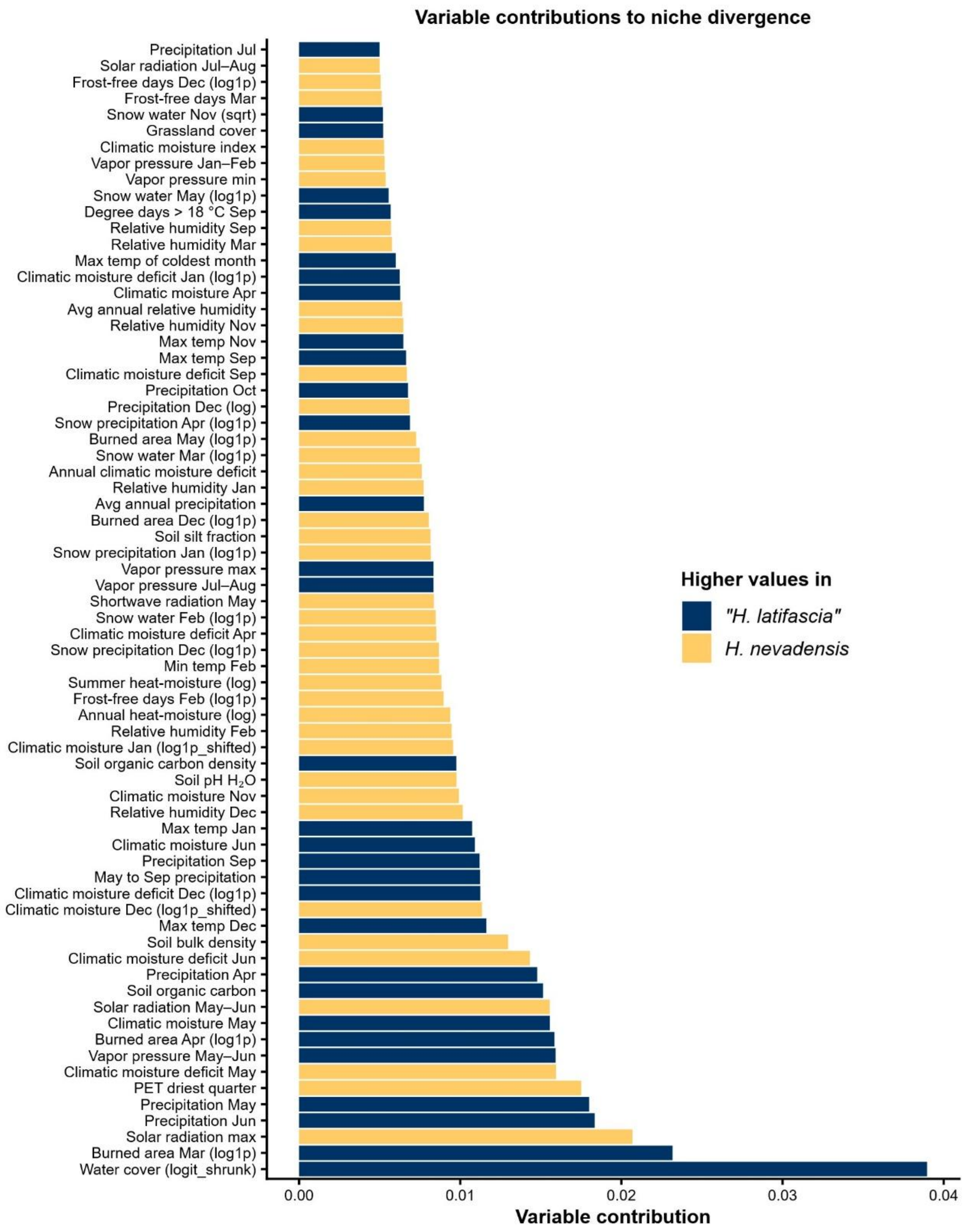


**Supplementary Figure S23: Contributions of environmental variables to multivariate niche divergence based on discriminant analysis of principal components (DAPC) niche divergence test between "*Hemileuca latifascia*" and *H. nevadensis* (empirical dataset 1) after analogous-environment filtering following Brown and Carnaval (2019).** Bars show variable loadings on the discriminant axis, with colors indicating which species exhibits higher raw environmental values. Only the 70 most important variables are displayed. Variable transformations are shown in parentheses, and months are abbreviated using three-letter codes.

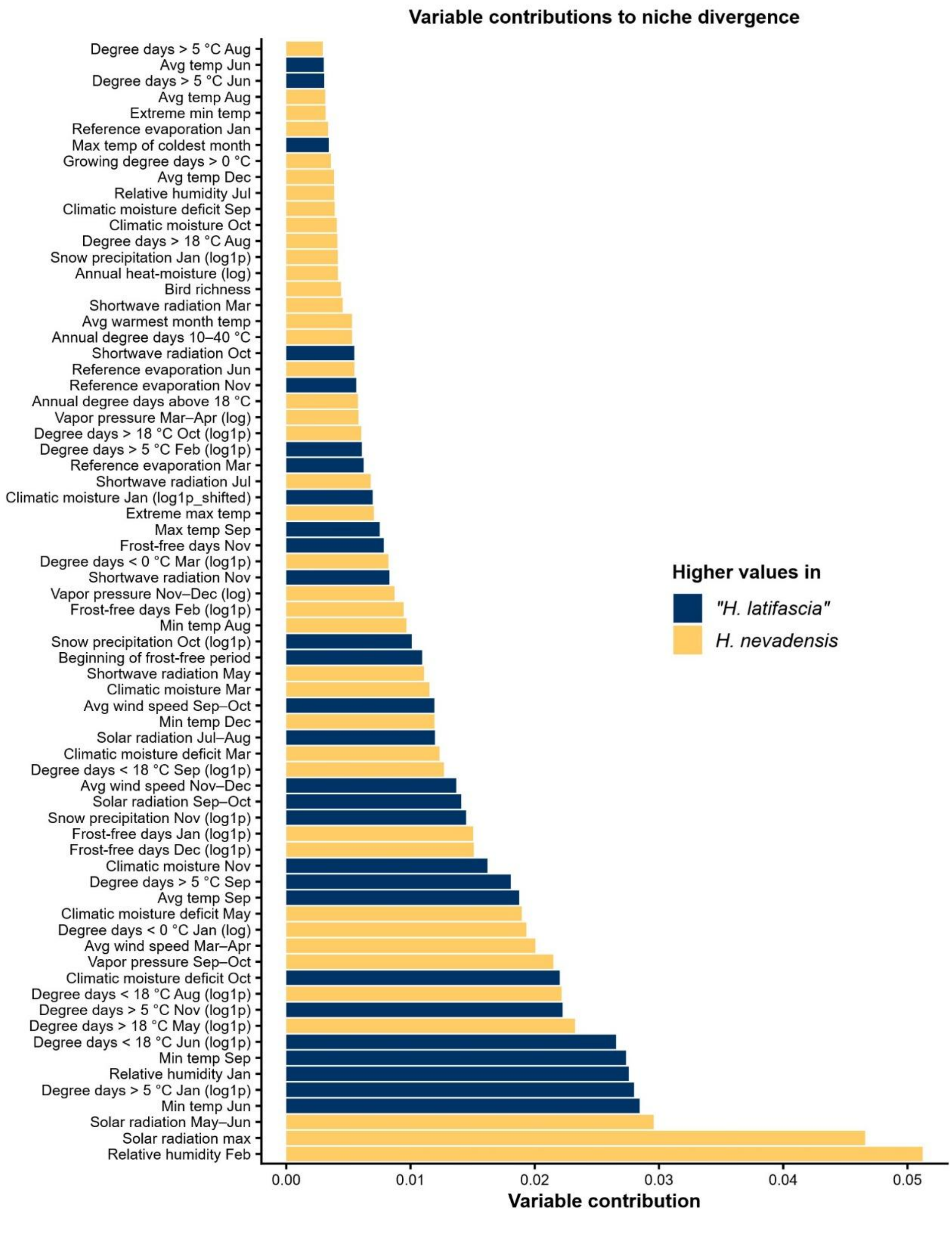


**Supplementary Figure S24: Contributions of environmental variables to multivariate niche divergence based on discriminant analysis of principal components (DAPC) niche divergence test between "*Hemileuca latifascia*" and *H. nevadensis* (empirical dataset 1) for full environmental dataset (analogous and non-analogous).** Bars show variable loadings on the discriminant axis, with colors indicating which species exhibits higher raw environmental values. Only the 70 most important variables are displayed. Variable transformations are shown in parentheses, and months are abbreviated using three-letter codes.

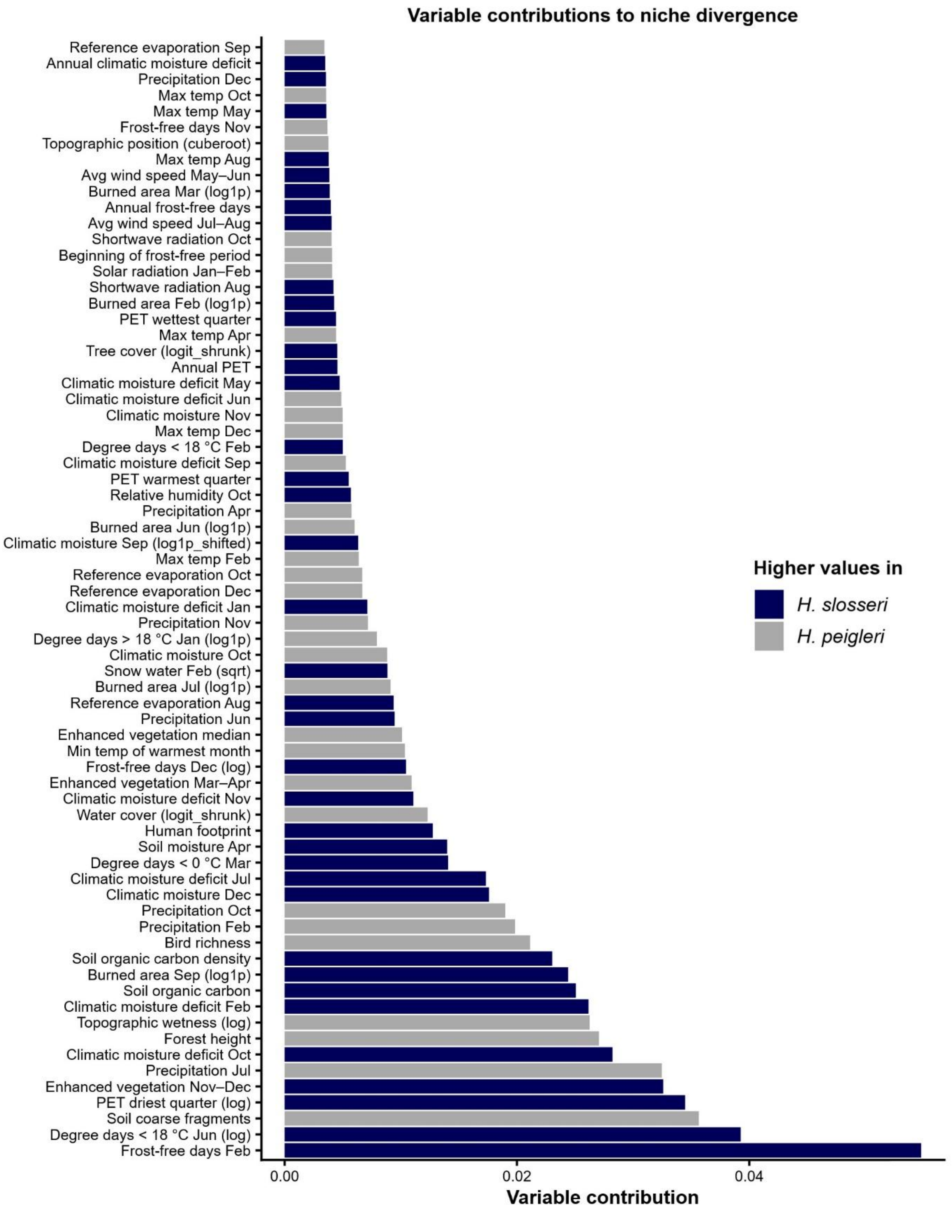


**Supplementary Figure S25: Contributions of environmental variables to multivariate niche divergence based on discriminant analysis of principal components (DAPC) niche divergence test between *Hemileuca slosseri* and *H. peigleri* (empirical dataset 2) for full environmental dataset (analogous and non-analogous).** Bars show variable loadings on the discriminant axis, with colors indicating which species exhibits higher raw environmental values. Only the 70 most important variables are displayed. Variable transformations are shown in parentheses, and months are abbreviated using three-letter codes.

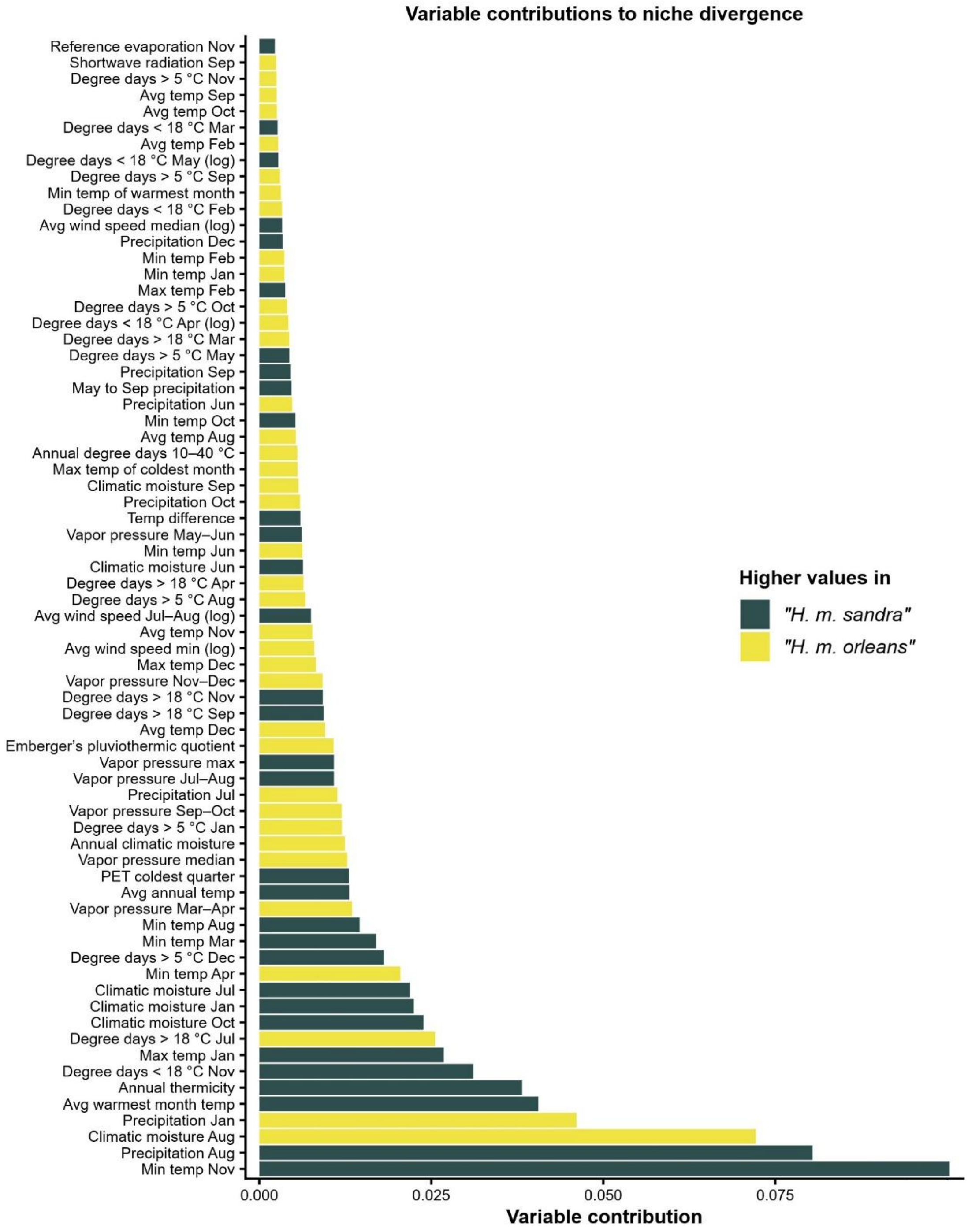


**Supplementary Figure S26: Contributions of environmental variables to multivariate niche divergence based on discriminant analysis of principal components (DAPC) niche divergence test between "*Hemileuca maia sandra*" and "*H. m. orleans*" (empirical dataset 3) for full environmental dataset (analogous and non-analogous).** Bars show variable loadings on the discriminant axis, with colors indicating which species exhibit higher raw environmental values. Only the 70 most important variables are displayed. Variable transformations are shown in parentheses, and months are abbreviated using three-letter codes.

**Supplementary Table S1:** Summary of functions in *NicheDiv R* package

| Function name | Description | Package dependencies |
|---|---|---|
| *calc.niche.divergence.metrics* | Computes niche divergence metrics (Schoener's D, NDS, NE, ND, θ) based on DAPC discriminant axis | *base R, base R stats* |
| *convert.integer.to.numeric* | Converts all integer columns in the occurrence and background data frame to numeric | *base R* |
| *crop.background.buffered* | Clips background points to buffered range polygons (hull/points/alpha/bbox) in a metric CRS | *base R stats, concaveman, sf* |
| *extract.env.and.background* | Extract comprehensive set of monthly, seasonal and annual environmental layers and create background points from occurrence data (coordinates) | *base R graphics, base R grDevices, base R methods, base R stats, base R utils, ClimateNAr, data.table, dplyr, geodata, sf, terra, tools, whitebox* |
| *filter.analogous.variables* | Keeps environmental variables analogous between taxa via CV, 1D KDE overlap, and fast 2D bin overlap | *base R graphics, base R stats, base R utils, dplyr, parallel, sf* |
| *transform.skewed.variables* | Identifies and transforms skewed environmental variables | *base R stats, base R utils* |
| *plot.DAPC.niche.divergence* | Plots DAPC results (discriminant axis) of niche divergence as density plots | *base R stats, base R utils, ggplot2, grid,* |
| *plot.DAPC.permutation* | Plots permutation null histograms vs observed assignment accuracy | *base R stats, base R utils, ggplot2, grid* |
| *plot.DAPC.var.contributions* | Bar plot of variable contributions to DAPC discriminant axis | *base R, base R grid, base R utils, ggplot2* |
| *plot.top.DAPC.predictors* | Density plots of raw distributions for strongest predictors by species | *base R, base R grid, ggplot2, stats, tidyr, tidyselect* |
| *plot.occurrences.map* | Plot occurrence and background points on political map | *base R graphics, base R grDevices, base R stats, maps, sf* |
| *remove.low.CV.vars* | Removes low-variation variables based on CV threshold from occurrence and background datasets | *base R, sf* |
| *run.DAPC.crossval.permutation* | Runs cross-validated DAPC with permutation test, reports performance, and computes ARI | *ade4, base R stats, base R utils, gstat, MASS, mclust, parallel, sf* |
| *sample.down* | Down-samples to N rows without replacement using Poisson-tail weights favoring fewer NAs (or uniform) | *base R, sf* |
| *thin.occurrence* | Spatially thins occurrences and checks residual autocorrelation (Moran's I) | *base R, fields, sf, spdep* |

*R* package versions: *ade4 v.1.7.23* (Bougeard & Dray, 2018; Chessel et al., 2004; Dray et al., 2007; Dray & Dufour, 2007; Thioulouse et al., 2018), *adegenet v.2.1.11* (Jombart, 2008, 2022; Jombart & Ahmed, 2011), *base R* (R Core Team, 2025), base R *graphics v.4.4.1* (R Core Team, 2025), *base R grDevices v.4.4.1* (R Core Team, 2025), *base R stats v.4.4.1* (R Core Team, 2025), *base R tools v.4.4.1* (R Core Team, 2025), *base R utils v.4.4.1* (R Core Team, 2025), *ClimateNAr v.3.1* (Wang et al., 2016), *concaveman v.1.1* (Gombin et al., 2020), *data.table v.1.17.8* (Barrett et al., 2006), *dplyr v.1.1.4* (Wickham et al., 2023), *fields v.16.3.1* (Nychka et al., 2021), *geodata v.0.6.6* (Hijmans et al., 2024), *ggplot2 v.4 (Wickham, 2016)*, *MASS v.7.3.65* (Venables & Ripley, 2002), *mclust v.6.1.1* (Scrucca et al., 2023), *parallel v.4.4.1* (R Core Team, 2025), *sf v.1.0.21* (Pebesma & Bivand, 2023), *spdep v.1.3.13* (Bivand, 2022; Bivand et al., 2013; Bivand & Wong, 2018; Pebesma & Bivand, 2023), *terra v.1.8.7* (Hijmans, 2025), *tidyr v.1.3.1* (Wickham et al., 2024), *whitebox v.2.4.3* (Lindsay, 2016; Wu & Brown, 2022). Abbreviations. 1D = univariate, 2D = bivariate, ARI = adjusted Rand index, CRS = coordinate reference system, CV = coefficient of variation, DAPC = discriminant analysis of principal components, KDE = kernel density estimation, NAs = missing values, NE = niche breadth exclusivity, ND = niche divergence magnitude, NDS = niche dissimilarity, θ = niche divergence angle.

**Supplementary Table S2:** Results of DAPC and six alternative niche divergence tests for "*Hemileuca latifascia*" and *H. nevadensis* (empirical dataset 1)

| Divergence test | Result metrics |
|---|---|
| DAPC | Mean assignment accuracy = 0.99, Adjusted Rand Index = 0.98, accuracy permutation test: p-value = 0.001 and mean = 0.62, Schoener's D overlap = 0.01, Schoener's D overlap after Humboldt-like background weighting = 0.02, niche dissimilarity (NDS) = 0.99, niche breadth exclusivity (NE) = 0.65, niche divergence magnitude (ND) = 1.18, ND after Humboldt-like background weighting = 1.15, niche divergence angle (θ) = 56.3° |
| env-PCA (ecospat) | Schoener's D = 0.29, niche equivalency test: p-value for D = 0.001, niche similarity tests: p-value for species "*H. latifascia*" vs. *H. nevadensis* background = 0.001, p-value for *H. nevadensis* vs. "*H. latifascia*" background = 1.00 |
| PERMANOVA | Permutational analysis of multivariate dispersions (test for homogeneity of dispersion): F = 8.07 and p-value = 0.001, PERMANOVA: F = 16.72 and p-value = 0.74 and R2 = 4.79% |
| hypervolume | Observed Jaccard similarity index = 0.37, observed Sørensen-Dice similarity index = 0.54, 42.8% of combined niche is unique to "*H. latifascia*", 49.0% of combined niche is unique to *H. nevadensis*, Euclidean centroid distance = 1.84, minimum inter-hypervolume distance = 0.11 |
| MVNH | Niche size (log-det) of "*H. latifascia*" = –2.10, Niche size (log-det) of *H. nevadensis* = 2.37, absolute niche size difference: 98.9%, BD (Bhattacharyya Dissimilarity - overall dissimilarity) = 2.28, MD (Mahalanobis Distance - centroid shift) = 0.58, DR (Determinant Ratio - breadth difference) = 1.70, MD/DR ratio: 0.34 |
| PCA space | Schoener's D (PC1–PC2 KDE) = 0.61, Ascanio metrics along PC1: niche dissimilarity (NDS) = 0.12, niche breadth exclusivity (NE) = 0.00, niche divergence magnitude (ND) = 0.12, niche divergence angle (θ) = 90.0° |
| Logistic regression | AIC null model = 465.0, AIC full model = 238.1, likelihood-ratio-test p-value < 0.0001, effect size (McFadden pseudo-$R^2$) = 0.56 |

See Supplementary Methods M6 for full descriptions of each niche divergence test. Abbreviations: DAPC = discriminant analysis of principal components, MVNH = multivariate normal hypervolume, PCA = principal component analysis, PERMANOVA = permutational multivariate analysis of variance.

**Supplementary Table S3:** Results of DAPC divergence tests across empirical dataset 1–3 using no analogous restrictions (full environmental dataset)

| Divergence test | Result metrics |
|---|---|
| Empirical dataset 1 | 167 samples for each group, 295 variables, optimal number of PCs retained for DAPC = 161, cumulative variance explained by retained PCs = 100%, mean assignment accuracy = 1.00, Adjusted Rand Index = 1.00, accuracy permutation test: p-value = 0.001 and mean = 0.83, Schoener's D overlap = 0.00, Schoener's D overlap after Humboldt-like background weighting = 0.00, niche dissimilarity (NDS) = 1.00, niche breadth exclusivity (NE) = 1.00, niche divergence magnitude (ND) = 1.41, ND after Humboldt-like background weighting = 1.41, niche divergence angle (θ) = 45.0° |
| Empirical dataset 2 | 39 samples for each group, 264 variables, optimal number of PCs retained for DAPC = 53, cumulative variance explained by retained PCs = 99.8%, mean assignment accuracy = 1.00, Adjusted Rand Index = 1.00, accuracy permutation test: p-value = 0.001 and mean = 0.93, Schoener's D overlap = 0.00, Schoener's D overlap after Humboldt-like background weighting = 0.00, niche dissimilarity (NDS) = 1.00, niche breadth exclusivity (NE) = 0.88, niche divergence magnitude (ND) = 1.33, ND after Humboldt-like background weighting = 1.41, niche divergence angle (θ) = 45.0° |
| Empirical dataset 3 | 263 samples for each group, 239 variables, optimal number of PCs retained for DAPC = 206, cumulative variance explained by retained PCs = 100%, mean assignment accuracy = 1.00, Adjusted Rand Index = 1.00, accuracy permutation test: p-value = 0.001 and mean = 0.79, Schoener's D overlap = 0.00, Schoener's D overlap after Humboldt-like background weighting = 0.02, niche dissimilarity (NDS) = 1.00, niche breadth exclusivity (NE) = 1.00, niche divergence magnitude (ND) = 1.41, ND after Humboldt-like background weighting = 1.41, niche divergence angle (θ) = 45.0° |

Empirical dataset 1 taxon pair: "*Hemileuca latifascia*" vs *H. nevadensis*, empirical dataset 2 taxon pair: *H. slosseri* vs *H. peigleri*, empirical dataset 3 taxon pair: "*H. maia sandra*" vs "*H. m. orleans*". Abbreviations: DAPC = discriminant analysis of principal components.

**Supplementary Table S4:** Results of DAPC divergence tests across empirical dataset 1–3 using the Brown and Carnaval (2019) analogous filtering of samples

| Divergence test | Result metrics |
|---|---|
| Empirical dataset 1 | 40 samples for each group, 295 variables, optimal number of PCs retained for DAPC = 14, cumulative variance explained by retained PCs = 92.3%, mean assignment accuracy = 1.00, Adjusted Rand Index = 1.00, accuracy permutation test: p-value = 0.001 and mean = 0.67, Schoener's D overlap = 0.00, Schoener's D overlap after Humboldt-like background weighting = 0.04, niche dissimilarity (NDS) = 1.00, niche breadth exclusivity (NE) = 0.79, niche divergence magnitude (ND) = 1.27, ND after Humboldt-like background weighting = 1.26, niche divergence angle (θ) = 51.8° |
| Empirical dataset 2 | No analogous environment remaining after trimming |
| Empirical dataset 3 | Only two samples left in analogous environment remaining after trimming |

Empirical dataset 1 taxon pair: “*Hemileuca latifascia*” vs *H. nevadensis*, empirical dataset 2 taxon pair: *H. slosseri* vs *H. peigleri*, empirical dataset 3 taxon pair: “*H. maia sandra*” vs “*H. m. orleans*”. Abbreviations: DAPC = discriminant analysis of principal components.

**Supplementary Table S5:** Results of DAPC and six alternative niche divergence tests for *H. slosseri* and *H. peigleri* (empirical dataset 2)

| **Divergence test** | **Result metrics** |
|---|---|
| DAPC | Mean assignment accuracy = 0.83, Adjusted Rand Index = 0.44, accuracy permutation test: p-value = 0.003 and mean = 0.70, Schoener's D overlap = 0.44, Schoener's D overlap after Humboldt-like background weighting = 0.39, niche dissimilarity (NDS) = 0.56, niche breadth exclusivity (NE) = 0.00, niche divergence magnitude (ND) = 0.56, ND after Humboldt-like background weighting = 0.71, niche divergence angle (θ) = 90.0° |
| env-PCA (ecospat) | Schoener's D = 0.33, niche equivalency test: p-value for D = 0.29, niche similarity tests: p-value for species *H. slosseri* vs. *H. peigleri* background = 0.78, p-value for *H. peigleri* vs. *H. slosseri* background = 0.77 |
| PERMANOVA | Permutational analysis of multivariate dispersions (test for homogeneity of dispersion): F = 0.00 and p-value = 1.00, PERMANOVA: F = 3.21 and p-value = 0.14 and R2 = 4.05% |
| hypervolume | Observed Jaccard similarity index = 0.32, observed Sørensen-Dice similarity index = 0.48, 50.3% of combined niche is unique to *H. slosseri*, 53.0% of combined niche is unique to *H. peigleri*, Euclidean centroid distance = 1.51, minimum inter-hypervolume distance = 0.12 |
| MVNH | Niche size (log-det) of *H. slosseri* = –1.72, Niche size (log-det) of *H. peigleri* = –1.03, absolute niche size difference: 49.7%, BD (Bhattacharyya Dissimilarity - overall dissimilarity) = 1.68, MD (Mahalanobis Distance - centroid shift) = 0.34, DR (Determinant Ratio - breadth difference) = 1.34, MD/DR ratio: 0.26 |
| PCA space | Schoener's D (PC1–PC2 KDE) = 0.81, Ascanio metrics along PC1: niche dissimilarity (NDS) = 0.12, niche breadth exclusivity (NE) = 0.00, niche divergence magnitude (ND) = 0.12, niche divergence angle (θ) = 90.0° |
| Logistic regression | AIC null model = 110.1, AIC full model = 85.9, likelihood-ratio-test p-value < 0.0001, effect size (McFadden pseudo-$R^2$) = 0.28 |

See Supplementary Methods M6 for full descriptions of each niche divergence test. Abbreviations: DAPC = discriminant analysis of principal components, MVNH = multivariate normal hypervolume, PCA = principal component analysis, PERMANOVA = permutational multivariate analysis of variance.

**Supplementary Table S6:** Results of DAPC and six alternative niche divergence tests for "*H. maia sandra*" and "*H. m. orleans*" (empirical dataset 3)

| Divergence test | Result metrics |
|---|---|
| DAPC | Mean assignment accuracy = 0.90, Adjusted Rand Index = 0.64, accuracy permutation test: p-value = 0.001 and mean = 0.57, Schoener's D overlap = 0.29, Schoener's D overlap after Humboldt-like background weighting = 0.11, niche dissimilarity (NDS) = 0.78, niche breadth exclusivity (NE) = 0.08, niche divergence magnitude (ND) = 0.78, ND after Humboldt-like background weighting = 0.88, niche divergence angle (θ) = 84.0° |
| env-PCA (ecospat) | Schoener's D = 0.29, niche equivalency test: p-value for D = 0.001, niche similarity tests: p-value for species "*H. m. sandra*" vs. "*H. m. orleans*" background = 0.86, p-value for "*H. m. orleans*" vs. "*H. m. sandra*" background = 0.86 |
| PERMANOVA | Permutational analysis of multivariate dispersions (test for homogeneity of dispersion): F = 12.59 and p-value = 0.004, PERMANOVA: F = 128.76 and p-value = 0.001 and R2 = 19.43% |
| hypervolume | Observed Jaccard similarity index = 0.13, observed Sørensen-Dice similarity index = 0.24, 75.5% of combined niche is unique to "*H. m. sandra*", 77.2% of combined niche is unique to "*H. m. orleans*", Euclidean centroid distance = 3.41, minimum inter-hypervolume distance = 0.29 |
| MVNH | Niche size (log-det) of "*H. m. sandra*" = 1.80, Niche size (log-det) of "*H. m. orleans*" = –2.28, absolute niche size difference: 5793.8%, BD (Bhattacharyya Dissimilarity - overall dissimilarity) = 1.47, MD (Mahalanobis Distance - centroid shift) = 0.60, DR (Determinant Ratio - breadth difference) = 0.87, MD/DR ratio: 0.70 |
| PCA space | Schoener's D (PC1–PC2 KDE) = 0.48, Ascanio metrics along PC1: niche dissimilarity (NDS) = 0.42, niche breadth exclusivity (NE) = 0.00, niche divergence magnitude (ND) = 0.42, niche divergence angle (θ) = 90.0° |
| Logistic regression | AIC null model = 745.1, AIC full model = 283.7, likelihood-ratio-test p-value < 0.0001, effect size (McFadden pseudo-$R^2$) = 0.67 |

See Supplementary Methods M6 for full descriptions of each niche divergence test. Abbreviations: DAPC = discriminant analysis of principal components, MVNH = multivariate normal hypervolume, PCA = principal component analysis, PERMANOVA = permutational multivariate analysis of variance.

**Supplementary Table S7:** Number of retained principal component axes and total amount of explained variance of our DAPC niche divergence approach and seven alternative methods across empirical and simulated datasets

| Dataset | Method | Number of PCs | Explained variance (%) |
|---|---|---|---|
| **Simulation** | DAPC | 34.6 35.0 | 97.3 87.6 |
| | env-PCA (ecospat) | 2.0 2.0 | 44.3 65.9 |
| | PERMANOVA | 6.6 7.2 | 65.2 79.8 |
| | hypervolume | 6.8 7.0 | 69.1 79.6 |
| | MVNH | 17.4 23.0 | 90.6 90.2 |
| | PCA space (Schoeners D and Ascanio) | 2.0 2.0 | 43.8 58.4 |
| | Logistic regression | 49.7 134.0 | 100.0 99.0 |
| **Empirical** | DAPC | 30 17 19 | 99.9 98.9 99.8 |
| | env-PCA (ecospat) | 2 2 2 | 30.5 26.8 36.9 |
| | PERMANOVA | 4 18 2 | 49.8 100 45.8 |
| | hypervolume | 5 4 6 | 54.7 50.7 73.1 |
| | MVNH | 18 12 11 | 90.4 90.1 90.9 |
| | PCA space (Schoeners D and Ascanio) | 1–2 1–2 1–2 | 23.4–34.0 17.0–32.2 29.2–45.8 |
| | Logistic regression | 16 3 20 | 87.1 41.9 100 |

See Supplementary Methods M6 for full descriptions of each niche divergence test. Mean values are shown across two simulations sets with 100 simulated species each and three empirical datasets. Double or triple values represent values for both simulation pairs or the three empirical datasets. Abbreviations: DAPC = discriminant analysis of principal components, MVNH = multivariate normal hypervolume, PCA = principal component analysis, PERMANOVA = permutational multivariate analysis of variance.

**Supplementary Table S8:** Runtime of DAPC and six alternative niche divergence tests across empirical and simulated datasets

| Dataset | Method | Runtime (min) |
|---|---|---|
| **Simulation** | DAPC | 1.01 4.57 |
| | env-PCA (ecospat) | 12.67 11.57 |
| | PERMANOVA | 1.87 3.31 |
| | hypervolume | 3.79 5.14 |
| | MVNH | 0.00 0.01 |
| | PCA space (Schoeners D and Ascanio) | 0.00 0.00 |
| | Logistic regression | 0.00 0.01 |
| **Empirical** | DAPC | 0.32 0.17 0.22 |
| | env-PCA (ecospat) | 16.3 9.9 11.2 |
| | PERMANOVA | 0.03 0.03 0.07 |
| | hypervolume | 0.68 0.15 0.96 |
| | MVNH | 0.00 0.00 0.00 |
| | PCA space (Schoeners D and Ascanio) | 0.00 0.00 0.00 |
| | Logistic regression | 0.00 0.00 0.00 |

See Supplementary Methods M6 for full descriptions of each niche divergence test. Mean values are shown across two simulations sets with 100 simulated species each and 3 empirical datasets. Double or triple values represent values for both simulation pairs or the three empirical datasets. Abbreviations: DAPC = discriminant analysis of principal components, MVNH = multivariate normal hypervolume, PCA = principal component analysis, PERMANOVA = permutational multivariate analysis of variance.

**Supplementary Table S9:** Description of all *ClimateNA* variables (Wang et al., 2016)

| Variable | Yearly/Monthly | Description |
|---|---|---|
| MAT | Y | Mean annual temperature (°C) |
| MWMT | Y | Mean warmest month temperature (°C) |
| MCMT | Y | Mean coldest month temperature (°C) |
| TD | Y | Temperature difference MWMT – MCMT (°C) |
| MAP | Y | Mean annual precipitation (mm) |
| MSP | Y | May–Sept precipitation (mm) |
| AHM | Y | Annual heat–moisture index |
| SHM | Y | Summer heat–moisture index |
| bFFP | Y | Julian date of beginning of frost-free period |
| eFFP | Y | Julian date of end of frost-free period |
| FFP | Y | Frost-free period length (days) |
| CMD | Y | Climatic moisture deficit (mm) |
| CMI | Y | Climate moisture index |
| DD_0 | Y | Degree-days below 0 °C |
| DD5 | Y | Degree-days above 5 °C |
| DD_18 | Y | Degree-days below -18 °C |
| DD18 | Y | Degree-days above 18 °C |
| DD1040 | Y | Degree-days between 10–40 °C |
| EMT | Y | Extreme minimum temperature over 30 years (°C) |
| EXT | Y | Extreme maximum temperature over 30 years (°C) |
| Eref | Y | Hargreaves reference evapotranspiration (mm) |
| Rsds | Y | Mean annual solar radiation (MJ/m²/day) |
| NFFD | Y | Number of frost-free days |
| PAS | Y | Precipitation as snow (mm) |
| RH | Y | Mean annual relative humidity (%) |
| Tmax | M | Monthly maximum temperature (°C; Tmax01–Tmax12) |
| Tmin | M | Monthly minimum temperature (°C; Tmin01–Tmin12) |
| Tave | M | Monthly mean temperature (°C; Tave01–Tave12) |
| PPT | M | Monthly total precipitation (mm; PPT01–PPT12) |
| Rsds | M | Monthly solar radiation (MJ/m²/day; rsds01–rsds12) |
| DD_0 | M | Monthly degree-days below 0 °C (DD_0_01–DD_0_12) |
| DD5 | M | Monthly degree-days above 5 °C (DD5_01–DD5_12) |
| DD_18 | M | Monthly degree-days below -18 °C (DD_18_01–DD_18_12) |
| DD18 | M | Monthly degree-days above 18 °C (DD18_01–DD18_12) |
| NFFD | M | Monthly frost-free days (NFFD01–NFFD12) |
| PAS | M | Monthly precipitation as snow (mm; PAS01–PAS12) |
| Eref | M | Monthly Hargreaves reference evapotranspiration (mm; Eref01–Eref12) |
| CMD | M | Monthly climatic moisture deficit (mm; CMD01–CMD12) |
| RH | M | Monthly relative humidity (%) (RH01–RH12) |
| CMI | M | Monthly climate moisture index (CMI01–CMI12) |

Pearson, K. (1896). VII. Mathematical contributions to the theory of evolution.—III. Regression, heredity, and panmixia. *Philosophical Transactions of the Royal Society of London. Series A, Containing Papers of a Mathematical or Physical Character*, *187*, 253–318. https://doi.org/10.1098/rsta.1896.0007

Pebesma, E., & Bivand, R. (2023). *Spatial data science: With applications in R*. Chapman and Hall/CRC. https://doi.org/10.1201/9780429459016

Peduzzi, P., Concato, J., Kemper, E., Holford, T. R., & Feinstein, A. R. (1996). A simulation study of the number of events per variable in logistic regression analysis. *Journal of Clinical Epidemiology*, *49*(12), 1373–1379. https://doi.org/10.1016/S0895-4356(96)00236-3

Peigler, R. S., & Stone, S. E. (1989). Taxonomic and biological notes on the Hemileuca maia complex (Saturniidae) with description of a new species from Texas and New Mexico. *The Lepidopterological Society of Japan*, *40*(2), 149–166.

Peterson, A. T., Papeş, T., & Eaton, M. (2007). Transferability and model evaluation in ecological niche modeling: A comparison of GARP and Maxent. *Ecography*, *30*(4), 550–560. https://doi.org/10.1111/j.0906-7590.2007.05102.x

Peterson, A. T., Soberón, J., Pearson, R. G., Anderson, R. P., Martínez-Meyer, E., Nakamura, M., & Araújo, M. B. (2011). *Ecological niches and geographic distributions*. Princeton University Press. https://doi.org/10.1515/9781400840670

Phillips, S. J., Anderson, R. P., & Schapire, R. E. (2006). Maximum entropy modeling of species geographic distributions. *Ecological Modelling*, *190*(3–4), 231–259. https://doi.org/10.1016/j.ecolmodel.2005.03.026

Phillips, S. J., Dudík, M., Elith, J., Graham, C. H., Lehmann, A., Leathwick, J., & Ferrier, S. (2009). Sample selection bias and presence-only distribution models: implications for background and pseudo-absence data. *Ecological Applications*, *19*(1), 181–197. https://doi.org/10.1890/07-2153.1

Phipson, B., & Smyth, G. K. (2010). Permutation p-values should never be zero: Calculating exact p-values when permutations are randomly drawn. *Statistical Applications in Genetics and Molecular Biology*, *9*(1). https://doi.org/10.2202/1544-6115.1585

Poggio, L., de Sousa, L. M., Batjes, N. H., Heuvelink, G. B. M., Kempen, B., Ribeiro, E., & Rossiter, D. (2021). SoilGrids 2.0: producing soil information for the globe with quantified spatial uncertainty. *SOIL*, *7*(1), 217–240. https://doi.org/10.5194/soil-7-217-2021

Potapov, P., Li, X., Hernandez-Serna, A., Tyukavina, A., Hansen, M. C., Kommareddy, A., Pickens, A., Turubanova, S., Tang, H., Silva, C. E., Armston, J., Dubayah, R., Blair, J. B., & Hofton, M. (2021). Mapping global forest canopy height through integration of GEDI and Landsat data. *Remote Sensing of Environment*, *253*, 112165. https://doi.org/10.1016/j.rse.2020.112165

Preimesberger, W., Scanlon, T., Su, C.-H., Gruber, A., & Dorigo, W. (2021). Homogenization of structural breaks in the global ESA CCI Soil Moisture multisatellite climate data record. *IEEE Transactions on Geoscience and Remote Sensing*, *59*(4), 2845–2862. https://doi.org/10.1109/TGRS.2020.3012896

R Core Team. (2025). *R: A language and environment for statistical computing* (4.4.1). R Foundation for Statistical Computing. https://www.r-project.org/.

Rödder, D., & Engler, J. O. (2011). Quantitative metrics of overlaps in Grinnellian niches: Advances and possible drawbacks. *Global Ecology and Biogeography*, *20*(6), 915–927. https://doi.org/10.1111/j.1466-8238.2011.00659.x

Schmidt, C. (2022). Biogeography and taxonomy of the Hemileuca maia-nevadensis Stretch species complex (Lepidoptera: Saturniidae): Insights to the midcontinental populations. *BioRxiv Preprint*. https://doi.org/10.1101/2022.04.07.487535

Schoener, T. W. (1968). The anolis lizards of Bimini: Resource partitioning in a complex fauna. *Ecology*, *49*(4), 704–726. https://doi.org/10.2307/1935534

Scott, D. W. (2015). *Multivariate density estimation: Theory, practice, and visualization* (2nd ed.). Wiley.

Scrucca, L., Fraley, C., Murphy, T. B., & Adrian E., R. (2023). *Model-based clustering, classification, and density estimation using mclust in R*. Chapman and Hall/CRC. https://doi.org/10.1201/9781003277965

Silverman, B. W. (1986). *Density estimation for statistics and data analysis*. Chapman and Hall.

Simpson, G. L. (2025). *permute: Functions for generating restricted permutations of data* (0.9-8).

Snyder, J. P. (1987). *Map projections: A working manual (U.S. Geological Survey Professional Paper 1395)*. Government Printing Office.

Soberón, J. (2007). Grinnellian and Eltonian niches and geographic distributions of species. *Ecology Letters*, *10*(12), 1115–1123. https://doi.org/10.1111/j.1461-0248.2007.01107.x

Soberón, J., & Peterson, A. T. (2005). Interpretation of models of fundamental ecological niches and species' distributional areas. *Biodiversity Informatics*, *2*(0). https://doi.org/10.17161/bi.v2i0.4

Sokal, R. R., & Rohlf, F. J. (2012). *Biometry: The principles and practice of statistics in biological research* (4th ed.). W. H. Freeman.

Sørensen, R., Zinko, U., & Seibert, J. (2006). On the calculation of the topographic wetness index: Evaluation of different methods based on field observations. *Hydrology and Earth System Sciences*, *10*(1), 101–112. https://doi.org/10.5194/hess-10-101-2006

Sørensen, T. (1948). A method of establishing groups of equal amplitude in plant sociology based on similarity of species content and its application to analyses of the vegetation on Danish commons. *Biologiske Skrifter*, *5*, 1–34.

St Laurent, R. (2023). Family Saturniidae. In *Annotated taxonomic checklist of the Lepidoptera of North America, north of Mexico* (pp. 282–285). Wedge Entomological Research Foundation.

Stevens, V. M., Turlure, C., & Baguette, M. (2010). A meta-analysis of dispersal in butterflies. *Biological Reviews*, *85*(3), 625–642. https://doi.org/10.1111/j.1469-185X.2009.00119.x

Thioulouse, J., Dray, S., Dufour, A.-B., Siberchicot, A., Jombart, T., & Pavoine, S. (2018). *Multivariate analysis of ecological data with ade4*. Springer New York. https://doi.org/10.1007/978-1-4939-8850-1

Title, P. O., & Bemmels, J. B. (2018). ENVIREM: An expanded set of bioclimatic and topographic variables increases flexibility and improves performance of ecological niche modeling. *Ecography*, *41*(2), 291–307. https://doi.org/10.1111/ecog.02880

Tukey, J. W. (1977). *Exploratory data analysis*. Addison–Wesley.

Turek, M. E., Poggio, L., Batjes, N. H., Armindo, R. A., de Jong van Lier, Q., de Sousa, L., & Heuvelink, G. B. M. (2023). Global mapping of volumetric water retention at 100, 330 and 15 000 cm suction using the WoSIS database. *International Soil and Water Conservation Research*, *11*(2), 225–239. https://doi.org/10.1016/j.iswcr.2022.08.001

Tuskes, P. M., Tuttle, J. P., & Collins, M. M. (1996). *The wild silk moths of North America: A natural history of the Saturniidae of the United States and Canada*. Cornell University Press.

Vaidyanathan, R., Xie, Y., Allaire, J., Cheng, J., Sievert, C., & Russell, K. (2023). *htmlwidgets: HTML widgets for R* (1.6.4). https://CRAN.R-project.org/package=htmlwidgets

van Buuren, S. (2018). *Flexible imputation of missing data*. Chapman & Hall/CRC.

VanDerWal, J., Shoo, L. P., Graham, C., & Williams, S. E. (2009). Selecting pseudo-absence data for presence-only distribution modeling: How far should you stray from what you know? *Ecological Modelling*, *220*(4), 589–594. https://doi.org/10.1016/j.ecolmodel.2008.11.010

Veloz, S. D. (2009). Spatially autocorrelated sampling falsely inflates measures of accuracy for presence-only niche models. *Journal of Biogeography*, *36*(12), 2290–2299. https://doi.org/10.1111/j.1365-2699.2009.02174.x

Venables, W. N., & Ripley, B. D. (2002). *Modern applied statistics with S* (4th ed.). Springer.

Venter, O., Sanderson, E. W., Magrach, A., Allan, J. R., Beher, J., Jones, K. R., Possingham, H. P., Laurance, W. F., Wood, P., Fekete, B. M., Levy, M. A., & Watson, J. E. M. (2016). Sixteen years of change in the global terrestrial human footprint and implications for biodiversity conservation. *Nature Communications*, *7*(1), 12558. https://doi.org/10.1038/ncomms12558

Vittinghoff, E., & McCulloch, C. E. (2007). Relaxing the rule of ten events per variable in logistic and Cox regression. *American Journal of Epidemiology*, *165*(6), 710–718. https://doi.org/10.1093/aje/kwk052

Wagner, D. L. (2010). *Wagner, David L. Caterpillars of eastern North America. A guide to identification and natural history*. Princeton University Press. https://doi.org/10.1515/9781400834143.16b

Wagner, H. H., & Dray, S. (2015). Generating spatially constrained null models for irregularly spaced data using Moran spectral randomization methods. *Methods in Ecology and Evolution*, *6*(10), 1169–1178. https://doi.org/10.1111/2041-210X.12407

Wang, T., Hamann, A., Spittlehouse, D., & Carroll, C. (2016). Locally downscaled and spatially customizable climate data for historical and future periods for North America. *PLOS ONE*, *11*(6), e0156720. https://doi.org/10.1371/journal.pone.0156720

Warren, D. L., Glor, R. E., & Turelli, M. (2008). Environmental niche equivalency versus conservatism: Quantitative approaches to niche evolution. *Evolution*, *62*(11), 2868–2883. https://doi.org/10.1111/j.1558-5646.2008.00482.x

Warren, D. L., Glor, R. E., & Turelli, M. (2010). ENMTools: A toolbox for comparative studies of environmental niche models. *Ecography*, *33*(3), 607–611. https://doi.org/10.1111/j.1600-0587.2009.06142.x

Warton, D. I., & Hui, F. K. C. (2011). The arcsine is asinine: The analysis of proportions in ecology. *Ecology*, *92*(1), 3–10. https://doi.org/10.1890/10-0340.1

Weil. R., & Brady, N. (2016). *The nature and properties of soils* (15th ed.). Pearson.

Wickham, H. (2016). *ggplot2: Elegant graphics for data analysis*. Springer Verlag. https://ggplot2.tidyverse.org

Wickham, H., François, R., Henry, L., Müller, K., & Vaughan, D. (2023). *dplyr: A grammar of data manipulation* (1.1.4).

Wickham, H., Vaughan, D., & Girlich, M. (2024). *tidyr: Tidy messy data* (1.3.1).

Wilson, M. F. J., O'Connell, B., Brown, C., Guinan, J. C., & Grehan, A. J. (2007). Multiscale terrain analysis of multibeam bathymetry data for habitat mapping on the continental slope. *Marine Geodesy*, *30*(1–2), 3–35. https://doi.org/10.1080/01490410701295962

Wu, Q., & Brown, A. (2022). *whitebox: "WhiteboxTools" R frontend* (2.4.0). https://CRAN.R-project.org/package=whitebox

Zar, J. H. (2010). *Biostatistical analysis*. Pearson.

Zizka, A., Silvestro, D., Andermann, T., Azevedo, J., Duarte Ritter, C., Edler, D., Farooq, H., Herdean, A., Ariza, M., Scharn, R., Svantesson, S., Wengström, N., Zizka, V., & Antonelli, A. (2019). CoordinateCleaner: Standardized cleaning of occurrence records from biological collection databases. *Methods in Ecology and Evolution*, *10*(5), 744–751. https://doi.org/10.1111/2041-210X.13152

Zurell, D., Berger, U., Cabral, J. S., Jeltsch, F., Meynard, C. N., Münkemüller, T., Nehrbass, N., Pagel, J., Reineking, B., Schröder, B., & Grimm, V. (2010). The virtual ecologist approach: simulating data and observers. *Oikos*, *119*(4), 622–635. https://doi.org/10.1111/j.1600-0706.2009.18284.x
